# Overcoming Immune Checkpoint Inhibitor Resistance via Potent and Selective Dual αvβ6/8 Inhibitors Based on Engineered Lasso Peptides

**DOI:** 10.1101/2025.01.28.635346

**Authors:** Anna Lechner, Peter A. Jordan, Gabriella Costa Machado da Cruz, Jacob Lamson, Jessica Gordon, Bethany K. Okada, Kelsey Anderson, Rajan Chaudhari, Christopher J. Rosario, Jasmine Mikesell, Scott A. McPhee, Mark J. Burk

## Abstract

Integrins αvβ6 and αvβ8 in the tumor microenvironment (TME) have been shown to activate immunosuppressive TGF-β, which serves as an important mechanism for immune checkpoint inhibitor resistance in a range of tumors. In this study, we demonstrate the utility of lasso peptides as versatile scaffolds for designing new therapeutics. A series of highly potent and selective dual αvβ6/8 inhibitors were engineered through a combination of epitope scanning, computational design, and directed evolution. Several analogs, such as lassotides **36** and **47**, were fully characterized and physicochemical, in vitro pharmacological, and in vivo data are reported. Lassotide **47**, a half-life extended derivative of **36**, was shown to strongly sensitize anti-mPD-1-resistant tumors in mice when dosed in combination with the checkpoint inhibitor. The **47**/anti-mPD-1 combination was shown to halt tumor growth and regress tumors in mouse models of triple negative breast and ovarian cancers. Dual inhibition of αvβ6/8 integrins expressed in the TME thus represents a promising tumor-specific strategy to overcome TGF-β-driven resistance and enhance the anti-tumor efficacy of immune checkpoint inhibitors.

## Introduction

Integrins are a class of ubiquitous transmembrane cell-surface receptors that regulate cell adhesion, migration, proliferation, and survival in mammals.^1,2,3^ In humans, each integrin consists of an α-subunit and a β-subunit, of which there are 18 and 8 variants, respectively, creating 24 different heterodimeric receptors.^2^ Given the large number of diseases impacted by integrins, integrin inhibitors have been extensively explored for new therapeutic applications.^4,5^

An important integrin subclass consists of eight receptors that bind tightly to the arginine-glycine-aspartate (RGD) epitope present in their endogenous ligands, which include fibrinogen and extracellular matrix (ECM) proteins, such as fibronectin and vitronectin.^6^ Two members of the RGD-binding integrin subclass are αvβ6 and αvβ8, both of which have been shown to be significantly upregulated on tumors and play important roles in cancer progression, invasion, and metastasis.^7^ For example, integrin αvβ6 is overexpressed on a wide range of different solid tumors, including colorectal, gastric, pancreatic, ovarian, cervical, triple negative breast, non-small cell lung cancers, head and neck squamous cell and oral squamous carcinomas.^8^ In addition to enhancing migration and invasion of tumors, αvβ6 also is known as a primary activator of transforming growth factor beta (TGF-β), which is a cytokine that promotes cancer fibrosis, epithelial-to-mesenchymal transition, and angiogenesis and potently suppresses the anti-tumor immune response of CD8^+^ T cells and other immune cells in the tumor microenvironment (TME).^9,10,11^ Similarly, the integrin receptor αvβ8 has been shown to be abundantly overexpressed on various tumors such as melanoma, pancreatic, colon, prostate, ovarian, triple negative breast, and non-small cell lung cancers, as well as head and neck squamous cell and renal carcinomas^12,13,14^ and on activated tumor-associated CD4^+^ Tregs.^15,16^ As with αvβ6,^17^ integrin αvβ8 binds to the epitope RGDL(XXL) in the sequestering latency associated peptide (LAP)^18^ and this interaction liberates immunosuppressive TGF-β from its non-covalent latency complex in the TME.^19^ Importantly, TGF-β activation enhances immune evasion in tumors and serves as a primary source of tumor resistance to immune checkpoint inhibitors such as anti-PD-1 and anti-PD-L1 therapies.^20,21,22,23^ Systemic blockade of pleiotropic TGF-β has been shown to complement immune checkpoint inhibitors and enhance their anti-tumor activity, albeit with a range of treatment-associated toxicities such as skin lesions, and other undesired adverse events.^24,25^ Thus, reducing the activity of TGF-β through inhibition of activating αvβ6 and αvβ8 integrins overexpressed in the TME represents a compelling tumor-specific strategy for the development of new immuno-oncology agents for treating a broad range of cancers.^26,27^

Herein, we describe the design and optimization of new dual inhibitors targeting both αvβ6 and αvβ8 integrins. Our approach leverages a novel class of diverse naturally occurring scaffolds called lasso peptides.^28.29^ Lasso peptides are highly stable, compact, modular structures (15-25 amino acids) comprising a distinctive knotted lariat fold that engenders a unique 3D arrangement of side-chain functionality for engagement with complex biological receptors. Lassotides (non-natural lasso peptide variants) were rapidly engineered through epitope scanning, computational design, and evolution methods to afford analogs with high potency and selectivity for αvβ6 and αvβ8. Detailed in vitro and in vivo characterization demonstrated the drug-like properties and utility of these compounds, ultimately culminating in robust efficacy in anti-PD-1-resistant murine models of triple negative breast and ovarian cancers. These studies validate the great potential of lasso peptides as a rich, untapped source of novel therapeutics.

## Results

### Lassotides (engineered lasso peptides) and integrin inhibitor discovery

The unique structures of lasso peptides are formed by the coordinated action of two main enzymes, a lasso peptidase and a lasso cyclase, which collaboratively cleave a linear precursor peptide to release a leader sequence. The resulting core peptide subsequently is converted by the lasso cyclase into a threaded conformation through amide bond formation between the amino-terminus and the side-chain carboxyl group of an Asp or Glu residue at position 7, 8, or 9, whereby the incipient macrocycle traps the C-terminal tail inside the ring, thus creating the characteristic loop-ring-tail architecture (**Figure 1**).^28^

**Figure 1.**
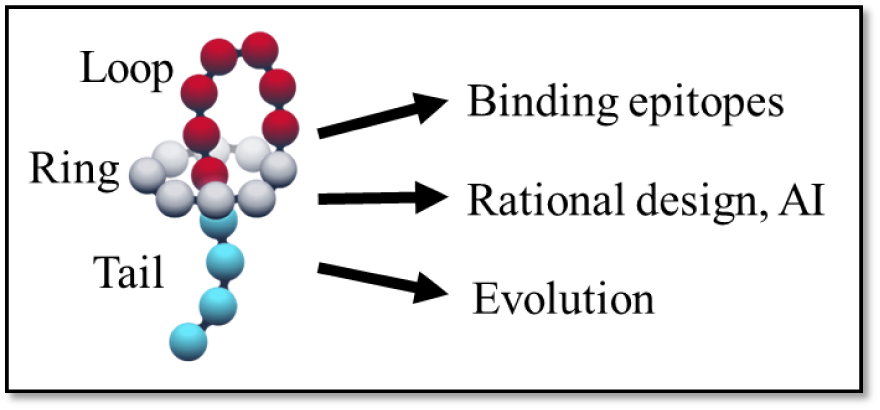
The general structure of lasso peptides showing the loop-ring-tail architecture and methods used to engineer desirable properties.

Lasso peptides of many shapes and sizes are produced by bacteria for modulating different biological targets and to date over 7000 curated lasso scaffold sequences have been predicted by the RODEO algorithm using GenBank bacterial whole genome sequence data.^30,31^ Thus, lasso peptides are constrained scaffolds that offer a diverse set of starting points for drug discovery, and which contain a hypervariable loop that mimics the CDR3H loops of antibodies, but in a much smaller format (2-3 kDa vs 150 kDa). The length of known lasso peptide loops spans 3-20 amino acids, which provides a wider range of loop sizes to engineer relative to antibodies. Like other polypeptides, lasso peptides are ribosomally produced and are amenable to sequence-based mutagenesis methods for the discovery and optimization of lassotide variants (engineered lasso peptides) that bind tightly to receptors of interest. Importantly, the wild-type enzymes used in the biosynthetic process can limit the diversity of lassotides that can be produced through evolution, and we recently demonstrated the benefits of lasso cyclase engineering for expanding mutagenic tolerance.^32^

As described below, we initiated our efforts to discover potent and selective αvβ6/8 dual integrin inhibitors by introducing the RGDX integrin-binding epitope (X = any natural or unnatural amino acid) into selected lasso peptide scaffolds to assess integrin binding and inhibition.

### Epitope scanning

The modular loop-ring-tail structures of lasso peptides are ideally suited for grafting of known receptor binding epitopes derived from endogenous, antibody, or peptide ligands shown to engage a receptor of interest. This unique framework affords the opportunity to rapidly survey the impact of epitope sequence and position across the lasso backbone. Previous studies by Marahiel and co-workers demonstrated that the integrin binding epitope RGD could be introduced by substituting the GIG residues at positions 12-14 in the loop of natural lasso peptide MccJ25.^33,34^ This RGD grafted derivative, and an RGDF variant, were found to inhibit the integrin αvβ3 with high potency (IC50 = 12 and 4 nM, respectively). However, no binding data for αvβ6 and αvβ8 was reported.

Based on these results, we initially examined a range of RGDX epitopes in the loop of several wild type scaffolds, including MccJ25,^35^ klebsidin,^36^ and caulonodin V.^37^ The results of incorporating RGDX into different scaffold loops, as well as the impact of grafting RGD into the ring, changing the X residue, and varying the loop position, are shown in **Table 1**.

**Table 1.**
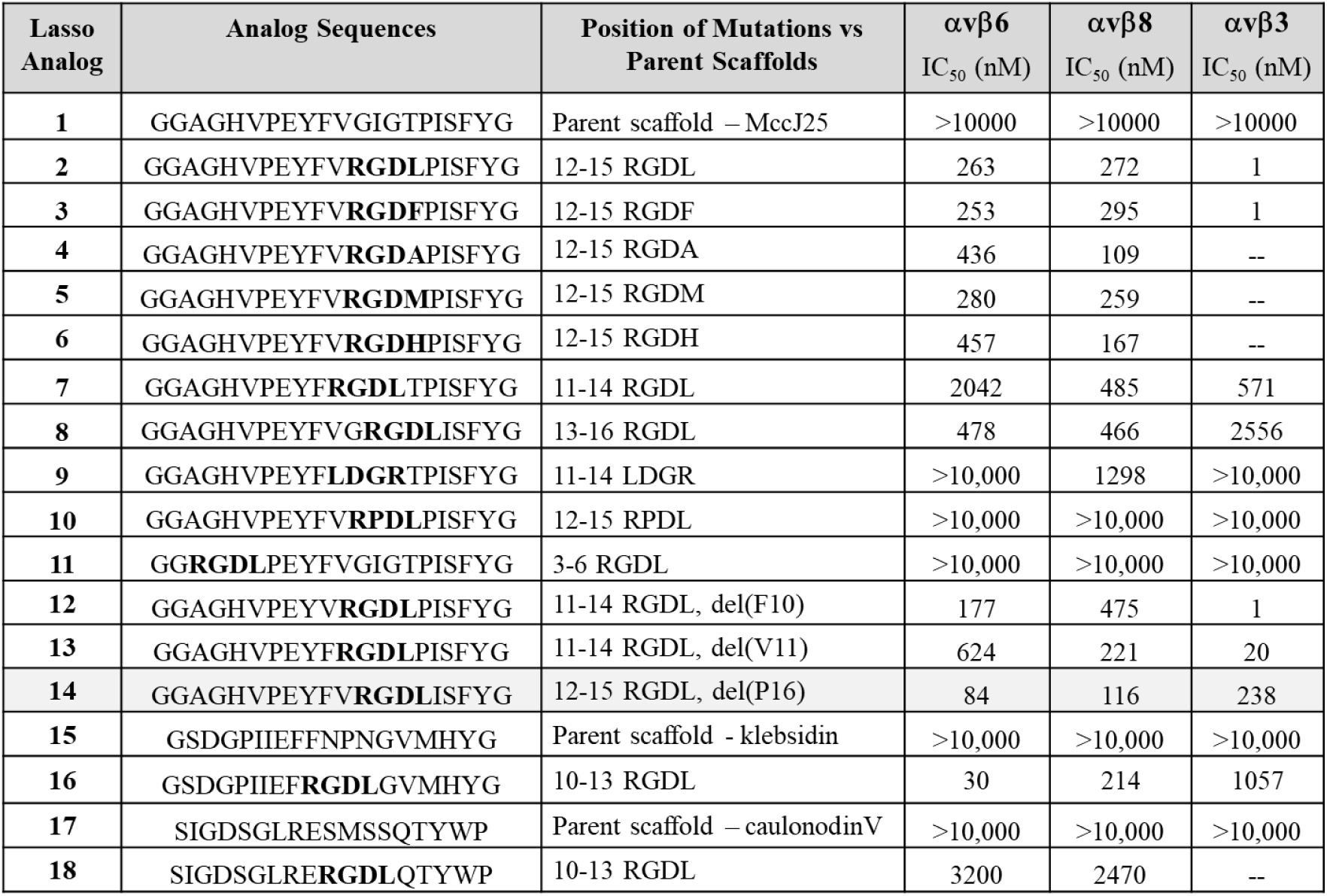
Integrin inhibition data for engineered lasso peptides bearing an RGDX epitope. Analog sequences are provided along with the position of mutations introduced relative to the parent scaffold. Lassotide inhibition potency was determined in ELISA or AlphaLISA competition assays vs latency-associated peptide (LAP; UniProt accession no. P01137.2; R&D Systems, cat. no. 246-LP). For clarity, inhibition data for the three integrins, αvβ3, αvβ6, and αvβ8, is presented as mean values determined from a minimum of 3 experiments (n ≥ 3). See Experimental Methods for details.

Integrins αvβ6 and αvβ8 have been shown to preferentially bind the RGDL epitope in the LAP-TGF-β latency complex.^17,38^ Grafting RGDL into the loop of wild type MccJ25 at positions 12-15 led to potent inhibition of αvβ3 (lassotide **2**), consistent with the results of Hegemann et al.,^34^ but only modest inhibition of αvβ6 and αvβ8 was observed. Substituting L with other residues at the X position (**3**-**6**) did not improve inhibitor potency. Changing the position of RGDL in the loop (**7**,**8**), reversing the sequence order to LDGR (**9**), and substituting P for G in the epitope (**10**) lowered potency, even for αvβ3. Replacing the 3-6 ring residues of MccJ25 with RGDL afforded **11** with no detectable integrin binding (IC50 >10 μM), likely due to repulsive interactions between the lasso tail and the integrin. Deletions of F10 or V11 in lassotide **2** yielded analogs **12** and **13** which retained potent αvβ3 inhibition. The first improvement in αvβ6/8 potency and selectivity was observed when P16 of lassotide **2** was deleted, affording lassotide **14** which displayed 2-3-fold selectivity favoring αvβ6/8 over αvβ3 and IC50 levels below 100 nM. Computational modeling suggested that the P16 deletion may position the RGDL epitope in a proper orientation for more selective αvβ6/8 engagement. Placing RGDL in the loop of the natural lasso peptide klebsidin led to promising inhibition and selectivity for αvβ6 but attempts to introduce further modifications into lassotide **16** were not successful. Engineering of klebsiden cyclase is likely required to expand the accessible range of analogs. The RGDL epitope in the loop of caulonodin V resulted in only modest inhibition.

From this initial series, lassotide **14** with ΔP16 afforded the best dual αvβ6/8 inhibition and selectivity vs αvβ3 and was produced at >25 mg/L titer in shake flasks using wild-type cyclase (see below). Hence, lassotide **14** was selected as the parent scaffold for further modifications.

### Computational design and evolution

Lassotide analog **14** contains a single loop deletion (ΔP16) relative to lassotide **2** and an RGDL epitope at positions 12-15 of the ten-residue loop that spans positions 9-18. To gain a sense for lassotide-integrin binding and design potential, the MOE molecular modeling system (Chemical Computing Group) was implemented to dock the modeled structure of lasso **14** into the known structures of αvβ6 (PDB: 4UM8)^17^ and αvβ8 (PBD: 6OM1),^38^ and Arg and Asp of the RGD epitope were locked into their proper binding interactions with αv-subunit Asp-218 and the MIDAS Mg^2+^ ion, respectively. Our models indicated that the conformation of the loop in lassotide **14**, and hence the binding orientation of the RGDL epitope, would be impacted by neighboring loop and proximal ring residues. Therefore, initial site-directed mutagenesis focused on varying residues adjoining the RGDL epitope, as well as positions in the ring that could interact with the integrin receptor and possibly influence the loop conformation. **Table 2** shows that changing the epitope-flanking aliphatic residues V11 and I16 to more polar amino acids R or Q led to significant improvements, with I16Q (**24**) showing up to 7-fold increase in potency for inhibition of both αvβ6 and αvβ8, relative to starting lassotide **14**. Similarly, increasing the size and polarity/charge of the A3 ring residue (e.g., A3R, **30**) afforded 3-4-fold improvement in IC50 values.

**Table 2.**
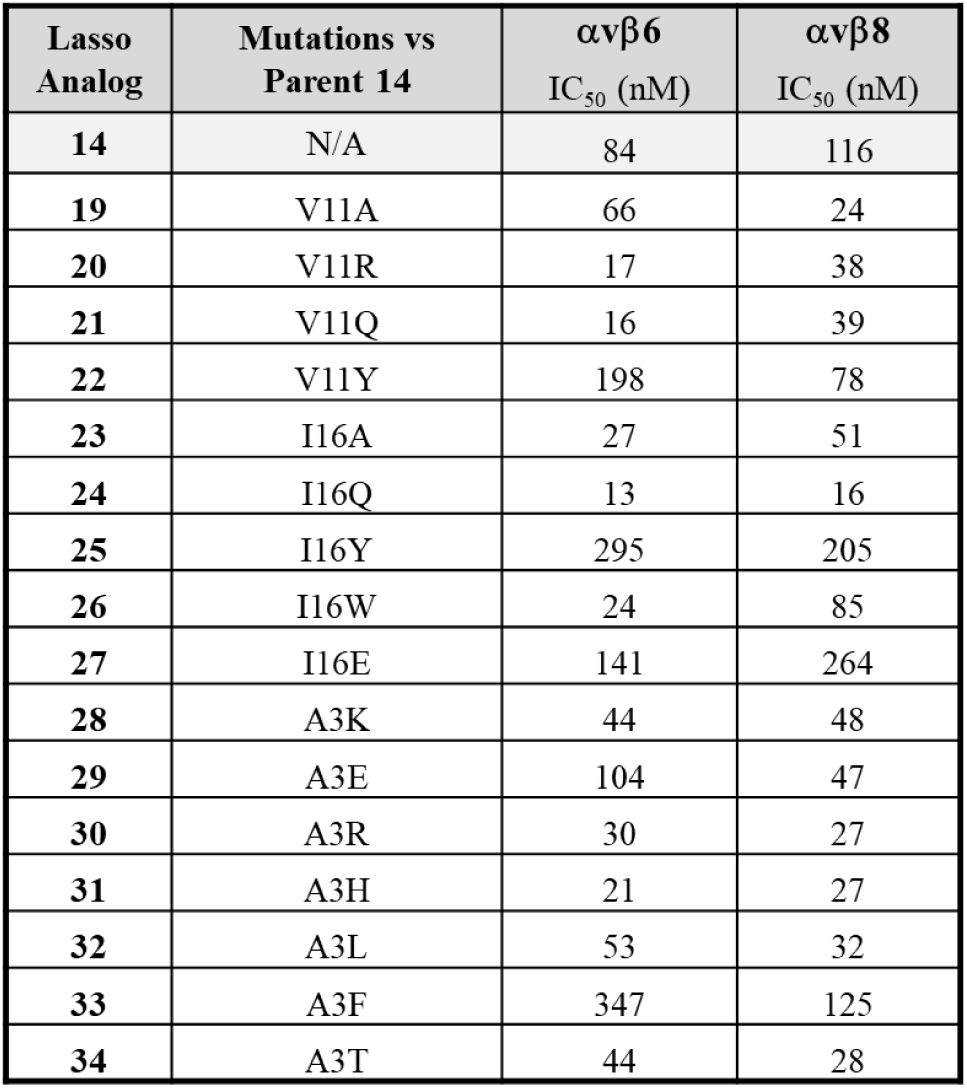
Integrin inhibition data for engineered lasso peptides derived from lassotide **14** with mutations introduced relative to this parent scaffold. Lassotide inhibition potency was determined in ELISA or AlphaLISA competition assays vs latency-associated peptide (LAP). For clarity, inhibition data for αvβ6 and αvβ8 is presented as mean values determined from a minimum of 3 experiments (n ≥ 3). See Experimental Methods for details.

Given the observed potency enhancements, we next explored combining these beneficial mutations and expanding evolution to other positions around the lasso scaffold. As mutations accumulated in individual scaffolds, the wild-type lasso cyclase enzyme required for lassotide folding was found to dictate whether specific analogs were produced or not. Despite these early challenges, over 1000 lassotide variants were generated and screened for integrin binding.

Further lassotide evolution efforts focused on variants **24** (I16Q), **28** (A3K), and **30** (A3R) since these analogs afforded favorable potency increases and were produced with reasonable titers. A series of mutational combinations and 2-site NNK libraries were screened for inhibition of LAP binding to integrins αvβ6 and αvβ8. **Table 3** provides representative SAR data. Substituting the C-terminal glycine of **28** and **30** with lysine afforded analogs **35** and **36**, respectively, which displayed up to 2-fold improved potency and incorporated useful handles for site-specific conjugation. Combining large polar groups at both A3 with I16Q provided variants **37** and **38** with single-digit nM IC50 values for both αvβ6 and αvβ8. Further variation of ring substituents led to additional improvements, although many desired analogs could not be produced using the wild-type cyclase enzyme.

**Table 3.**
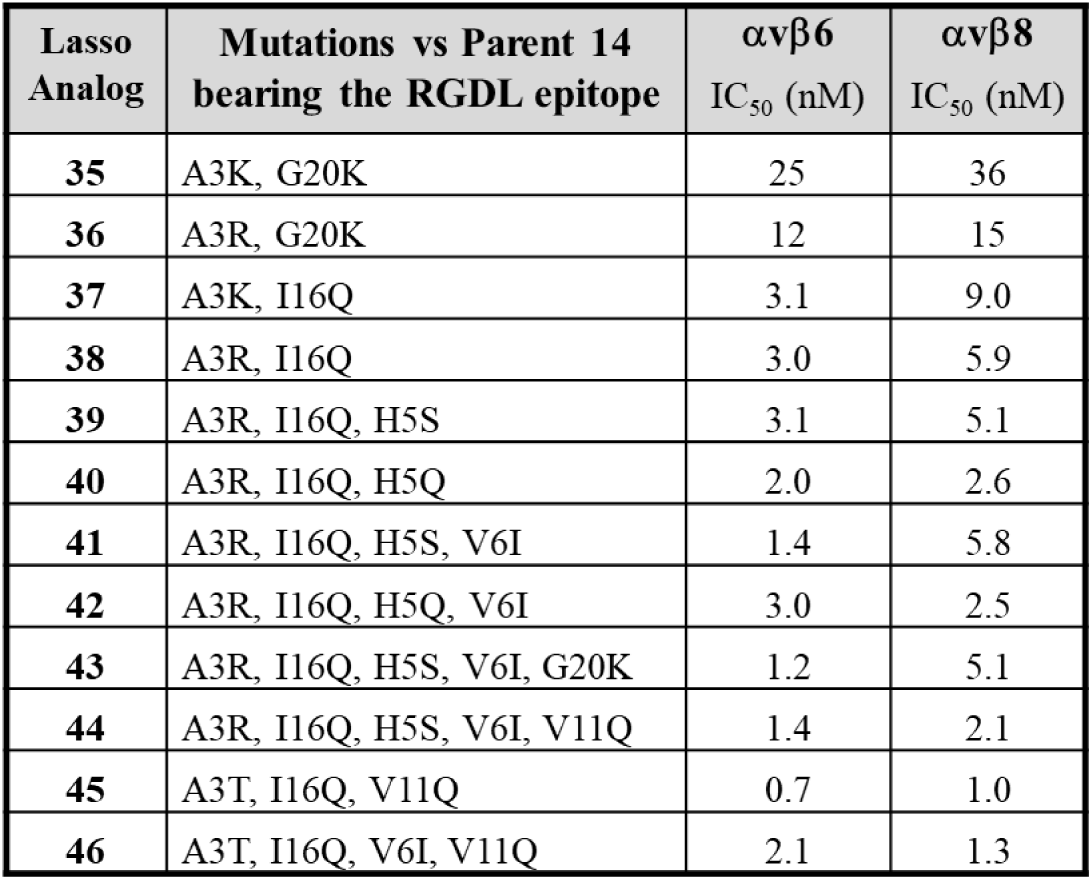
Integrin inhibition data for engineered lasso peptides that combine beneficial mutations at A3, V11, I16, and H5 positions of lassotide **14**. SAR data includes the G20K mutation, which provides a site for conjugation (see below) and was shown to be well tolerated. Lassotide inhibition potency was determined in ELISA competition assays vs latency-associated peptide (LAP). For clarity, inhibition data for integrins αvβ6 and αvβ8 is presented as mean values determined from a minimum of 3 experiments (n ≥ 3). See Experimental Methods for details.

As mutations are accumulated in a given scaffold, the lasso cyclase must fold substrates that are quite distant from the wild-type sequence. Lassotide integrin inhibitors of interest must bear up to 10 mutations vs the parent MccJ25 and many were not formed. Recently we reported an effective strategy involving lasso cyclase engineering to overcome this limitation by broadening amino acid tolerance and enhancing lasso diversification.^32^ Using these improved cyclase enzymes thus enabled the production of lassotides bearing both V11Q and I16Q mutations, leading to variants **44**-**46** that are highly potent inhibitors of both αvβ6 and αvβ8 (IC50 = 1-2 nM). It should be noted that inhibition of mouse and human integrins were virtually identical for all lassotides tested (See Supplementary **Table S1**).

### Lassotide production

Historically lasso peptides have been difficult to advance as viable therapeutic modality due to their inaccessibility by chemical synthesis and generally low levels of biological production, especially when the biosynthetic machinery is tasked with generating mutational variants. Relatively few lasso peptides have been reported to be produced at levels greater than 10 mg/L, and most are produced with barely detectable titers (< 1 mg/L) in their natural hosts.^39,40^ The parent lasso peptide used in this work, MccJ25, has been reported to be produced in *E. coli* at titers of 8-10 mg/L in shake flasks.^41^ In one recent case, the natural lasso peptide capistruin was produced at titers up to 240 mg/L in an engineered version of its wild-type host Burkholderia.^42^ We have found that engineering of the genetic constructs, the *E. coli* host, and lasso cyclase enzymes can greatly enhance production of wild-type and highly mutated lasso peptides.^32^ Thus, production of 572 mg/L of our parent scaffold MccJ25 was demonstrated in shake flasks using *E. coli* that contained the evolved McjC cyclase containing K252A and K388A mutations. To obtain larger quantities of purified analogs required for full pharmacology studies and in vivo efficacy testing, we sought to develop fermentation and downstream recovery processes for production of our lassotide-based integrin inhibitors. Our initial efforts using unoptimized strains under different conditions showed that lassotides **36** and **38** could be produced with titers up to 1.2 g/L and 4.0 g/L, respectively, in controlled fed-batch fermenters (5 L working volume), relative to 18 mg/L and 12 mg/L, respectively, in 2 L shake flasks. In addition, lassotide **35** containing two lysine residues for conjugation was fermentatively produced in four stirred tank reactors (5 L working volume) at an average titer of 5.7 ± 0.10 g/L over 72 hours, which to our knowledge is the highest titer reported to date for lasso peptide production (**Figure 2**). Titers were uniform across the 4 tanks, as indicated by the low SEM, and the high titers of **35** delivered in fermenters vs shake flasks (40 mg/L) correlated with higher cell densities (avg. OD600 = 140 at 72 h in fermenters vs 2-3 in shake flasks). Production was shown to escalate significantly after 30 h when cell growth slowed (see Supplementary **Figure S1**). Lassotides produced in *E. coli* were secreted, presumably by MccJ25 transporter McjD, which facilitated downstream recovery and allowed isolation of >80 g of lassotide **35** from 20 L of fermentation broth. We expect our ability to produce multi-gram to kg quantities of lasso peptides using optimized strains and processes will greatly accelerate development of lassotide-based therapeutics in the future.

**Figure 2.**
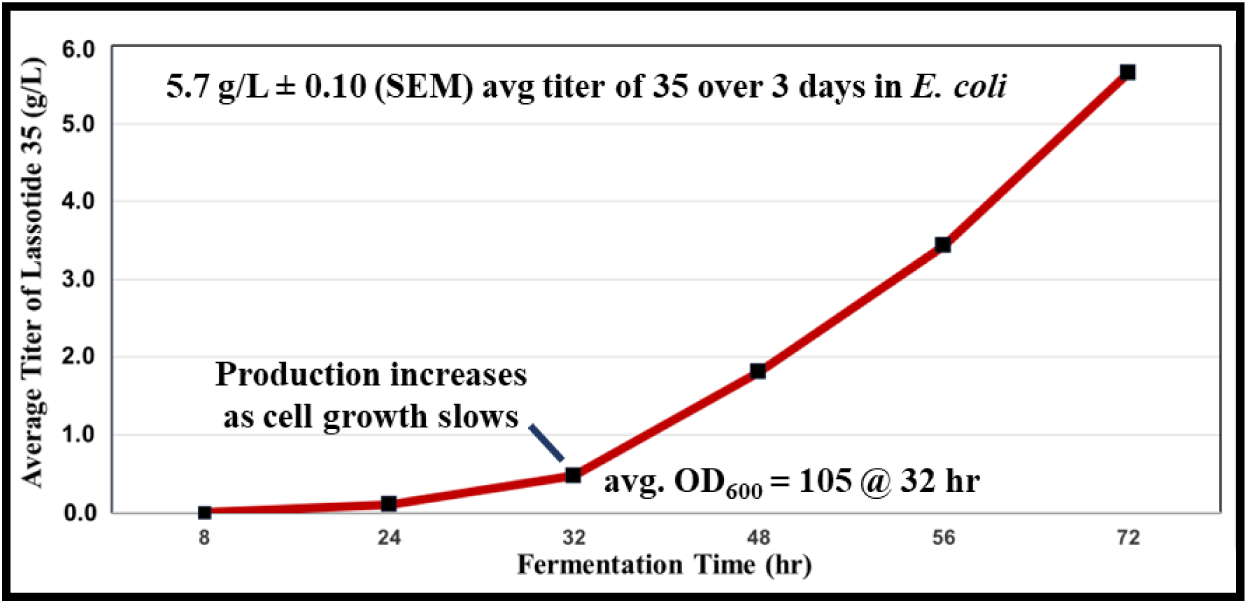
Production time course of lassotide **35** in four fed-batch fermenters (5 L working volume). Reactors were fed based on control of dissolved oxygen (DO) which was maintained at 20%. Production was shown to increase at 32 hours, when cell growth decreased. Average titers over 4 tanks are plotted ± SEM. See Experimental Methods and Figure 1S for further details.

### Lassotide conjugates

Lassotides consist of a modular 3D architecture that potentially allows post-translational modifications at ring or tail residues without undermining loop binding to receptors. For example, conjugation of groups to the C-terminus of a lassotide, beneath the ring, may enable the introduction of a new function into a lassotide scaffold while retaining high target affinity. In addition to exploring the impact of conjugation on integrin binding, we were interested in introducing modifications that could vary properties such as lassotide in vivo half-life. Neri and co-workers reported the small molecule albumin binder 4-(*p*-iodophenyl)buyrate (IPB), which was shown to increase the half-life of various biological conjugates.^43^ Three sites in lassotides **36**-**38** were examined (**Figure 3**). To ensure proper albumin binding by IPB, we introduced a short PEG2 linker to afford the acylating and amination agents **50** and **51**, respectively (**Figure 3A**). Rapid, clean acylation of K20 in **36** and K3 in **37** occurred through standard peptide coupling conditions or via the NHS ester of **50**, affording the conjugated analogs **47** and **48**. PEG2 amine **51** was found to preferentially amidate the C-terminal carboxyl group of **38** to afford **49** vs the side chain carboxyl group of aspartate in the RGDL epitope, possibly due to protective ionic interactions between Asp and Arg in the loop.^44^ Upon testing these conjugates for αvβ6/8 binding, we were encouraged to find that these three modifications did not negatively impact integrin binding, and in fact each conjugate generally showed discernible improvements in inhibition of αvβ6/8 vs the parent lassotides (**Figure 3B**).

**Figure 3.**
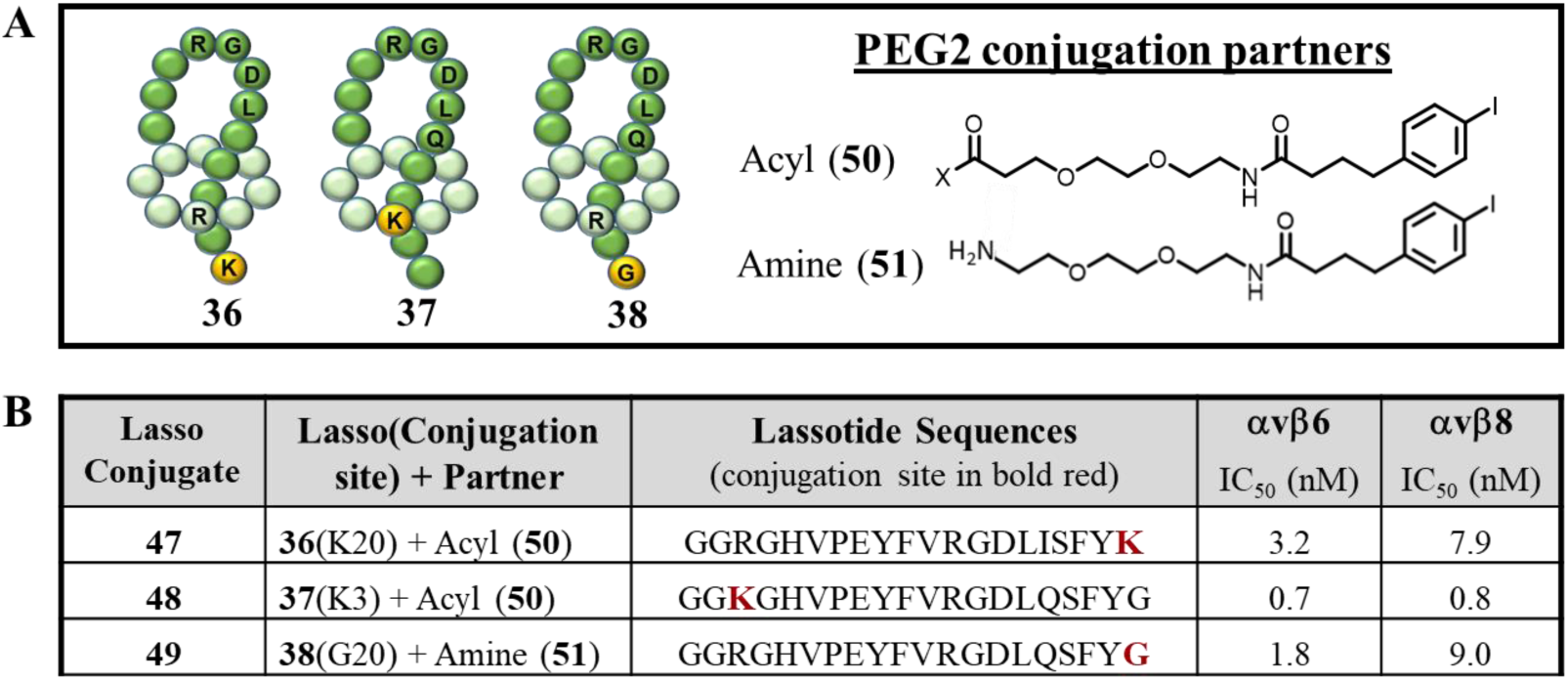
Conjugation studies involving lassotides **36**-**38**. **A.** General structures of lassotides **36**-**38**, showing sites of conjugation as yellow balls, as well as acylating agent (Acyl **50**) and amidating agent (Amine **51**), each bearing a PEG2 linker. **B**. Sequences and IC50 data for lassotide conjugates **47**-**49** vs αvβ6 and αvβ8.

### Integrin selectivity

The RGD-integrins comprise eight cell-surface receptors whose various activities rely on binding to the RGD epitope within their endogenous ligands.^6,45^ Given the broad functions of integrins, the safety of integrin inhibitors often depends on achieving a high degree of selectivity for integrins specifically linked to the pathology of a targeted disease. A lack of integrin selectivity has been associated with adverse events in clinical testing of integrin inhibitors.^46,47^ In our case, a dual integrin inhibitor was sought and a high degree of selectivity for αvβ6 and αvβ8 was desired to minimize adverse off-target effects. Therefore, we examined the selectivity of our lassotide-based inhibitors across the eight RGD-integrins. A set of three pairs of integrin inhibitors were tested, which consisted of the parent lassotides **36**, **37**, and **38**, along with their conjugated analogs, **47**, **48**, and **49**, respectively. The non-selective small molecule inhibitor MK-0849^48^ was used as the integrin assay positive control for all integrins except the platelet integrin αIIbβ3, where the selective inhibitor Tirofiban^49^ was employed. As shown in **Table 4**, all parent lassotides showed selectivity for αvβ6/8 relative to the other six RGD integrins, although **37** and **38** did inhibit αvβ1 with reasonable potency. The low observed inhibition of αIIbβ3 was desirable since blockade of this integrin compromises platelet aggregation and can lead to severe bleeding events.^47,50^ We were pleased to see that all conjugates (**47**, **48**, and **49**) not only displayed higher potency for αvβ6/8, but also exhibited significantly higher selectivity vs the other six RGD-integrins. In particular, conjugate **47** presented with 2-3-fold improvement in potency and >125-fold selectivity against all other RGD-integrins. Given the strong potency and selectivity displayed by **47**, studies described below focus on the properties and performance of this analog.

**Table 4.**
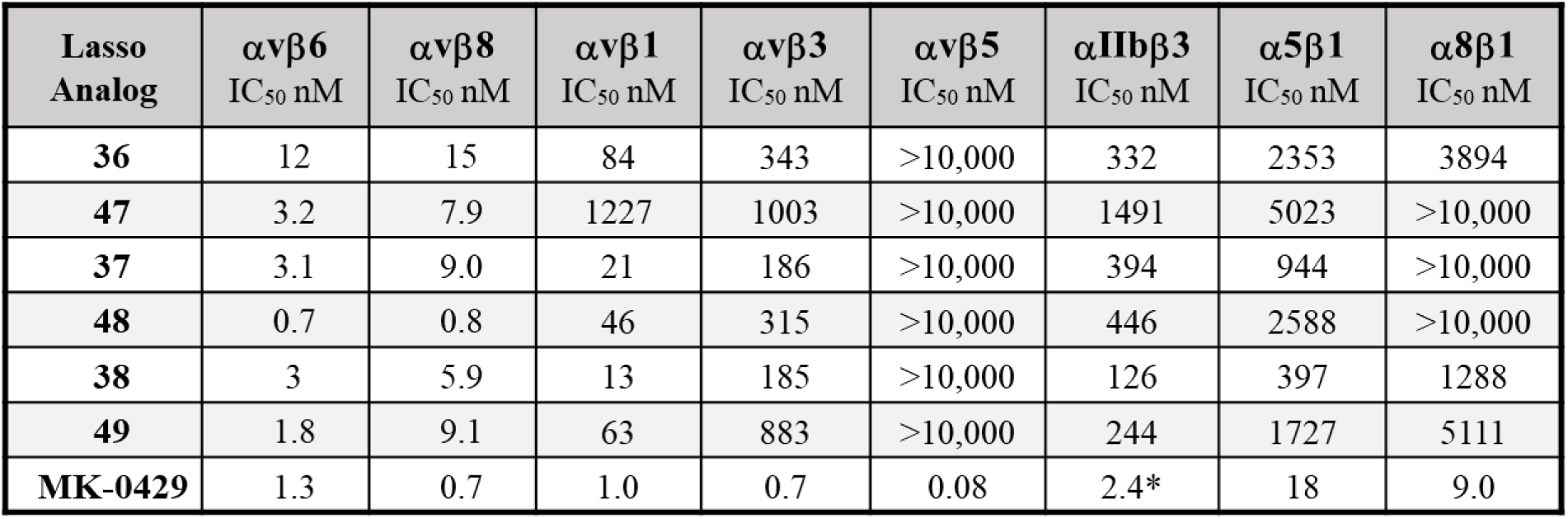
RGD-integrin selectivity of lassotides **36**-**38** and conjugates **47**-**49** (shaded), as measured by competitive inhibition vs LAP in ELISA assays against integrins αvβ3, αvβ6 and αvβ8 (as described above) and αvβ1, αvβ5, αIIbβ3, α5β1, and α8β1. *MK-0429^48^ was used as a positive control for all integrin assays except αIIbβ3, where Tirofiban^49^ was employed.

### Physicochemical properties

The general structure and physicochemical properties of lassotide **47** are shown in **Figure 4**. Of note is the good solubility (30 mg/mL) of **47** in simple acetate buffer at pH 4.5 with a clear colorless solution and no aggregation detected. Other solvents like DMSO and formulations containing 10-20% DMSO and PEG for in vivo studies allowed solubilities >200 mg/mL to be achieved.

**Figure 4.**
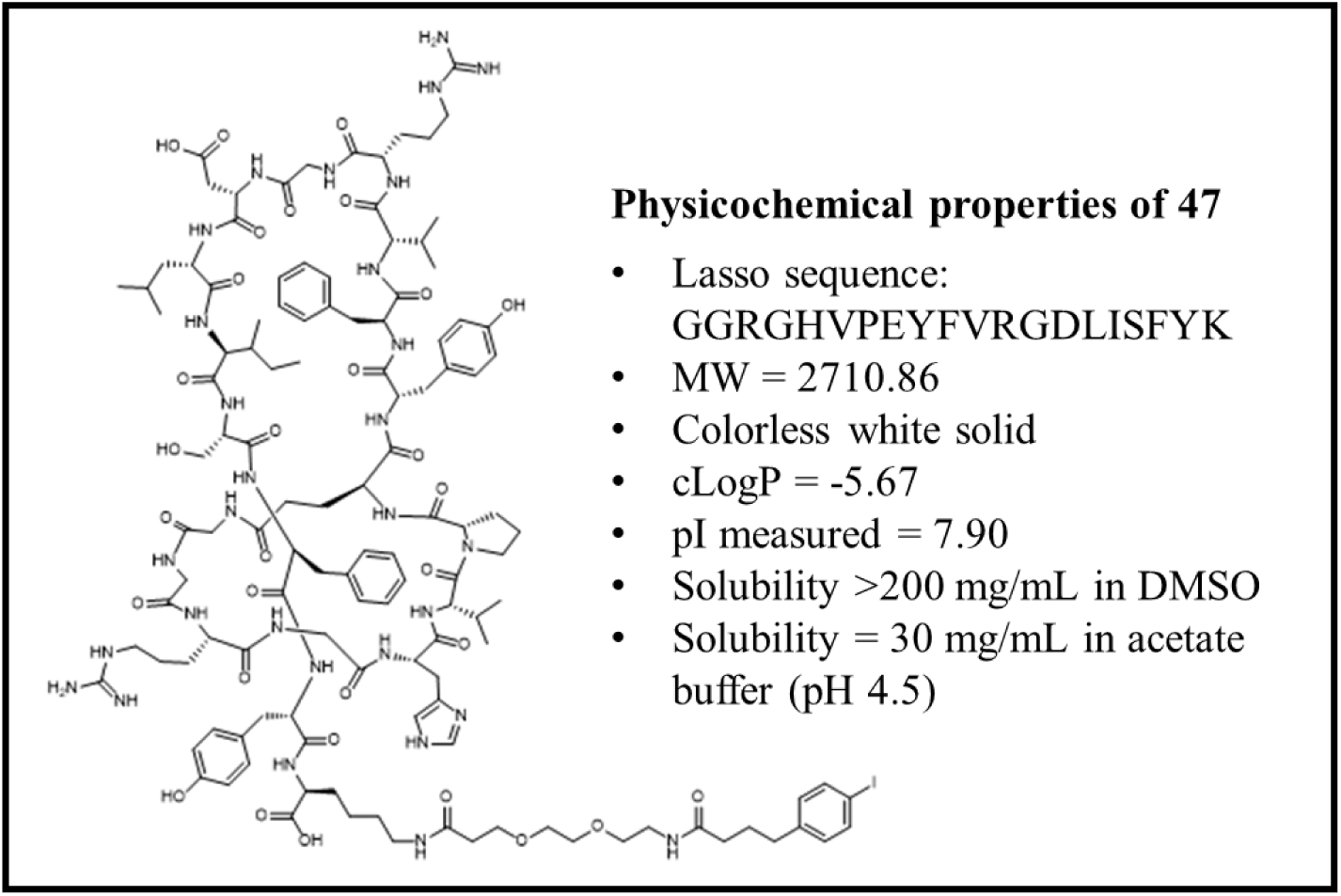
Structure and physicochemical properties of lassotide **47**.

### In vitro pharmacology and safety

Due to the promising potency, integrin selectivity, and solubility observed for analog **47**, comparative in vitro characterization was conducted on this conjugated lassotide and its parent **36**. Lasso peptides are known for their thermal and proteolytic stability^39,51^ and all lassotide integrin inhibitors examined, including **36** and **47**, were found to be robust in standard heat (95°C for 1 h) and proteolysis (carboxypeptidase Y treatment for 4 h) tests with little or no change observed in the HPLC traces. Interest in oral delivery led us to examine the stability of **36** and **47** in simulated intestinal fluid (SIF), which consists of trypsin, chymotrypsin, elastase, and carboxypeptidases that manifest a harsh proteolytic environment. Both lassotides **36** and **47** were highly susceptible to proteolytic cleavage by SIF with t1/2 < 2 min. Product analysis by LC-MS/MS indicated that rapid cleavage at the R-G linkage in the loop region occurred and that trypsin was responsible for the degradation. Trypsin is known to be selective for cleaving peptides and proteins exclusively C-terminal to Arg and Lys residues.^52^ Interestingly, no cleavage of the R-G linkage in the rings of **36** or **47** occurred and generation of a variant of **36** without RGDL in the loop led to prolonged SIF stability with t1/2 = 16 h. Hence, the present lassotides containing RGD in the loop region may be inherently unsuitable for oral delivery without further modifications.

Examination of mouse and human plasma and hepatocyte stability showed lassotides **36** and **47** to be generally resistant to plasma proteases and intrinsic liver metabolism (**Figures 5A** and **5B**). Interestingly, lassotide **47** was somewhat more susceptible than **36** to mouse hepatocyte degradation, possibly due to enzymes that are metabolizing the conjugated albumin-binding *p*-iodophenylbutyrate moiety in **47**. Plasma protein binding (PPB) studies on **36** and **47** demonstrated that the albumin-binding group does lead to significant increase in PPB, as expected, with lassotide **47** displaying >99% bound fraction in both human and mouse plasma samples vs 87% and 78%, respectively, for unconjugated **36** (**Figure 5C**).

**Figure 5.**
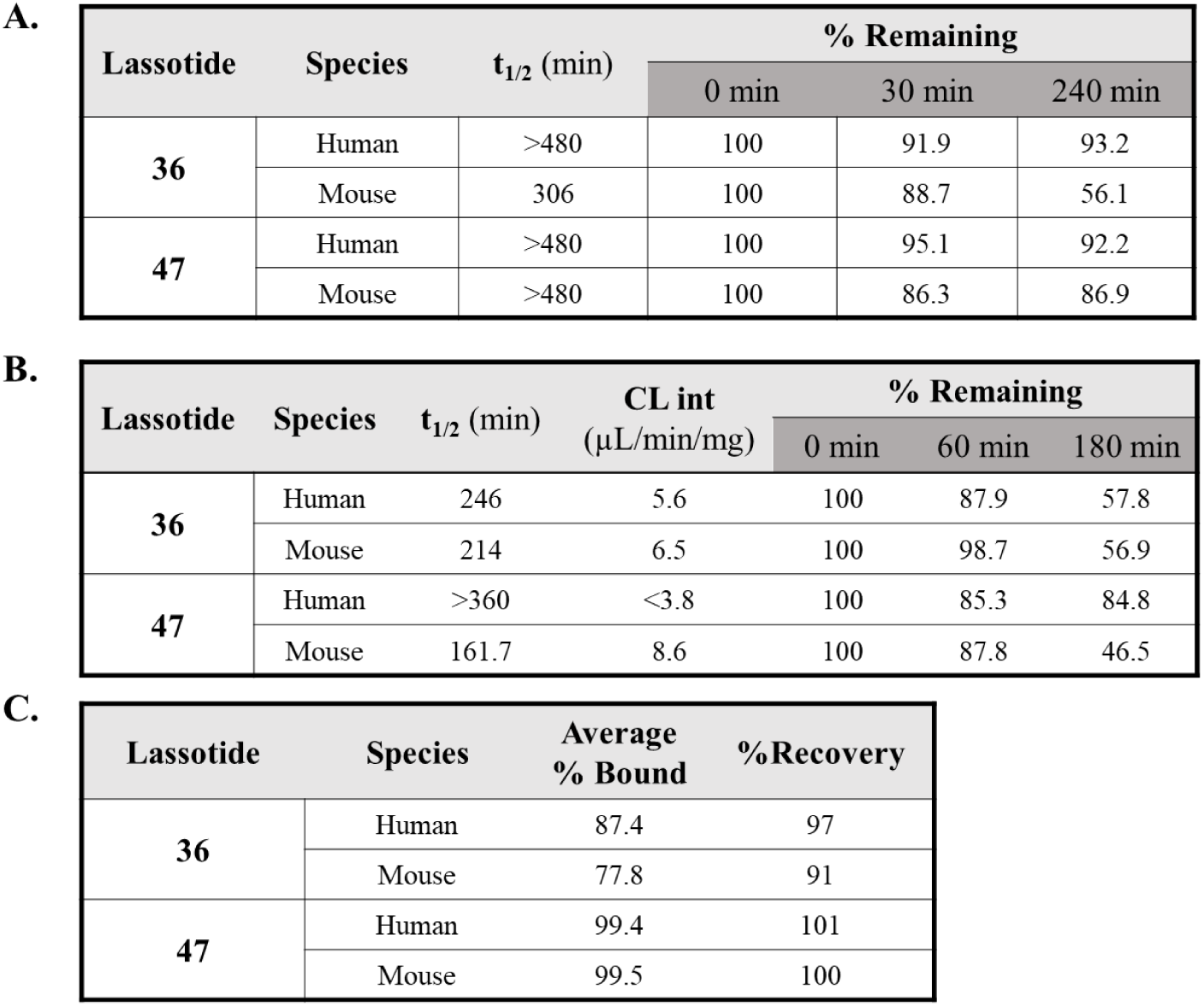
In vitro metabolic stability and plasma protein binding for lassotide **36** and conjugated derivative **47**. **A**. Plasma stability data, as measured by incubating **36** and **47** in human and mouse plasma. **B.** Hepatocyte metabolism stability data, as measured by incubating **36** and **47** with cryopreserved human and mouse hepatocytes. **C.** Plasma protein binding (PPB) data as measured by rapid equilibrium dialysis. Experiments were performed in duplicate, and data are presented as mean values. Lassotide concentrations = 2 μM in all studies. See Experimental Methods section for details.

Finally, in vitro safety pharmacology was performed on lassotides **36** and **47** through a detailed Safetyscan47 analysis (see Eurofins; https://www.eurofinsdiscovery.com/catalog/safety47-panel-dose-response-safetyscan-discoverx/87-1003DR). In this study, responses were tested against 78 different enzymes, transporters, ion channels, and receptors, many of which are known to be involved in adverse drug reactions (e.g., hERG).^53^ We were encouraged to see that neither lassotide **36** nor **47** elicited a response signal (RC50) up to 40 μM concentrations, indicating a very clean safety profile for both compounds (see Supplementary **Tables S2A** and **B**). Thus, adverse off-target effects often seen with small molecules are not expected with these lassotide-based drugs.

### In vivo properties

#### Pharmacokinetics

Pharmacokinetic (PK) measurements were conducted to assess the in vivo absorption, distribution, metabolism, and clearance of lassotides for the first time in mice and rats. In particular, Cmax, AUC (systemic exposure), and half-life (t1/2) can greatly impact the ability of lassotides to engage and modulate receptor function sufficiently to achieve the desired efficacy response in a disease model. Hence, the PK properties of lassotides **36** and **47** were examined to compare inherent in vivo behavior and understand the impact of conjugation at different doses and routes of administration. As shown in **Table 5**, unconjugated lassotide **36** dosed at 30 mg/kg in female Balb/c mice displayed modest Cmax and AUC values and a short half-life of 0.6 h. Lassotide **47** conjugated with an albumin-binding group exhibited a 39-fold increase in AUC and 9-fold improvement in t1/2 (5.5 h), consistent with greater plasma protein binding for **47** vs **36**. Dosing **47** at the same levels in rats also presented a favorable PK profile with a half-life of 8.5 h. Comparative dosing of **47** by IP and SC routes at 10 mg/kg in mice revealed virtually identical PK parameters, demonstrating the potential of subcutaneous administration for this lead compound. Interestingly, double acylation with Acyl **50** at K3 and K20 of lassotide **35** afforded the derivative **52** with two albumin-binding groups attached. IP dosing of **52** at 10 mg/kg in mice showed a further 4.4-fold increase in half-life (t1/2 = 20 h), demonstrating our ability to rationally modulate the half-life of lassotides.

**Table 5.**
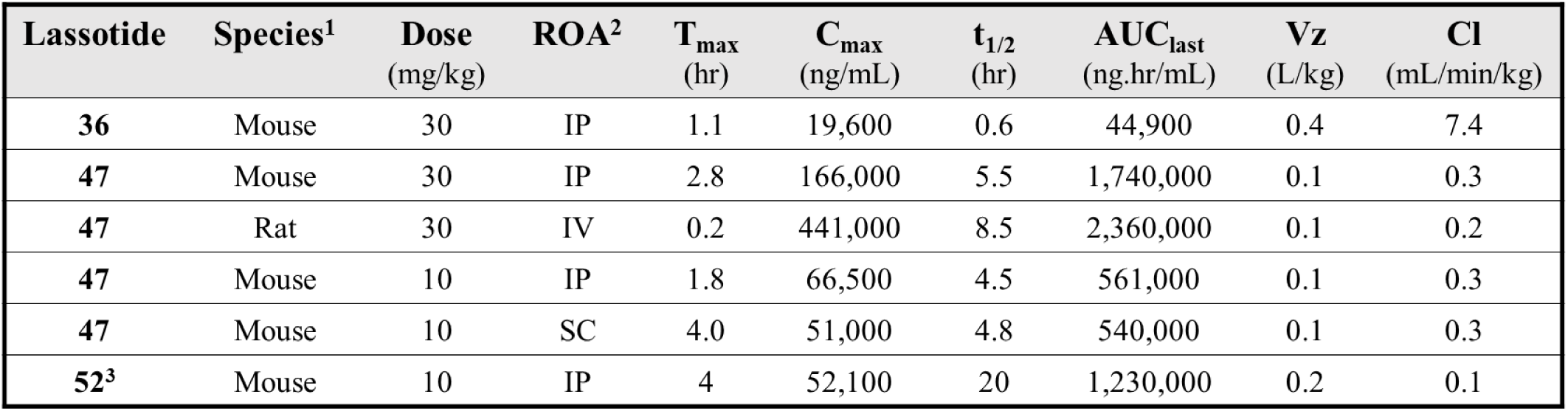
Pharmacokinetic data for lassotides **36**, **47**, and **52**. Lassotides were formulated in 10% DMSO, 30% PEG400, and 60% water. Timepoints were collected at: 10 min, 30 min, 1, 2, 4, 8, 12, and 24 hr (48 h timepoint was collected for longer half-life lassotide **52**). PK analytical data was processed using the Phoenix WinNonlin modeling software involving non-compartmental analysis (NCA) and statistical testing (ANOVA). ^1^ In mouse studies at 30 mg/kg, 3 or 5 female Balb/c mice were used; at 10 mg/kg, 5 female Balb/c mice were used. In rat study, 3 female Sprague-Dawley rats were used. ^2^ Route of administration (ROA) involved single-dose injections. IP = intraperitoneal; IV = intravenous; SC = subcutaneous. ^3^ Compound **52** is the doubly conjugated derivative formed by reacting lassotide **35** (K3, K20) with Acyl **50**. Data are presented as mean values for clarity. Full data for individual animals with statistical analysis is provided in Supplementary **Tables S3A-E**.

#### Tissue distribution

To gain a deeper understanding of how well lasso peptides distribute into tissues, lassotide **47** was IP injected into 6 female Balb/c mice and a range of tissues were collected at 3 h (n = 3) and 8 h (n = 3) timepoints. As shown in **Table 6**, high distribution was observed in ovary, breast, lung, and kidney tissues, which are important as targets for potential treatment of αvβ6/8-expressing solid tumors in these organs.^8,12,13,14^ Also, adequate distribution of **47** was seen in the liver and colon. Interestingly, modest distribution of **47** also was seen in the brain, which could provide an avenue through optimized lassotides for treating brain-associated diseases such as gliomas and brain metastases that overexpress αvβ6/8.^54,55,56^

**Table 6.**
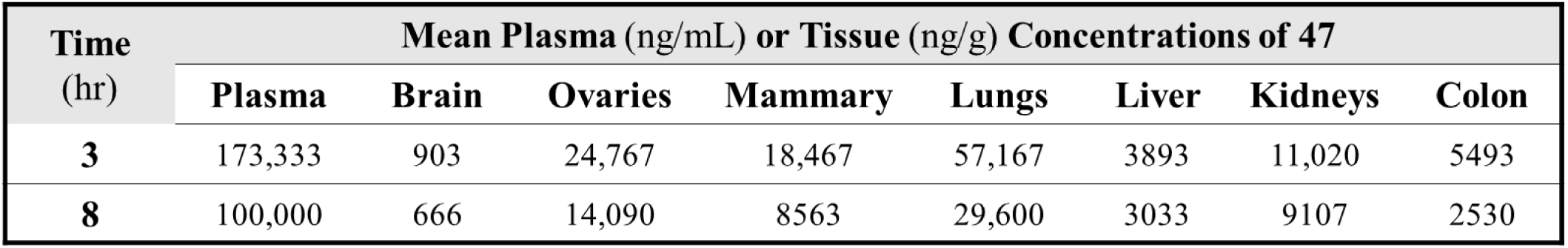
Tissue distribution data for lassotide **47** in mice. Six Balb/c female mice were dosed at 30 mg/kg by IP injection. Three mice were sacrificed and tissues bioanalyzed at 3 h and three were analyzed at 8 h. Data are presented as mean values. See Supplementary **Table S4** for individual mouse data and statistics.

#### Anti-tumor efficacy

Lassotide **47** displays desirable drug-like properties, and we next explored whether these observations would translate to in vivo anti-tumor efficacy. The presence of αvβ6 and αvβ8 on tumors and/or Tregs in the TME has been shown to liberate TGF-β,^19,21,23^ which is a strong immunosuppressant that serves as a primary resistance mechanism for immune checkpoint inhibitors such as anti-PD-1 monoclonal antibodies.^16,57,58^ Thus, blockade of TGF-β represents an attractive strategy to enhance the response and survival rates of existing immunotherapies. Recent proof-of-concept for the effectiveness of this approach was reported for bintrafusp alfa, a biologic therapy that targets both TGF-β and PD-L1 and showed improved survival in a murine model of ovarian cancer.^22^ Similarly, the anti-latent TGF-β mAb, SRK-181, was found to complement anti-PD-1 therapy and enhance efficacy in mouse tumor models, including EMT6.^21^ Despite positive efficacy, safety is a concern with these agents as direct systemic blockade of TGF-β is known to elicit a host of adverse events.^24,25^

Triple negative breast and ovarian cancer cells have been shown to overexpress αvβ6 and αvβ8 and are resistant to checkpoint immunotherapy wherein low response rates have led to few approvals for these two solid tumor types.^59,60^ Initial efficacy studies aimed at testing the relative performance of lassotides **36** and **47** in the syngeneic EMT6 mouse model of anti-mPD-1-resistant triple negative breast cancer.^61^ Reaction Biology (Malvern, PA, USA; https://www.reactionbiology.com) performed these studies, and they have demonstrated that this model highly expresses both αvβ6 and αvβ8 (see Supplemental **Figure S4**). Fast-growing EMT6 cells were implanted orthotopically into the inguinal mammary fat pads of female Balb/c mice (See Experimental Methods for details). Six groups of mice (n = 8) bearing EMT6 tumors (avg starting tumor volume 75 mm^3^) were treated with (i) vehicle + isotype control (clone 2A3), (ii) anti-mPD-1 (clone RMP1-14), (iii) lassotide **36** + isotype control, (iv) lassotide **47** + isotype control, (v) **36** + anti-mPD-1, and (vi) **47** + anti-mPD-1. A small initial dose range finding study indicated that lassotides **36** and **47** could be tolerated up to 200 mg/kg. Therefore, **36** and **47** were dosed IP at 100 mg/kg (BID and QD, respectively) and anti-mPD-1 was dosed IP at 10 mg/kg x 3 over the course of the 23-day dosing schedule. As shown **Figure 6**, lassotides **36** and **47** alone, and lassotide **36** in combination with anti-mPD-1 displayed little anti-tumor activity, similar to anti-mPD-1 alone. By contrast, the combination of lassotide **47** + anti-mPD-1 (blue trace) elicited a robust anti-tumor response with high statistical significance. Further efficacy studies focused on the lassotide **47** + anti-mPD-1 combination therapy.

**Figure 6.**
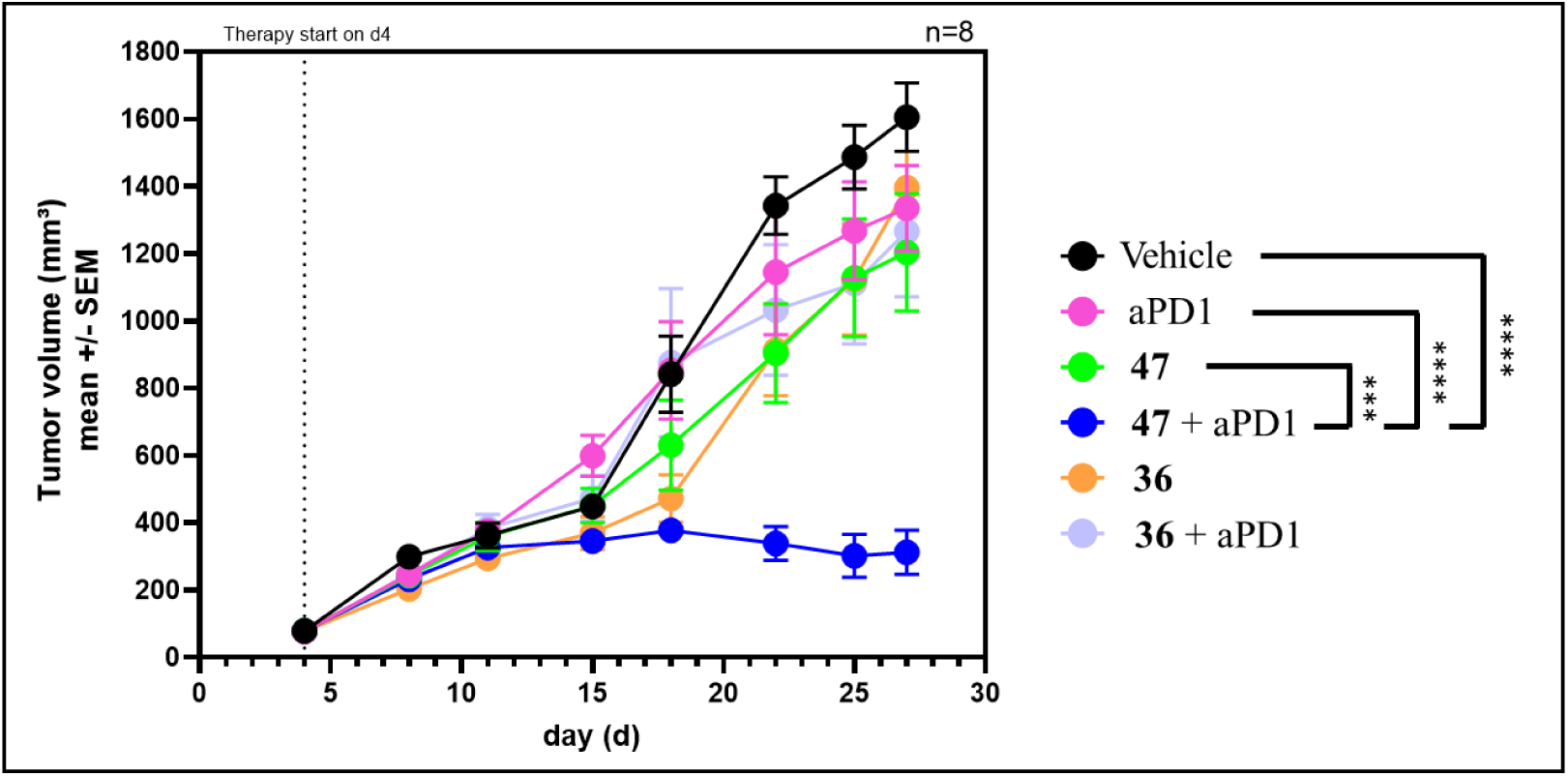
In vivo anti-tumor efficacy of lassotides **36** and **47** in combination with anti-mouse-PD-1 (aPD1). Six groups of female Balb/c mice (n = 8) were implanted with EMT6 tumors on day 0 and dosing did not start until day 4 when the average tumor volume reached 75 mm^3^. Lassotides **36** and **47** were formulated in 10% DMSO, 30% PEG400, 60% sterile water (Vehicle) and were IP dosed BID and QD, respectively, at 100 mg/kg starting on day 4. Anti-mPD-1 (Clone RMP1-14 listed as aPD1) was dosed IP at 10 mg/kg on days 4, 8, and 12. Vehicle and solo lassotide groups include isotype control antibody (clone 2A3). Data are presented as mean ± SEM. Data analysis and statistics between different groups were performed using a 2-way ANOVA model where p-values between groups are indicated and shown as *** p<0.001 or **** p<0.0001.

A second study involving the syngeneic EMT6 tumor model was conducted to explore the dose response of lassotide **47** in combination with anti-mPD-1. In this experiment, three groups of female Balb/c mice (n = 12) bearing EMT6 tumors were IP dosed once daily (QD) for 23 days with **47** at 10 mg/kg, 30 mg/kg, or 100 mg/kg (each in combination with anti-mPD-1). Two additional groups (n = 12) were dosed with vehicle + isotype control and solo anti-mPD-1, as described above. As shown in **Figure 7**, a classic dose response was observed with higher doses synergizing with anti-mPD-1 to deliver increasingly robust anti-tumor efficacy. The results for the 30 mg/kg and 100 mg/kg treatment groups display high statistical significance and 3 tumors fully regressed in the highest dosed group (see photos and individual mouse graphs in Supplemental **Figure S5**).

**Figure 7.**
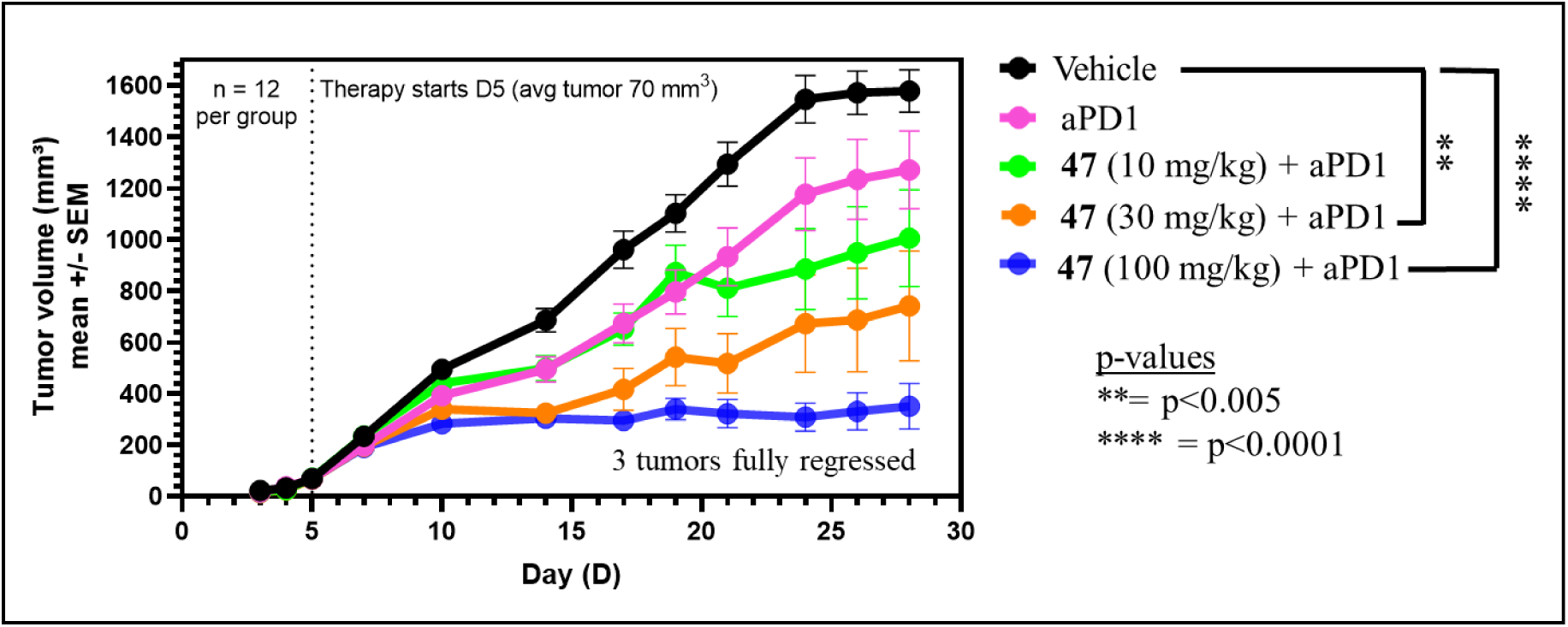
Anti-tumor efficacy dose response of **47** + anti-mPD-1 in EMT6 murine tumor model. aPD1 = anti-mPD-1. Vehicle = 10% DMSO, 30% PEG400, 60% water + Isotype control antibody (clone 2A3). Data are presented as mean ± SEM. Data analysis and statistics between different groups were performed using a 2-way ANOVA model where p-values between groups are indicated.

A third efficacy study was conducted by LabCorp (Burlington, NC, USA) and was structured to demonstrate anti-tumor activity against anti-PD-1-resistant ovarian cancer. LabCorp has developed the ID8-Luc-mCh-Puro.TD1 (ID8) syngeneic orthotopic model derived from a transformed ovarian epithelial cell line obtained from C57BL/6 mice. ID8 cells (1 x 10^7^) were implanted into the peritoneum of three groups of C57BL/6 mice (n = 7) and, after 17 days of growth, tumors were IP dosed over 28 days with vehicle + isotype control, anti-mPD-1, or **47** (100 mg/kg QD) + anti-mPD-1. Tumor growth was accurately monitored weekly through whole body bioluminescence imaging (BLI).^62^ As illustrated in **Figure 8**, the combination of **47** + anti-mPD-1 provided strong statistically significant efficacy wherein 5 out of 7 tumors regressed, 2 were complete regressions, and 1 mouse was a tumor-free survivor (see insert in **Fig 8A** graph showing individual mouse responses). Moreover, all mice were monitored for survival over an additional 25 days after treatment was halted. As shown in the Kaplan Meier survival curves (**Fig 8B**), the **47**/anti-mPD-1 combination was found to provide a significant durable survival benefit wherein 6 out of 7 mice in this treatment group survived and all other mice in the study succumbed to tumor burden before day 70. Weekly BLI measurements showed that tumors in the **47**/anti-mPD-1 treatment group did not grow during the 25-day survival phase after dosing terminated (Supplementary **Figure S6**).

**Figure 8.**
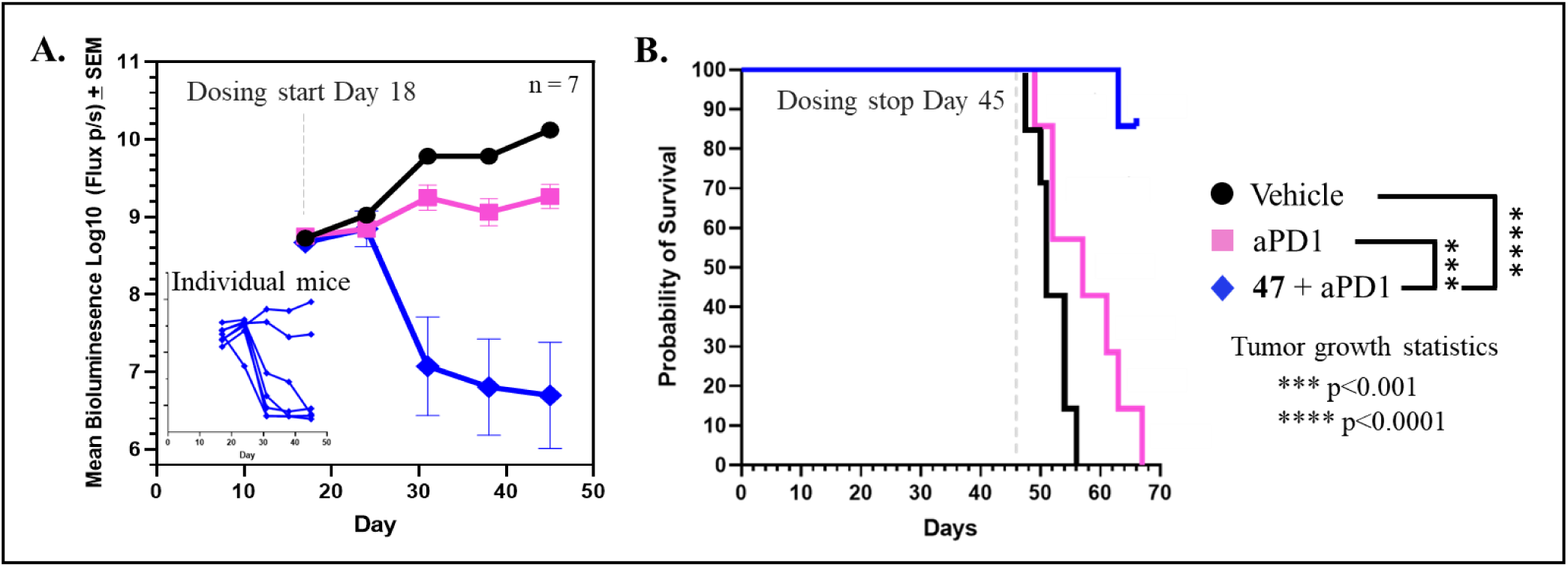
Anti-tumor efficacy of **47** + anti-mPD-1 in ID8 ovarian cancer model. **A**. Tumor growth responses as measured by whole body BLI. Three groups of female albino C57BL/6 mice (n= 7) were implanted with ID8-Luc-mCherry-Puro.TD1 cells intraperitoneally. Treatment started on day 18 with anti-mPD-1 solo (10 mg/kg x 4) and **47** (100 mg/kg QD) + anti-mPD-1. Anti-mPD-1 was dosed on days 18, 21, 25 and 28. Data are presented as mean ± SEM. Data analysis and statistics between different groups were performed using a 2-way ANOVA model where p-values between groups are indicated. **B**. Kaplan Meier survival curves for each treatment group over 70 days.

## Discussion

This study has demonstrated the facility with which lasso peptides can be engineered into highly potent and selective dual inhibitors of the integrins αvβ6 and αvβ8. High integrin inhibition potency, as measured by competitive binding ELISA assays vs the LAP protein, was achieved by a combination of (i) RGDL epitope scanning across the lasso peptide structure, (ii) computational modeling and design, and (iii) directed evolution involving residues in the loop and ring proximal to the RGDL epitope. As reported by Hegemann et al.,^34^ RGD in the loop of the natural scaffold MccJ25 was found to strongly inhibit αvβ3, but we found this derivative only weakly inhibited αvβ6 and αvβ8. Our ability to delete loop residues and ultimately expand the range of mutational variants that could be produced using engineered lasso cyclases^32^ enabled rapid progression to low nM/high pM inhibition of the desired integrins. Integrin selectivity is recognized as a critical factor that can impact inhibitor safety and efficacy.^7,46,63,64^ Importantly, we have discovered numerous lassotides that displayed high selectivity for the two desired RGD-integrins vs the six other integrins known to bind the RGD epitope. In particular, half-life extended **47**, which is acylated at C-terminal K20 of lassotide **36** with PEG2-IPB (Acyl **50**), displayed >125-fold selectivity for αvβ6 and αvβ8 over all other RGD-integrins. These results illustrate that all regions of the modular 3D lassotide structure may be modified to enhance their therapeutic attributes.

Lasso peptides represent an emerging therapeutic modality with advantageous drug-like properties. Lasso peptides have been reported to be thermostable and proteolytically resistant.^28,39,51^ Lassotide-based integrin inhibitors **36** and **47** showed promising physicochemical and in vitro properties, including high solubility (e.g., 30 mg/mL in acetate buffer), high stability to plasma and liver metabolism, a favorable safety profile, and tunable plasma protein binding which impacted in vivo PK and efficacy. However, **36** and **47** were found to be highly susceptible to trypsin cleavage at the R-G linkage in the loop, which will limit the potential for oral delivery of these integrin-targeted lassotides without further engineering to block this activity.

The advancement of lasso peptides as a useful drug modality has been hindered by the general inability to produce these naturally-derived products in quantities required for extensive in vitro and in vivo testing.^40^ The natural biosynthetic pathways and enzymes are not designed to produce large amounts of these products. This becomes even more problematic when mutations are introduced into the lasso substrates and the enzymes are tasked with converting a “foreign” precursor. Accordingly, we developed methods for engineering the pathway genetic constructs and the *E. coli* host, along with engineering of the lasso cyclase enzyme to be more tolerant to mutational variations.^32^ While optimization continues, we were able to demonstrate significant improvement in lassotide production, with lassotide **35** being formed at a titer of 5.7 g/L, the highest level reported for lasso peptides to date. These results establish a simple, low-cost, single-step fermentation process for producing ample quantities of lassotides and removes a barrier that has been thwarting therapeutic development of these useful molecules.

As a consequence of lasso peptide production constraints, only limited in vivo data previously has been reported and has largely been confined to wild-type MccJ25.^65,66,67,68^ Herein we have provided the first detailed in vivo characterization of engineered lasso peptides, including PK and efficacy studies demonstrating the strong anti-tumor activity of lassotide **47** in combination with anti-mPD-1 therapy. The integrin inhibitors developed here are based on the MccJ25 scaffold and, like the parent,^65^ our αvβ6/8-targeted lassotides have relatively short half-lives in mice (< 1 h), presumably due to rapid renal excretion. Accordingly, half-life extension techniques were explored. Typical lipid-based (e.g., stearate, palmitate) conjugates^69^ were readily formed by acylating Lys residues within lassotides such as **36**. However, solubilities suffered, and the lipid tails tended to interfere with our integrin binding assays. To address this challenge, we introduced the small molecule albumin binder 4-(*p*-iodophenyl)butyrate (IPB),^43^ which afforded lassotide **47** and increased half-life to 5.5 h and 8.5 h in mice and rats, respectively. In addition to t1/2, AUC values increased substantially (39-fold) and the enhanced exposure of **47** likely contributed to the strong efficacy we observed in murine tumor models. Another indicator of favorable systemic exposure for lassotide **47** was revealed through preliminary tissue distribution studies (**Tables 6 and S4**), which showed good concentrations of **47** in the key organs that would be candidates for solid tumor immunotherapy. Importantly, this data showed adequate distribution of **47** to ovaries and mammary tissues of Balb/c mice, which were the primary sites for our initial anti-tumor efficacy studies.

Our rationale for developing lassotides that potently and selectively inhibit αvβ6 and αvβ8 was based on the TGF-β-activating function of these two integrins in the tumor microenvironment (TME).^16,19,20,57,58^ In particular, the presence of these integrins on tumors and/or Tregs in the TME liberates TGF-β, which is a strong immunosuppressant that serves as a primary resistance mechanism for immune checkpoint inhibitors such as anti-PD-1 antibodies.^10,11,12^ Dual inhibition of αvβ6/8 integrins and the associated reduction of immunosuppressive TGF-β in the TME was expected to enhance the performance of anti-PD-1 therapy, which is known to be efficacious against a range of tumor types^70,71^ yet is subject to various resistance mechanisms.^72,73^ Thus, our objective was to sensitize anti-PD-1-resistant tumors using dual αvβ6/8 inhibitors.

Lassotides **36** and **47** initially were examined for anti-tumor efficacy alone and in combination with anti-mPD-1 therapy in the anti-PD-1-resistant EMT6 syngeneic mouse model of triple negative breast cancer. Several observations can be gleaned from **Figures 6** and **7**: (i) As expected, anti-mPD-1 alone showed little efficacy in this model, presumably due to the immunosuppressive environment and the presence of TGF-β-activating integrins that may contribute to diminished CD8+ T cell activity, thus leading to anti-mPD-1 resistance. (ii) Neither **36** nor **47** solo led to significant tumor growth reduction, which is consistent with the presence of an active PD-1/PD-L1 immune checkpoint pair in these tumors. Inhibiting αvβ6/8 integrins and reducing TGF-β alone appears to be insufficient to overcome immune checkpoint-driven tumor growth. (iii) Despite being dosed twice per day, lassotide **36** in combination with anti-mPD-1 showed little anti-tumor efficacy, which likely is associated with the short half-life and inadequate exposure of this integrin inhibitor, and (iv) Lassotide **47** in combination with anti-mPD-1 exhibited robust anti-tumor activity, essentially halting average tumor growth and leading to regression of numerous tumors. While further work is required, this data indicates that our lassotide-based dual αvβ6/8 integrin inhibitors synergize effectively with anti-PD-1 therapy to elicit strong tumor reduction/regression. Moreover, the difference in performance of lassotides **36** and **47** suggests that half-life extension and increased systemic exposure of **47** were critical factors for achieving compelling efficacy.

Notwithstanding approvals of dostarlimab and pembrolizumab for treating a subset of advanced ovarian cancers that have high microsatellite instability (MSI-H), DNA mismatch repair deficiency (dMMR), and/or high tumor mutational burden (TMB-H), immune checkpoint inhibitors have performed poorly in treating more prevalent epithelial ovarian cancer.^60,74,75^ There is a critical unmet need for improved treatments for this pernicious cancer. **Figure 8** shows that treatment of ID8 mouse ovarian cancer with the combination of **47** and anti-mPD-1 for 28 days led to regression of 5 out of 7 tumors. Importantly, tumor regression appeared to be durable and correlated with survival, with 6 out of 7 mice remaining alive and no further tumor growth for 25 days following treatment termination. These promising results indicate that dual αvβ6/8 therapeutics may be an effective combination partner that enhances the anti-tumor efficacy of immune checkpoint inhibitors in cases where integrin-activated TGF-β contributes to resistance.

### Conclusions

Overall, the results described herein highlight the utility of lasso peptides as a useful and versatile therapeutic modality that can be engineered for desirable potency, selectivity, and in vivo performance. The stability of lasso peptides should contribute to favorable safety properties, as our initial data suggests. Moreover, the promising efficacy of our dual αvβ6/8 inhibitor **47** in combination with anti-mPD-1 indicates a path forward for developing highly effective combinations with immune checkpoint inhibitors for tumors that display resistance to immunotherapies. Future studies will expand on the results presented herein and explore a wider range of tumor models. Engineering advanced lassotides based on **47** that display low pM affinity for αvβ6 and αvβ8 will be sought without compromising RGD-integrin selectivity. Such potency improvement may permit more complete tumor regression at lower effective doses. In addition, further half-life extension, such as shown for lassotide **52**, could allow less frequent subcutaneous dosing from once daily to weekly or longer. Finally, the small compact size and stability of lassotides may allow the development of oral therapeutics that would eliminate the need for injections and potentially enable alternatives to injected biologic drugs in the future.

### Experimental Methods

#### General Methods

Synthetic DNA primers and ultramer oligos were purchased from Integrated DNA and codon optimized genes were purchased from Twist Biosciences. New England Biolabs (Ipswich, MA) supplied the Q5 polymerase, T4 DNA ligase, NEBuilder® HiFi DNA Assembly, polynucleotide kinase, and Dpn1 enzymes for molecular cloning. Other reagents for molecular biology were purchased from Thermo Fisher Scientific (Waltham, MA) or Gold Biotechnology Inc. (St. Louis, MO). *E. coli* Dh5α was used for plasmid maintenance while BL21(DE3) was used for protein purification, lasso peptide production, and cell lysate production. Both competent cells were made in house unless otherwise noted. Chemical reagents used in this study were from Sigma-Aldrich (St. Louis, MO) and used without purification. All molecular biology reactions are conducted using standard plates, tubes, vials, and flasks typically employed when working with biological molecules such as DNA, RNA, and proteins. Amicon Ultra-4 centrifugal filters (30 kDa, 15 mL and 500 µL) were purchased from EMD Millipore. GENEWIZ from Azenta performed Sanger DNA sequencing for all plasmids constructed throughout this study. LC-MS/MS analyses (including Hi-resolution analysis) are performed on an Agilent 6530 Accurate-Mass Q-TOF MS equipped with a dual electrospray ionization source and an Agilent 1260 LC system with diode array detector, or an Agilent 1290 Infinity II HPLC System interfaced with an Agilent 6460C Triple Quadrupole LC/MS system. MS and UV data are analyzed with Agilent MassHunter Qualitative Analysis version B.05.00. Human and mouse integrins and biotinylated integrins were purchased from R&D Systems (Minneapolis, MN) or Acro Biosystems (Newark, DE). LAP protein was purchased from R&D Systems. Preparative HPLC was carried out using an Agilent 218 purification system (ChemStation software, Agilent) equipped with a ProStar 410 automatic injector, Agilent ProStar UV-Vis Dual Wavelength Detector, a 440-LC fraction collector and preparative HPLC column indicated below. Semi-preparative HPLC purifications were performed on an Agilent 1200 Series Instrument with a multiple wavelength detector and Phenomenex Luna 5µm C8(2) 250 x100 mm semi preparative column. NMR data are acquired using a 400 MHz or a 600 MHz Bruker Avance III spectrometer with a 1.7 mm cryoprobe. All signals are reported in ppm with the internal DMSO-d6 signal at 2.50 ppm (^1^H-NMR) or 39.52 ppm (^13^C-NMR). 1D data is reported as s = singlet, d = doublet, t = triplet, q=quadruplet, m = multiplet or unresolved, br = broad signal, coupling constant(s) in Hz.

Cell culture and fermentation experiments used one of the following media: (i) M9 minimal medium [17.1 g/L Na2HPO4· 12 H2O, 3 g/L KH2PO4, 0.5 g/L NaCl, 1 g/L NH4Cl, 1 mL/L MgSO4 solution (2 M), 0.2 mL/L CaCl2 solution (0.5 M), pH 7.0; after autoclaving, 10 mL/L sterilized glucose solution (40% w/v), 10 mL/L trace metals, and 10 mL/L standard vitamin mix - for trace metals solution, 27 g/L of FeCl3 · 6H2O, 2 g/L of ZnCl2 · 4H2O, 2 g/L of CaCl2 · 6H2O, 2 g/L of Na2MoO4 · 2H2O, 1.9 g/L of CuSO4 · 5H2O, 0.5 g/L of H3BO3, and 100 ml of concentrated HCl; and for vitamin solution, 0.42 g/L of riboflavin, 5.4 g/L of pantothenic acid, 6 g/L of niacin, 1.4 g/L of pyridoxine, 0.06 g/L of biotin, and 0.04 g/L of folic acid], (ii) Luria-Bertani (LB) medium [10 g/L casein peptone, 5 g/L yeast extract, 10g/L NaCl, pH 7.0], or (iii) terrific broth (TB) medium [12 g/L casein peptone, 24 g/L yeast extract, 2.2 g/L KH2PO4, 9.4 g/L K2HPO4, 4 mL/L glycerol, pH 7.0].

#### Molecular Biology and Cloning

All lasso peptides derived from MccJ25 were cloned via type IIs restriction-guided assembly of synthetic lasso precursor sequences (derived from the complete MccJ25 operon deposited in GenBank under the accession no: AF061787.1) into a receiver expression vector. The receiver vector (pLAM106 sequence, “A plasmid”; Supplementary **Figure S2**) was constructed via overlap assembly (NEBuilder HiFi DNA Assembly, New England Biolabs) from DNA fragments encoding a P15a origin, chloramphenicol resistance cassette CmR, a *sacB* counterselection marker (*Bacillus subtilis*), and p119 promoter from the Anderson promoter collection (Identifier: BBa_J23119, iGEM Registry of Standard Biological Parts). Correct assembly following transformation into *E. coli* DH5α (New England Biolabs) was validated via Sanger sequencing. Lasso peptide precursor sequences with flanking BsaI sites and relevant overhang sequences were synthesized as eBlocks (Integrated DNA Technologies) and directly amplified using primers oLAM59 and oLAM60. Purified PCR products were used in type-IIs restriction/ligation assembly in a 15 µL reaction mix with pLAM106, T4 Ligase, 1X T4 Ligase Buffer, BsaI-HF v2, 1X NEB rCutsmart Buffer. Initial 20 min digestion at 37 °C was followed by 15 cycles of alternating 37 °C incubation (10 min) and 16 °C incubation (10 min). Reactions were then held at 16 °C indefinitely prior to transformation into chemically competent DH5α *E. coli* (New England Biolabs) cells. Completed transformations were plated on LB media containing the appropriate antibiotics and 10% sucrose to select against undigested pLAM106 plasmids retaining the *sacB* cassette. Lasso peptide precursor peptide variants derived from *mcjA* were assembled into pLAM106 are referred to herein as “A-plasmids.” Single-site and multi-site mutations involving *mcjA* in this study were performed using blunt end ligation or QuikChange site-directed mutagenesis or 2-site NNK mutagenesis. Supplementary **Table S5** provides DNA sequences for plasmid construction used for MccJ25 variant production. Primers used for 2-site NNK mutagenesis at A3 and I16 positions of lassotide **14** are provided in Supplementary **Table S6**.

The MccJ25 biosynthetic operon expression vector (pLAM58 sequence, “B-plasmid”, Supplementary **Figure S3**) containing the biosynthetic enzymes and transporter was built from synthetic DNA fragments. Briefly, the genes encoding *mcjB* (peptidase, WP_097313465.1), *mcjC* (lasso cyclase, WP_256498469.1), and *mcjD* (transporter, WP_097313467.1) were synthesized together with their native promoter (Twist Biosciences). The genes and their promoter were then assembled into a pBR322 backbone containing a kanamycin (*kanR*) resistance marker via overlap assembly (NEBuilder HiFi DNA Assembly, New England Biolabs). Correct assembly following transformation into *E. coli* DH5α (New England Biolabs) was validated via Sanger sequencing.

### Lasso peptide production

#### *E. coli* culturing media

For all transformations, starter cultures, and precultures, Luria-Bertani (LB) solid [10 g/L casein peptone, 5 g/L yeast extract, 10g/L NaCl, 15 g/L agar, pH 7.0] and liquid [10 g/L casein peptone, 5 g/L yeast extract, 10 g/L NaCl, pH 7.0] media were used. For production cultures, supplemented M9 liquid (3 g/L KH2PO4, 12.8 g/L Na2HPO4.7H2O, 0.5 g/L NaCl, 1 g/L NH4Cl, 4 mL/L glycerol, 2 g/L casamino acids, 1 mL/L Teknova® vitamin mix, 0.49 g/L MgSO4, 0.015 g/L CaCl2) medium was used.

#### Lasso “B” plasmid transformation in chemical competent BL21 E coli cells

A 5 mL overnight preculture of chemically competent BL21(DE3) *E. coli* cells was inoculated with purified “B” plasmid. After 14-16 hours of shaking at 37°C and 200 rpm, 0.3 mL of this *E. coli* starter culture (1:100 dilution) was added to 30 mL of LB media and grown in a shaker at 37°C and 200 rpm. When the OD600 reached 0.4-0.5 (rough time estimate of 3 to 3.5 hours), the cells were immediately placed on ice. Cultures were centrifuged using 30 mL ice cold centrifuge tubes. Cells were harvested by centrifugation at 5000 x g for 9 minutes at 4°C. This process was repeated until all cultures had been centrifuged. Supernatants were decanted and pellets washed with 30 mL of 100 mM MgCl2. Cells were centrifuged again at 5000 x g for 9 minutes at 4°C. The cell pellets were washed with 30 mL of 100 mM CaCl2. Centrifugation was repeated and supernatant was decanted. Pellets were washed with 30 mL of 85 mM CaCl2, 15% glycerol. Centrifugation was repeated and supernatant decanted. Pellets were resuspended in 1.5 mL of 85 mM CaCl2, 15% glycerol and 50 µL aliquots of chemically competent BL21(DE3) E coli cells containing the Lasso “B” plasmid cells were transferred into 1.5 ml individual Eppendorf tubes which were pre-autoclaved and prechilled. Tubes were snap-frozen tubes after aliquoting using 100% ethanol and dry ice mixture.

#### Transformation of *E. coli* with “A” plasmids to create lasso production strains

Chemically competent BL21(DE3) *E. coli* cells (50 µL aliquots) that contain the Lasso “B” plasmid were incubated on ice for 30 mins and 2.5 µl of a Golden Gate reaction mixture containing “A” plasmid was added to the wells of plates containing the cells. The incubation mixture was heat shocked at 42°C for 30 seconds, and the plate immediately placed on ice for 5 mins. Cells then were immediately transferred into 500 µL of pre-aliquoted LB medium and then were shaken at 37°C degrees for at least 1 hour. After centrifugation at 9000 rpm, the cell pellets were recovered and supernatant discarded (about 400 µL), cells were resuspended in LB media + Kanamycin 50 µg/ml + Chloramphenicol 17 µg/ml + 10% sucrose in wells of plates and were incubated at 37°C for 16-18 h. These *E. coli* production strains were stored at -80°C as glycerol stocks (500 µL of the LB culture from a single colony well with 500 µL of a 50% glycerol solution transferred to a 2.0 mL cryogenic vial) and used for production of engineered lasso peptides.

### Production of lassotides

#### Shake flask cultures

Precultures were generated by inoculating individual sequence verified *E. coli* colonies containing the “A” and “B” plasmids for each engineered lasso peptide from either a patch plate or a premade glycerol stock into 10 mL of LB broth in culture tubes with 50 mg/mL of kanamycin and 17 mg/mL of chloramphenicol and shaking overnight at 37°C, 200 rpm. Precultures reached OD600 of 2 to 3 after 16 hours incubation. Subsequently, 5 mL of the corresponding preculture was inoculated into 125 mL of LB broth in 250 mL baffled flasks with 50 mg/mL of kanamycin and 17 mg/mL of chloramphenicol to reach OD600 ≥ 0.1. The pre-cultures were then allowed to grow for 4-5 hours at 37°C, 200 rpm to reach target OD between 2 to 3. These final pre-cultures (125 mL) then were inoculated into 1 L of M9 media supplemented with 50 mg/mL of kanamycin and 17 mg/mL of chloramphenicol in a 2 L baffled shake flask. Production cultures were grown for 3 days at 30°C, 200 rpm, after which lasso titer was assessed and engineered lasso peptides were isolated and purified as described below.

#### Lassotide isolation from cultures

Whole-cell broth was harvested in 750 mL Nalgene bottles via centrifugation at 5,000 x g, and the resulting supernatant was subjected to solid phase extraction (SPE). SPE columns were prepared as follows: For every liter of culture volume, 12.5 g of Diaion^®^ HP20SS polystyrene absorbent resin (Itochu Chemicals America, White Plains, NY) was packed into an empty fritted column. The column was washed with 5 column volumes of MeOH, then equilibrated with 5 column volumes of deionized water. The supernatants were loaded directly onto a prepared HP20ss column and fractionated as follows: The column was washed with 125 mL deionized water, then eluted with 125 mL 30% MeOH/water, 125 mL 50% MeOH/water, 125 mL 75% MeOH/water, and 125 mL 100% MeOH. Each fraction was run on the LC-MS to determine which contained the lasso peptide and approximate the quantity. Fractions containing lasso peptides were pooled and concentrated to dryness on a rotary evaporator and lyophilizer. These fractions were then further purified by semi-preparative HPLC.

#### Semi-preparative HPLC purification

(1-10 mg scale). Semi-preparative HPLC purifications were performed on an Agilent 1100 Series Instrument with a multiple wavelength detector. The semipreparative HPLC method included the following: column: Phenomenex Luna 5 µm C18(2) 250 mm x 10 mm, flow rate: 4 mL/min, temperature: RT, mobile phases: water, MeOH, acetonitrile, trifluoroacetic acid, in different percentages and used as gradients, injection volume: 0.01 to 1.0 mL. All lassotides were purified to ≥ 97% purity levels. HPLC traces for representative purified lasso peptides are shown in Supplementary **Figure S7**.

#### Production in fermenters

(Lassotide **35).** Seed cultures were generated by inoculating a 250 mL baffled flask with 50 mL LB medium [0.5 g tryptone, 0.25 g yeast extract, 0.5 g NaCl, 47.5 mL DI water] and subsequently adding 0.05 mL of 50 mg/mL kanamycin, 0.025 mL of 34 mg/mL chloramphenicol, and 0.5 mL of production strain glycerol stock. Cultures were grown for 5 hours at 37°C and 200 rpm, reaching OD600 of ca. 1.5. Precultures were prepared by inoculating a 1 L baffled flask with 2 mL of seed culture into 200 mL of HDF media [0.17 g citric acid, 0.016 g concentrated NH4OH, 0.4 g Na2SO4, 0.62 g (NH4)2SO4, 2.92 g K2HSO4, 0.64 g NaH2PO4-H2O, 0.02 g Sigma 204 (antifoam), 180 mL DI water] together with 14 mL glucose (600 g/L), 0.8 mL Teknova vitamin stock and trace elements, 0.2 mL thiamine-HCl (100 g/L), 0.82 mL MgSO4 (400 g/L), 0.20 mL kanamycin (50 mg/mL), and 0.1 mL chloramphenicol (34 mg/mL) in a 1L baffled flask and pH was adjusted to 6.0. Cultures were grown for 10 hours at 37°C and 200 rpm, reaching OD600 of 3.0-5.0. Four fed-batch stirred tank reactors (5 L working volume) bearing Rushton impellers (7 ¼ inch and 10 ¼ inch from the lid) were charged with 3.0 L of HFD media, followed by 208 mL glucose (600 g/L), 12 mL Teknova vitamin stock and trace elements, 3.0 mL thiamine-HCl (100 g/L), 12.2 mL MgSO4 (400 g/L), 3.0 mL kanamycin (50 mg/mL), and 1.5 mL chloramphenicol (34 mg/mL), and then inoculated with 30 mL (1%) of the preculture. Fermenters were fed with HFD medium using a feed strategy controlled by dissolved oxygen measurements with a set point of 20% at air flow of 1.5-3.0 L/min and agitation set at 900 rpm. Temperature was set at 30°C and pH was maintained at 6.8 through the addition of 28% NH4OH or 4N H2SO4. Fermentations were monitored over time wherein the mean maximum cell density (OD600 = 140) and titers of 5.39 g/L, 5.64 g/L, 5.70 g/L, and 5.95 g/L for **35** were achieved over 72 h across the 4 tanks, leading to a mean titer of 5.67 ± 0.10 g/L (SEM). Scale-up production of all other lassotides in fermenters, including **36** and **38**, followed the same general protocol.

#### Lassotide recovery from fermenters

A simple process for downstream recovery was developed for all secreted lassotides produced in fermenters that involved: (i) addition of Diaion^®^ HP20SS absorption resin (Itochu Chemicals; 33 g/L whole cell broth), (ii) stirring for 4 h at 30°C, (iii) filtration through a nylon mesh filtration apparatus, (iv) washing the retentate with MeOH to desorb the lassotides, and (v) concentration to crude product. Lassotides were purified by preparative HPLC.

#### Preparative HPLC purification

(10 mg-30 g scale). Preparative HPLC was carried out using a Shimadzu LC 8A HPLC equipped with a Vydac 10-15 µm C18, 300 mm x 47 mm, preparative column (0.1-30 g scale), or an Agilent 1100 or 1260 purification system (ChemStation software, Agilent) equipped with an autosampler, multiple wavelength detector, Prep-LC fraction collector and Phenomenex Luna 5 µm C18(2) 150 x 30 mm preparative column (10-100 mg scale). The 1260 system additionally included an Agilent MSD. The preparative HPLC method for the pooled lasso peptide containing fractions (100 mg scale) included the following: Column: Phenomenex Luna® preparative column 5 µM, C18(2) 100 Å 150 mm x 30 mm, Flow rate: 20 mL/min, Temperature: RT, Mobile Phases: HPLC grade water, MeOH, acetonitrile, isopropyl alcohol, trifluoroacetic acid (TFA), in different percentages and used as gradients, Injection amount: variable 5-200 µL. Example method for lassotide **35**: Solvent A, water with 0.05% TFA; solvent B, acetonitrile with 0.05% TFA. 24% - 26% B over 20.0 min, then 20% to 95% B over 1 min followed by 95% B for 3 min. 5-min post-run equilibration time. Analytical HPLC traces for large scale purifications of lassotides **35**, **36**, **38**, and **47** are shown in Supplementary **Figure S8**.

#### Purity QC

The purity of each purified lassotide sample of lasso peptide purified by semi-preparative or preparative are confirmed by HPLC using (1) an Agilent 6460C Triple Quadrupole LC/MS/MS system (LC/TQ) equipped with a Jet stream source (AJS), an Agilent 1290 Infinity II LC system, and a diode array detector (DAD) or (2) a Shimadzu LC-10ADvp equipped with UV-vis detector (210-215 nm detection for peptides) and a Vydac 5 µm C18 column, 250 mm x 4.6 mm. Where possible, MS/MS fragmentation was used to further characterize lasso peptides based on the fragmentation rules described in Fouque et al.,^76^ and to confirm amino acid sequences. High-resolution LCMS data were acquired, as described above, to further confirm peptide structures. The analytical LCMS method for purity assessment was as follows: 1 µL samples were injected onto a Phenomenex Kinetex XB-C18 column (1.7 µm, 50 x 2.1 mm) operating at 0.4 mL/min with 1 minute at 0% acetonitrile (+ 0.1% (v/v) formic acid), a gradient of 0% to 40% over 8 minutes, then 40% to 100% acetonitrile from 8 minutes to 10 minutes followed by a 4-minute hold at 100%. Analytical samples were preceded by four blank injections, with the 4th blank used for background subtraction. Column: Phenomenex Kinetex 1.7 μm XB-C18 100 A, 50 mm x 2.1 mm, Flow rate: 0.4 mL/min, Temperature: 40 °C, mobile phase A: 0.1% formic acid in water (LCMS grade), mobile Phase B: acetonitrile (LCMS grade), Injection volume: 1 µL, gradient: 5% B for 1.0 min, then 5 to 50% B over 7 min followed by 50 to 95% B over 2 min and 95% B for 2 min. 2 min post run equilibration time. After a blank subtraction, the peaks from the TIC, UV210, and UV280 signals were integrated, and purity was reported as the area sum % for each signal using Agilent MassHunter Qualitative Analysis 10.0.

### Preparation of conjugation intermediate Acyl 50

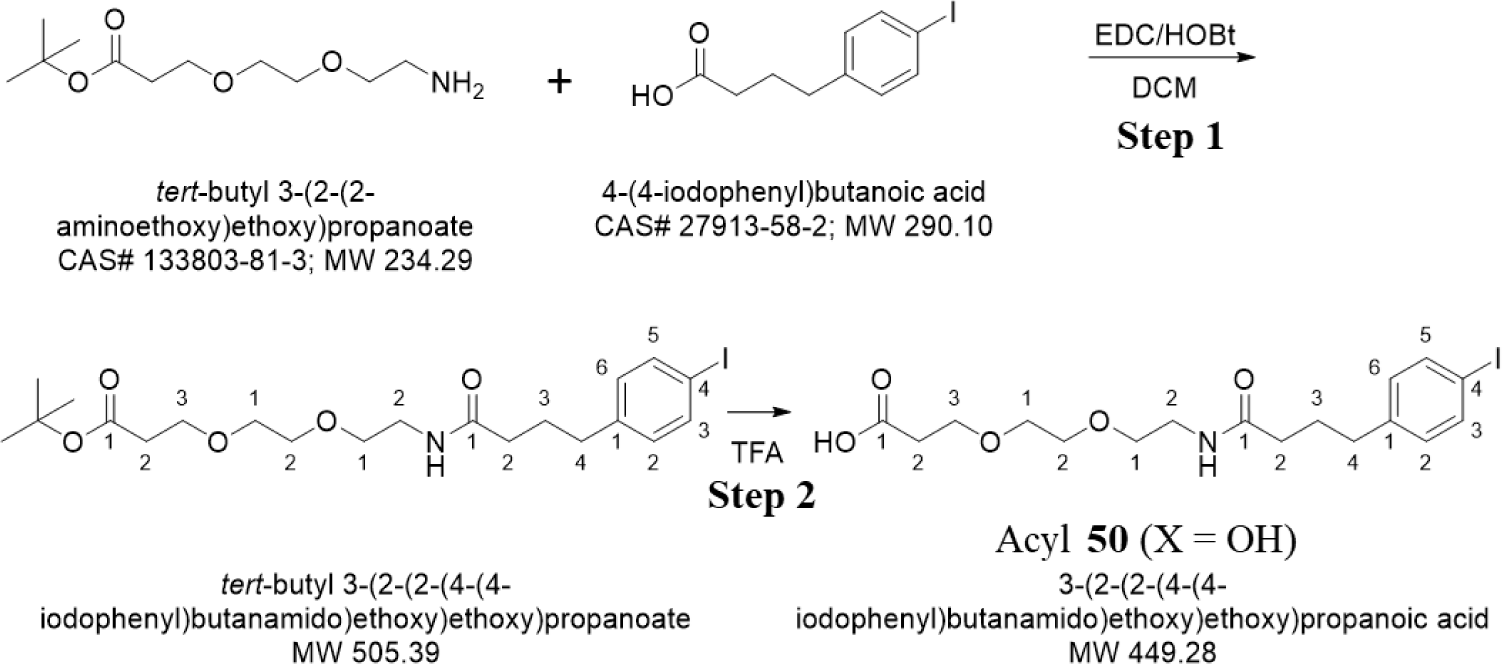

#### Step 1

Synthesis of *tert*-butyl 3-(2-(2-(4-(4-iodophenyl)butanamido)ethoxy)ethoxy)propanoate 4-(4-iodophenyl)butanoic acid (500 mg, 1.72 mmol; Combi Blocks: Cat # QC-8160), EDC (496 mg, 2.59 mmol), and HOBt (396 mg, 2.59 mmol) were mixed in 15 mL of dichloromethane and stirred for 15 minutes. tert-butyl 3-(2-(2-aminoethoxy)ethoxy)propanoate (485 mg, 2.07 mmol; BroadPharm: Catalog # BP-20523) was added and stirred for 1 hour. The solution was washed with 1M HCl (3 x 10 mL), saturated Na2CO3 (3 x 10 mL), and brine (10 mL), dried over Na2SO4 and filtered. The solvent of the filtrate was removed *in vaccu* to get the crude product as an oil. The crude product was purified with flash chromatography with 1-5% MeOH in DCM to obtain the final product, *tert*-butyl 3-(2-(2-(4-(4-iodophenyl)butanamido)ethoxy)ethoxy)propanoate (770 mg, 1.52 mmol) as a pale yellow oil in 88.4% yield. Analytical HPLC purity chromatogram, LCMS data (*m/z* = 505.7), and the ^1^H NMR spectrum are shown in **Figure S9A-C**.

#### Step 2

Synthesis of 3-(2-(2-(4-(4-iodophenyl)butanamido)ethoxy)ethoxy)propanoic acid (Acyl **50**)

To a solution of tert-butyl 3-(2-(2-(4-(4-iodophenyl)butanamido)ethoxy)ethoxy)propanoate (970 mg, 1.92 mmol) in dichloromethane (5 mL) was added trifluoracetic acid (5 mL) at room temperature. The solution was stirred for 15 minutes and the solvents were removed in vacuo to afford the product, 3-(2-(2-(4-(4-iodophenyl)butanamido)ethoxy)ethoxy)propanoic acid (Acyl **50**, 845 mg, 1.88 mmol) as a pale yellow oil in 98% yield. LCMS data (*m/z* = 449.4) and the ^1^H NMR spectrum for Acyl **50** are provided in **Figures S10A** and **S10B**.

### Preparation of lassotide 47

To a solution of lassotide **36** (438 mg, 0.192 mmol) and Et3N (402 μL, 2.88 mmol) in DMF (9.5 mL) was added a solution of 3-(2-(2-(4-(4-iodophenyl)butanamido)ethoxy)ethoxy)propanoic acid (Acyl **50**, 104 mg, 0.192 mmol), Et3N (134 μL, 0.961 mmol), and PyBop (100 mg, 0.192 mmol) in DMF (9.5 mL). The mixture was stirred for 4 hours at room temperature. The reaction was quenched with 380 mL of water with 0.1% TFA. The crude was purified by C18 flash chromatography first and further purified by HPLC to get for final product (306 mg, 0.113 mmol), 58.7% yield. See **Figure S8D** for analytical HPLC showing >98% purity. Exact Mass *m/z* = 2709.2241.

### Integrin inhibition assays

Unless otherwise noted below, all integrin inhibition and selectivity data for engineered lasso peptides shown in **Tables 1**-**4** and **Figure 3** were determined using standard ELISA assays. Human integrins αvβ1, αvβ3, αvβ5, αvβ6, αvβ8, α5β1, and αIIbβ3 were purchased from R&D Systems (Minneapolis, MN) or Acro Biosystems (Newark, DE, USA). Human integrin α8β1 and mouse integrins αvβ6 and αvβ8 were purchased from Acro Biosystems. Vitronectin, fibrinogen, and fibronectin were purchased from Sigma Aldrich (St. Louis, MO, USA). Nephronectin and LAP protein were purchased from R&D Systems. For the integrins αvβ1, αvβ3, αvβ5, αvβ6 and αvβ8, binding was visualized using a mouse anti-human integrin αv (CD51) monoclonal antibody targeting the αv subunit (cat. no. MAB1219 was purchased from R&D Systems). For integrin α5β1, binding was visualized using a mouse anti-human integrin a5 (CD49e) monoclonal antibody that targets the α5 subunit (cat. no. MAB1864-SP was purchased from R&D Systems). For integrin α8β1, binding was visualized using a mouse anti-human integrin α8 monoclonal antibody that targets the α8 subunit (cat. no. MAB6194-SP was purchased from R&D Systems). For integrin αIIbβ3, binding was visualized using a mouse anti-human integrin αIIb (CD41a) monoclonal antibody that targets the αIIb subunit (cat. No. 16-0411-85 purchased from ThermoFisher Scientific). In all cases, anti-mouse IgG-peroxidase from Sigma Aldrich (Sigma, cat. no. DC02L) was used as secondary antibody, containing a peroxidase conjugate that is employed for the visualization and quantification. Peroxidase development is performed by using the supplier’s recommended methods with the substrate 3,3,5,5’-tetramethylethylenediamine (TMEDA) purchased from Sigma Aldrich and by adding 3M H2SO4 to stop the reaction. The absorbance (450, 492 nm) is recorded with a Victor Nivo plate reader (Perkin Elmer, San Jose, CA). Every concentration is analyzed in triplicate and the resulting inhibition curves are analyzed using GraphPad Prism software. The inflection point of these curves describes the IC50 value. Each plate also contained either cilengitide or MK-0429 as reference compounds, except for assays involving αIIbβ3, which used tirofiban (positive control compounds were purchased from MedChemExpress, Monmouth Junction, NJ, USA). Blocking and binding steps are performed with TS buffer (20 mM Tris-HCl pH 7.5, 150 mM NaCl, 1 mM CaCl2, 1 mM MgCl2, and 1 mM MnCl2) containing 1% BSA. After the incubation time, washing steps are conducted with PBST buffer (10 mM Na2HPO4, pH 7.5, 150 mM NaCl, and 0.01% Tween 20).

For αvβ3 and αvβ5 assays, a clear flat-bottom 96-well ELISA plate (from Sigma Aldrich) was coated overnight with 100 µL of vitronectin (2 µg/mL) in carbonate buffer (15 mM Na2CO3, 35 mM NaHCO3, pH 9.6) at a temperature of 4°C. After removing the solutions from the plate, the wells are blocked for 1 h at room temperature with 150 μL TSB buffer per well. The plate is subsequently washed three times with 200 μL PBST buffer per well. Afterward, 4.0 μg/mL of the soluble integrin αvβ3, or 1 μg/mL of the soluble integrin αvβ5 premixed with an engineered lasso peptide (serial dilutions 10 mM to 0.001 nM), or with cilengitide or MK-0429 as a positive control compound, and the mixture was incubated in separate ECM-coated wells for 2 h at room temperature. Each concentration of inhibitor was performed in duplicate. After washing three times, the each well of the plate was treated with 100 μL of the primary anti-av antibody (MAB1219, diluted 1:500 in TSB buffer, 1.0 μg/mL) and 100 μL of the secondary antibody (anti-mouse IgG-peroxidase, diluted 1:385 in TSB buffer, 2.0 μg/mL) per well and incubated for 1 h at room temperature. After this treatment, the plate was washed three times and the binding was then visualized using TMEDA. For this substrate, the oxidation reaction was carried out for 5 min and the absorbance was measured at 450 nm. IC50 was determined as explained below.

For αvβ6 and αvβ8 assays, clear flat-bottom immune-nonsterile 96-well plates were coated with 50 mL of 0.4 µg/ml LAP (R&D Systems, cat. no. 246-LP) in carbonate buffer (15 mM Na2CO3, 35 mM NaHCO3, pH 9.6) overnight at 4 °C. The plates were then washed three times with 200 mL PBST (PBS, 0.05% Tween 20) containing 1 mM MgCl2 in a Tecan Hydroflex plate washer. The plates were blocked with 150 mL TSB buffer (20 mM Tris HCl [pH 7.5], 150 mM NaCl, 1 mM CaCl2, 1 mM MgCl2, 1 mM MnCl2, 1% BSA) for 1 hour at room temperature. The plates were washed three times with 200 mL PBST containing 1 mM MgCl2. To each well, 50 mL of integrin (0.5 µg/ml αvβ6 (R&D Systems, cat. no. 3817-AV-050) or αvβ8 (R&D Systems, cat. no. 4135-AV-050)), that was preincubated with serially diluted inhibitor (from 0.001 nM to 10 µM), or with MK-0429 positive control, for at least 30 minutes in TSB buffer, was added and incubated for 1 hour at room temperature. Each concentration of inhibitor was performed in duplicate. The plates were then washed three times in 200 µl PBST containing 1 mM MgCl2. To each well, 50 mL of 0.5 µg/mL anti-αV antibody (R&D Systems, cat. no. MAB1219) was added and incubated for 1 hour at room temperature. The plates were then washed three times in 200 mL PBST containing 1 mM MgCl2. To each well, 50 mL of goat anti-mouse IgG peroxidase conjugate (Sigma, cat. no. DC02L) at concentrations of 0.03 µg/ml for αvβ6 or 0.07 µg/ml for αvβ8 was added and incubated for 1 hour at room temperature. The plates were then washed three times in 200 mL PBST containing 1 mM MgCl2. To each well, 50 mL TMB substrate (ThermoFisher, cat. no. 34021) was added and incubated for 10-30 minutes at room temperature. The reaction was terminated with 50 mL 6N H2SO4. The plate was then measured at an OD450 in a Perkin Elmer Victor Nivo multimode plate reader.

For αvβ1 assay, a clear flat-bottom 96-well ELISA plate (from Sigma Aldrich) was coated overnight with 100 µL of fibronectin (4 µg/mL) in carbonate buffer (15 mM Na2CO3, 35 mM NaHCO3, pH 9.6) at a temperature of 4°C. The plates were then washed three times with 200 mL PBST (PBS, 0.05% Tween 20) containing 1 mM MgCl2 in a Tecan Hydroflex plate washer. The plates were blocked with 150 mL TSB buffer (20 mM Tris HCl [pH 7.5], 150 mM NaCl, 1 mM CaCl2, 1 mM MgCl2, 1 mM MnCl2, 1% BSA) for 1 hour at room temperature. The plates were washed three times with 200 mL PBST containing 1 mM MgCl2. To each well, 50 mL of integrin (10 µg/ml) αvβ1 (R&D Systems, cat. no. 6579-AVB-050) that was preincubated with serially diluted inhibitor (from 0.001 nM to 10 µM) or with control positive MK-0429, for at least 30 minutes in TSB buffer, was added and incubated for 1 hour at room temperature. Each concentration of inhibitor was performed in duplicate. The plates were then washed three times in 200 µl PBST containing 1 mM MgCl2. To each well, 50 mL of 0.5 µg/mL anti-αV antibody (R&D Systems, cat. no. MAB1219) was added and incubated for 1 hour at room temperature. The plates were then washed three times in 200 mL PBST containing 1 mM MgCl2. To each well, 50 mL of goat anti-mouse IgG peroxidase conjugate (Sigma, cat. no. DC02L) at concentrations of 0.05 µg/ml and incubated for 1 hour at room temperature. The plates were then washed three times in 200 mL PBST containing 1 mM MgCl2. To each well, 50 mL TMB substrate (ThermoFisher, cat. no. 34021) was added and incubated for 10-30 minutes at room temperature. The reaction was terminated with 50 mL 6N H2SO4. The plate was then measured at an OD450 in a Perkin Elmer Victor Nivo multimode plate reader.

For α5β1 assay, a clear flat-bottom 96-well ELISA plate (from Sigma Aldrich) was coated overnight with 100 µL of fibronectin (5 µg/mL) in carbonate buffer (15 mM Na2CO3, 35 mM NaHCO3, pH 9.6) at a temperature of 4°C. The plates were then washed three times with 200 mL PBST (PBS, 0.05% Tween 20) containing 1 mM MgCl2 in a Tecan Hydroflex plate washer. The plates were blocked with 150 mL TSB buffer (20 mM Tris HCl [pH 7.5], 150 mM NaCl, 1 mM CaCl2, 1 mM MgCl2, 1 mM MnCl2, 1% BSA) for 1 hour at room temperature. The plates were washed three times with 200 mL PBST containing 1 mM MgCl2. To each well, 50 mL of integrin (0.5 µg/ml) α5β1 (R&D Systems, cat. no. 3230-A5-050) that was preincubated with serially diluted inhibitor (from 0.001 nM to 10 µM) or with control MK-0429, for at least 30 minutes in TSB buffer, was added and incubated for 1 hour at room temperature. Each concentration of inhibitor was performed in duplicate. The plates were then washed three times in 200 µl PBST containing 1 mM MgCl2. To each well, 50 mL of 0.5 µg/mL anti-α5 antibody (R&D Systems, cat. no. MAB1864-SP) was added and incubated for 1 hour at room temperature. The plates were then washed three times in 200 mL PBST containing 1 mM MgCl2. To each well, 50 mL of goat anti-mouse IgG peroxidase conjugate (Sigma, cat. no. DC02L) at concentrations of 0.05 µg/ml was added and incubated for 1 hour at room temperature. The plates were then washed three times in 200 mL PBST containing 1 mM MgCl2. To each well, 50 mL TMB substrate (ThermoFisher, cat. no. 34021) was added and incubated for 10-30 minutes at room temperature. The reaction was terminated with 50 mL 6N H2SO4. The plate was then measured at an OD450 in a Perkin Elmer Victor Nivo multimode plate reader.

For α8β1 assay, a clear flat-bottom 96-well ELISA plate (from Sigma Aldrich) was coated overnight with 100 µL of nephronectin (4.5 µg/mL) in carbonate buffer (15 mM Na2CO3, 35 mM NaHCO3, pH 9.6) at a temperature of 4°C. The plates were then washed three times with 200 mL PBST (PBS, 0.05% Tween 20) containing 1 mM MgCl2 in a Tecan Hydroflex plate washer. The plates were blocked with 150 mL TSB buffer (20 mM Tris HCl [pH 7.5], 150 mM NaCl, 1 mM CaCl2, 1 mM MgCl2, 1 mM MnCl2, 1% BSA) for 1 hour at room temperature. The plates were washed three times with 200 mL PBST containing 1 mM MgCl2. To each well, 50 mL of integrin (4 µg/ml) α8β1 (Acro Biosystems, cat. no. IT1-H52W9) that was preincubated with serially diluted inhibitor (from 0.001 nM to 10 µM) or with control MK-0429, for at least 30 minutes in TSB buffer, was added and incubated for 1 hour at room temperature. Each concentration of inhibitor was performed in duplicate. The plates were then washed three times in 200 µl PBST containing 1 mM MgCl2. To each well, 50 mL of 0.5 µg/mL anti-α8 antibody (R&D Systems, cat. no. MAB6194-SP) was added and incubated for 1 hour at room temperature. The plates were then washed three times in 200 mL PBST containing 1 mM MgCl2. To each well, 50 mL of goat anti-mouse IgG peroxidase conjugate (Sigma, cat. no. DC02L) at concentrations of 0.05 µg/ml was added and incubated for 1 hour at room temperature. The plates were then washed three times in 200 mL PBST containing 1 mM MgCl2. To each well, 50 mL TMB substrate (ThermoFisher, cat. no. 34021) was added and incubated for 10-30 minutes at room temperature. The reaction was terminated with 50 mL 6N H2SO4. The plate was then measured at an OD450 in a Perkin Elmer Victor Nivo multimode plate reader.

For αIIbβ3 assay, a clear flat-bottom 96-well ELISA plate (from Sigma Aldrich) was coated overnight with 100 µL of fibrinogen (10 µg/mL) in carbonate buffer (15 mM Na2CO3, 35 mM NaHCO3, pH 9.6) at a temperature of 4°C. The plates were then washed three times with 200 mL PBST (PBS, 0.05% Tween 20) containing 1 mM MgCl2 in a Tecan Hydroflex plate washer. The plates were blocked with 150 mL TSB buffer (20 mM Tris HCl [pH 7.5], 150 mM NaCl, 1 mM CaCl2, 1 mM MgCl2, 1 mM MnCl2, 1% BSA) for 1 hour at room temperature. The plates were washed three times with 200 mL PBST containing 1 mM MgCl2. To each well, 50 mL of integrin (5 µg/ml) αIIbβ3 (R&D Systems, cat. no. 7148-A2-025) that was preincubated with serially diluted inhibitor (from 0.001 nM to 10 µM) or with control tirofiban, for at least 30 minutes in TSB buffer, was added and incubated for 1 hour at room temperature. Each concentration of inhibitor was performed in duplicate. The plates were then washed three times in 200 µl PBST containing 1 mM MgCl2. To each well, 50 mL of 0.5 µg/mL anti-αIIb antibody (ThermoFisher, cat. no. 16-0411-85) was added and incubated for 1 hour at room temperature. The plates were then washed three times in 200 mL PBST containing 1 mM MgCl2. To each well, 50 mL of goat anti-mouse IgG peroxidase conjugate (Sigma, cat. no. DC02L) at concentrations of 0.05 µg/ml was added and incubated for 1 hour at room temperature. The plates were then washed three times in 200 mL PBST containing 1 mM MgCl2. To each well, 50 mL TMB substrate (ThermoFisher, cat. no. 34021) was added and incubated for 10-30 minutes at room temperature. The reaction was terminated with 50 mL 6N H2SO4. The plate was then measured at an OD450 in a Perkin Elmer Victor Nivo multimode plate reader.

Integrin inhibition for lassotides **1-11**, **20**-**27**, and **33** was measured using AlphaLISA assays. This assay consisted of mixing 0.4 µg/mL biotinylated αvβ6 (Acro Biosystems, cat. no. IT6-H82E4), αvβ8 (Acro Biosystems, cat. no. IT8-H82W5) or αvβ3 (Acro Biosystems cat no. IT3-H82W9) and 0.72 µg/mL LAP with serially diluted inhibitor (from 0.001 nM to 10 µM) in binding buffer (25 mM HEPES [pH 7.4], 137 mM NaCl, 1 mM MgCl2, 1 mM MnCl2, 2 mM CaCl2, 2.7 mM KCl, and 0.05% Tween-20) in a total volume of 16 µL for 1.5-2 h in an AlphaPlate-384 plate (Perkin Elmer, cat. no., 6005350). To each well, 8 µL of 100 µg/mL anti-goat IgG acceptor beads (Perkin Elmer, cat. No. AL107C) and 3.84 µg/mL anti-human LAP (R&D Systems, cat. no. AF-246-NA) in binding buffer was added and incubated for 1 h. To each well, 8 µL of 50 µg/mL streptavidin beads (Perkin Elmer, cat. no. 67669992) was added and incubated for 45 min. The plate was then measured in a Perkin Elmer Victor Nivo multimode plate reader at excitation/emission of 680 nm/615 nm. Assays were performed in triplicate or greater (n ≥ 3).

The IC50 values for all integrin assays were calculated using non-linear regression analysis with GraphPad PRISM expressed as the concentration of inhibitor required to reduce LAP binding to the integrin by 50%.

### Plasma Stability Assays

(performed by Quintara Discovery). Human and mouse plasma (by default K2-EDTA anticoagulant) were obtained from Bioreclamation. Assays were carried out in 96-well microtiter plates. Engineered lassotides were incubated in duplicate at 37°C in the presence of human and mouse plasma. Reaction mixtures (50 μL) contained 1 μM final concentration of test lassotide added from a 10 mM DMSO stock solution. The extent of metabolism was calculated based on the disappearance of the test compound, compared to the 0-min control reaction incubations. Propantheline was included as a positive control to verify assay performance. At each of four time points (0, 0.5, 1, and 4 h), 300 μL of quench solution (50% acetonitrile, 50% methanol, and 0.05% formic acid, warmed to 37°C) containing internal standards was added to each well. Plates were sealed, vortexed, and centrifuged at 4°C for 15 minutes at 4000 rpm. The supernatant was transferred to fresh plates for analysis. All samples were analyzed on LC-MS/MS using an AB Sciex API 4000 instrument, coupled to a Shimadzu LC-20AD LC Pump system. Analytical samples were separated using Waters Atlantis T3 dC18 reverse phase HPLC column (10 mm x 2.1 mm) at a flow rate of 0.5 mL/min. Mobile phase consisted of 0.1% formic acid in water (solvent A) and 0.1% formic acid in 100% acetonitrile (solvent B). The extent of metabolism was calculated as the disappearance of the test compound, compared to the 0-min control reaction incubations. Initial rates were calculated for the compound concentration and used to determine half-life (t1/2) values.

### Hepatocyte stability

(performed by Quintara Discovery). Cryopreserved human and mouse hepatocytes (500,000 cells/mL) in DMEM were incubated with lassotides (1 mM from a 10 mM DMSO stock solution) in duplicate at 37°C. Midazolam was used as a positive control. Timepoints were collected at 0, 60, 120, and 180 min and the extent of metabolism was calculated based on the disappearance of the test compound, compared to the 0-min control reaction incubations. Plates were sealed, vortexed, and centrifuged at 4°C for 15 minutes at 4000 rpm. The supernatant was transferred to fresh plates for LC/MS/MS analysis. All samples were analyzed on LC-MS/MS using an AB Sciex API 4000 instrument, coupled to a Shimadzu LC-20AD LC Pump system. Analytical samples were separated using a Waters Atlantis T3 dC18 reverse phase HPLC column (10 mm x 2.1 mm) at a flow rate of 0.5 mL/min. The mobile phase consisted of 0.1% formic acid in water (solvent A) and 0.1% formic acid in 100% acetonitrile (solvent B). The extent of metabolism was calculated as the disappearance of the test compound, compared to the 0-min control reaction incubations. Initial rates were calculated for the compound concentration and used to determine half-life (t1/2) values and intrinsic clearance (CLint).

### Plasma protein binding

(PPB, performed by Quintara Discovery). The rapid equilibrium dialysis (RED) device, along with a Teflon base plate, (Pierce, Rockford, IL, USA) was used for the PPB binding studies. Human and mouse plasma were obtained commercially. The pH of the plasma was adjusted to 7.4 prior to the experiment. DMSO stocks (1 mM) of lassotide test articles were spiked in duplicate into plasma to make a final concentration of 2 μM. For recovery determinations, aliquots (100 μL) were transferred to a fresh 96-well deep-well plate as the time = 0 (T0) samples. An equal volume of blank PBS buffer was added to the plate to create a 50:50 plasma:buffer matrix. The T0 samples are incubated at 37°C for 4 hours. The spiked plasma solutions (300 μL) were placed into the sample chamber (indicated by the red ring); and 500 μL of PBS buffer, pH 7.4, was added to the adjacent chamber. The plate was sealed with a self-adhesive lid and incubated at 37°C on an orbital shaker (250 rpm). After 4 hours, aliquots from the RED plate (100 μL) were removed from each side of the insert (plasma and buffer) and dispensed into the 96-well plate. Subsequently, 100 μL of blank plasma was added to the buffer samples and 100 μL of blank buffer was added to all collected plasma samples. Finally, 300 μL of quench solution (50% acetonitrile, 50% methanol, and 0.05% formic acid, warmed up at 37°C) containing internal standards was added to each well. Plates were sealed, vortexed, and centrifuged at 4°C for 15 minutes at 4000 rpm. Supernatant was transferred to fresh plates for LC-MS/MS analysis. Reference compound propranolol was included in every experiment. All samples were analyzed on LC-MS/MS using an AB Sciex API 4000 instrument, coupled to a Shimadzu LC-20AD LC Pump system. Analytical samples were separated using a Waters Atlantis T3 dC18 reverse phase HPLC column (20 mm x 2.1 mm) at a flow rate of 0.5 mL/min. The mobile phase consisted of 0.1% formic acid in water (solvent A) and 0.1% formic acid in acetonitrile (solvent B). The percentage of engineered lasso peptide bound to protein is calculated by the following equations: % Free = (Concentration in buffer chamber/Concentration in plasma chamber) × 100% and % Bound = 100% - % Free. The percentage of test compound recovered was calculated by the following equation: % Recovery = (Concentration in buffer chamber*500 + Concentration in plasma chamber*300)/(Concentration in T0 sample*300) × 100%. Ratios of the peak areas of analytes over internal standard, instead of concentrations, were used for calculations. All the samples were diluted by quench solution to around 400 nM to be within compounds’ linear ranges.

### Tissue distribution

(animal study performed by Inotiv and analysis by Veloxity Labs). Lassotide **47** was dosed at 30 mg/kg by single IP injection into 6 female Balb/c mice (9 weeks of age). Three female mice were sacrificed (CO2 exposure) at each time point (3 and 8 hours post-dose) and the following tissues were surgically removed and collected for bioanalysis: brain, ovaries, breast tissue including mammary gland, lungs, colon, kidney, liver, and plasma. Collected tissues and plasma samples were shipped to Veloxity for analysis. Tissue samples were weighed to 0.01 g using analytical balance, then transferred to homogenization tube. Homogenization solution (70% isopropanol) added to each sample in a ratio of 3:1 (v/w). Homogenization beads were added to each tube. Samples were homogenized at 5,000 rpm using Omni Bead Mill homogenizer for 2 cycles at 1 min per cycle. Resultant tissue homogenate samples were extracted with isopropanol and analyzed for concentrations of lassotide **47** by LC-MS/MS (Shimazdu Nexera X2 LC-30AD linked to a Sciex 6500+ MS positive MRM mode). See Supplementary **Table S4** for full data and statistical analysis.

### Pharmacokinetic analysis

(animal study performed by Inotiv and analysis by Veloxity Labs). Pharmacokinetic (PK) assessments of lassotides 36, 47, and 52 were conducted using 3 female Balb/c mice (weighing 20 - 25 g) or 3 female Sprague Dawley rats (200 - 220 g), were obtained from Envigo (Indianapolis, IN, USA) and acclimated for at least 3 days. All animals were housed in separate cages and given ad libitum access to water and food pellets. Mice and rats were dosed with lassotides **36** and **47** at 30 mg/kg through intraperitoneal or intravenous injection of 10 mL per kg of test article formulated in 10% DMSO, 30% PEG400, and 60% water. Mice also were dosed with lassotides 47 and 52 at 10 mg/kg by intraperitoneal or subcutaneous injection. On the day of dosing and at indicated timepoints 0.083 or 0.167, 0.5, 1, 2, 4, 8, 24 hours post-dose, all mice were bled via microsampling. Lassotide **52** had an extended half-life and a 48 h timepoint was included. All timepoints, except for the last timepoint were bled via the tail vein, and the terminal bleed was via the vena cava or by cardiac puncture. Approximately 10-30 μL of whole blood is collected per timepoint. Samples will be collected in K3EDTA tubes and centrifuged under refrigerated conditions to collect plasma. The plasma samples were frozen at -80 °C until transfer to Veloxity Labs for bioanalysis by LC-MS/MS (Shimazdu Nexera X2 LC-30AD linked to a Sciex 6500+ MS positive MRM mode; Phenomenex Kinetex Biphenyl, 30×2.1mm, 2.6 µm; Mobile Phase A: 0.1% formic acid in water, Mobile Phase 2: 0.1% formic acid in acetonitrile, Wash: 50% MeOH, Flow rate 0.4 mL/min). Supplementary **Tables S3A-E** provide full PK data for individual mice and statistical analysis.

### EMT6 in vivo efficacy study 1

(performed by Reaction Biology). In vivo efficacy assessment of lassotides **36** and **47** was performed using the syngeneic orthotopic EMT6 mouse tumor model. 500,000 EMT6 cells were implanted in the mammary fat pad of female Balb/c mice and tumors were allowed to grow. Mice were randomized into six groups (n = 8) and treatment was initiated on Day 4 after implantation, when tumors had reached an average volume of 75 mm³. EMT6 tumors were treated with (i) vehicle + isotype control (clone 2A3), (ii) anti-mPD-1 (clone RMP1-14), (iii) lassotide **36** + isotype control, (iv) lassotide **47** + isotype control, (v) **36** + anti-mPD1, and (vi) **47** + anti-mPD-1. Lassotides **36** and **47** were formulated in 10% DMSO, 30% PEG400, 60% water (vehicle) and dosed at 100 mg/kg BID and QD, respectively. Isotype antibody control and anti-mPD1 were dosed at 10 mg/kg every 3-4 days over the course of the 23-day dosing schedule. Solo lassotide dosing groups included the isotype control. Animal weights were measured 3 times per week; mean animal weights increased slightly during the course of the study. Tumor volumes were measured by calipering 2 times per week after randomization. Euthanasia (CO2 exposure) was conducted at study end or when ethical abortion criteria were reached (e.g., weight loss >20%, severely impaired CNS function/movement, loss of righting reflex, excessive ulceration due to daily injections). Tumor volume data was analyzed and statistics performed using 2-way ANOVA modeling and GraphPad Prism 8.0 Software. Tumor growth curves are shown in **Figure 7**.

### EMT6 in vivo efficacy study 2

(performed by Reaction Biology). A dose response study involving lassotide **47** was performed using the EMT6 tumor model. The experimental set-up was similar to that described above in efficacy study 1, except that three groups of female Balb/c mice (n = 12) bearing orthotopic EMT6 tumors were IP dosed once daily (QD) for 23 days with **47** at 10 mg/kg, 30 mg/kg, and 100 mg/kg (each in combination with anti-mPD-1). Two additional groups were dosed with vehicle and solo anti-mPD-1, as described above. Tumor growth curves are shown in **Figure 8**.

### ID8 in vivo efficacy study

(performed by LabCorp). In vivo anti-tumor efficacy of lassotide **47** was conducted using LabCorp’s orthotopic ovarian cancer mouse model ID8-Luc-mCh-Puro.TD1 (ID8). C57BL/6 albino mice (C57BL/6BrdCrHsd-Tyrc) were purchased from Envigo, acclimatized for 3 days with a Teklad 2918.15 Rodent Diet and ad libitum water, and then implanted with 1 x 10^7^ ID8 cells in DMEM by IP injection. After 17 days of tumor growth (avg tumor volume 100 mm^3^), mice were randomized into 3 groups (n = 7) and dosed on Day 18 with vehicle + isotype control (clone 2A3, 10 mg/kg x 3), anti-mPD-1 (clone RMP1-14, 10 mg/kg x 3), and lassotide **47** (100 mg/kg QD in 10% DMSO, 30% PEG400, 60% water) in combination with anti-mPD-1 (10 mg/kg x 3). Animal weights were measured 3 times per week. Tumor volumes were measured weekly after randomization by whole body bioluminescence imaging (BLI) using a VIS Lumina S5 Imager (Perkin Elmer, Waltham, MA). Animals were anesthetized with 1-2% isoflurane gas, SC injected with D-luciferin, and images were obtained within 10 min for each measurement. Bioluminescence imaging (BLI) of luciferase-expressing tumor cell lines enables a noninvasive determination of site-localized tumor burden. The quantity of emitted light from the tumor after systemic injection of D-luciferin correlates with viable tumor burden. Treatment was terminated after 25 days of dosing and animal body weight and survival was monitored daily for an additional 25 days. Tumor growth and Kaplan Meier survival curves are shown in **Figure 9**. Median survival times for each group are indicated within the figure as results of the curve comparison and P-values were calculated for the survival curves vs the negative control (Group 1 – vehicle + isotype control) using the (Log-rank (Mantel-Cox) test. Individual tumor burden data for the **47**/anti-mPD-1 treatment group, acquired through BLI measurements across the 70-day study, is shown in Supplementary **Figure S6**.

### Data Analysis and Statistics

Standard non-compartmental analysis (NCA) and statistics involving PK data was conducted using Pheonix WinNonlin software. Graphical representation and statistical analysis of integrin inhibitor binding, stability, efficacy, and survival data was performed using GraphPad Prism 8.0, Graphpad Software, San Diego, California, USA. Differences were considered significant when P-values were <0.05. For efficacy studies, data are displayed as means +/- SEM and P-values were calculated compared to negative controls and/or the positive controls using the one-way Repeated Measures ANOVA with Bonferroni post-test. In addition, statistical analysis was performed using two-way Repeated Measures ANOVA for multiple comparisons throughout treatment days in comparison to the positive controls. For survival studies, comparison of survival curves was calculated based on Log-rank (Mantel-Cox) test using a proportional hazards model.

## Supplementary Information

### Supplementary Tables

**Table S1.**
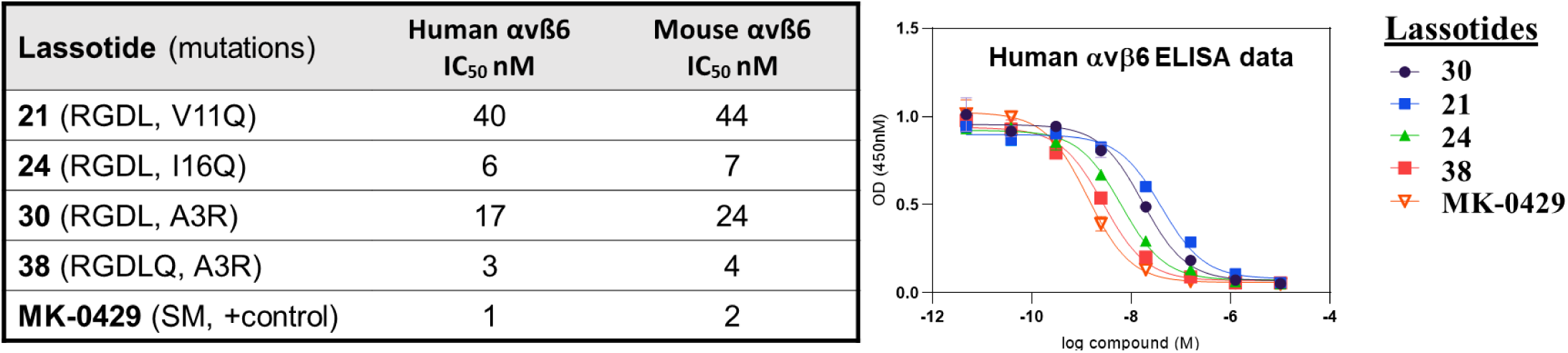
ELISA data showing lassotide integrin inhibition of human and mouse αvβ6. ELISA assays were run in triplicate and mean IC50 values are shown.

**Tabel S2A.**
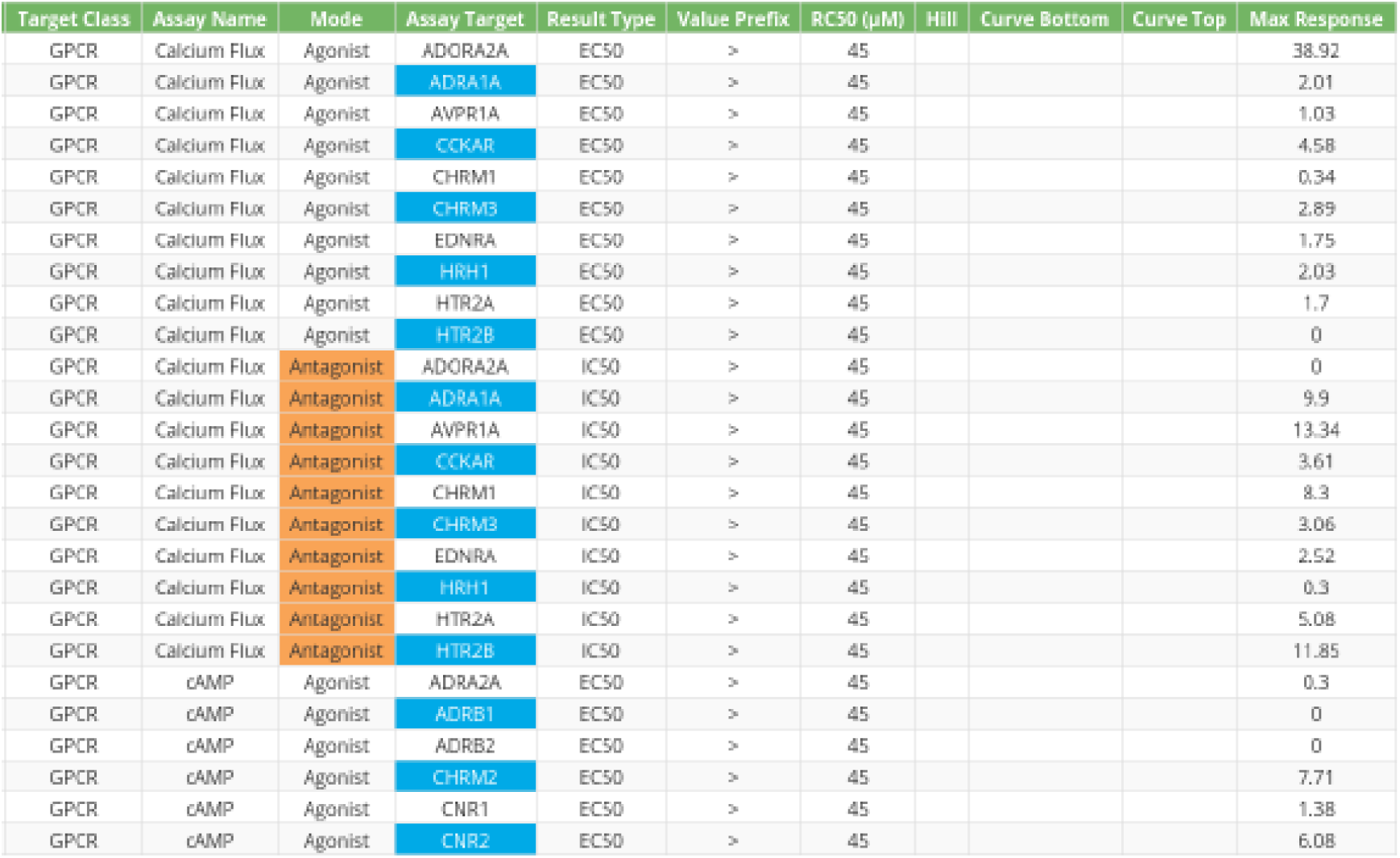
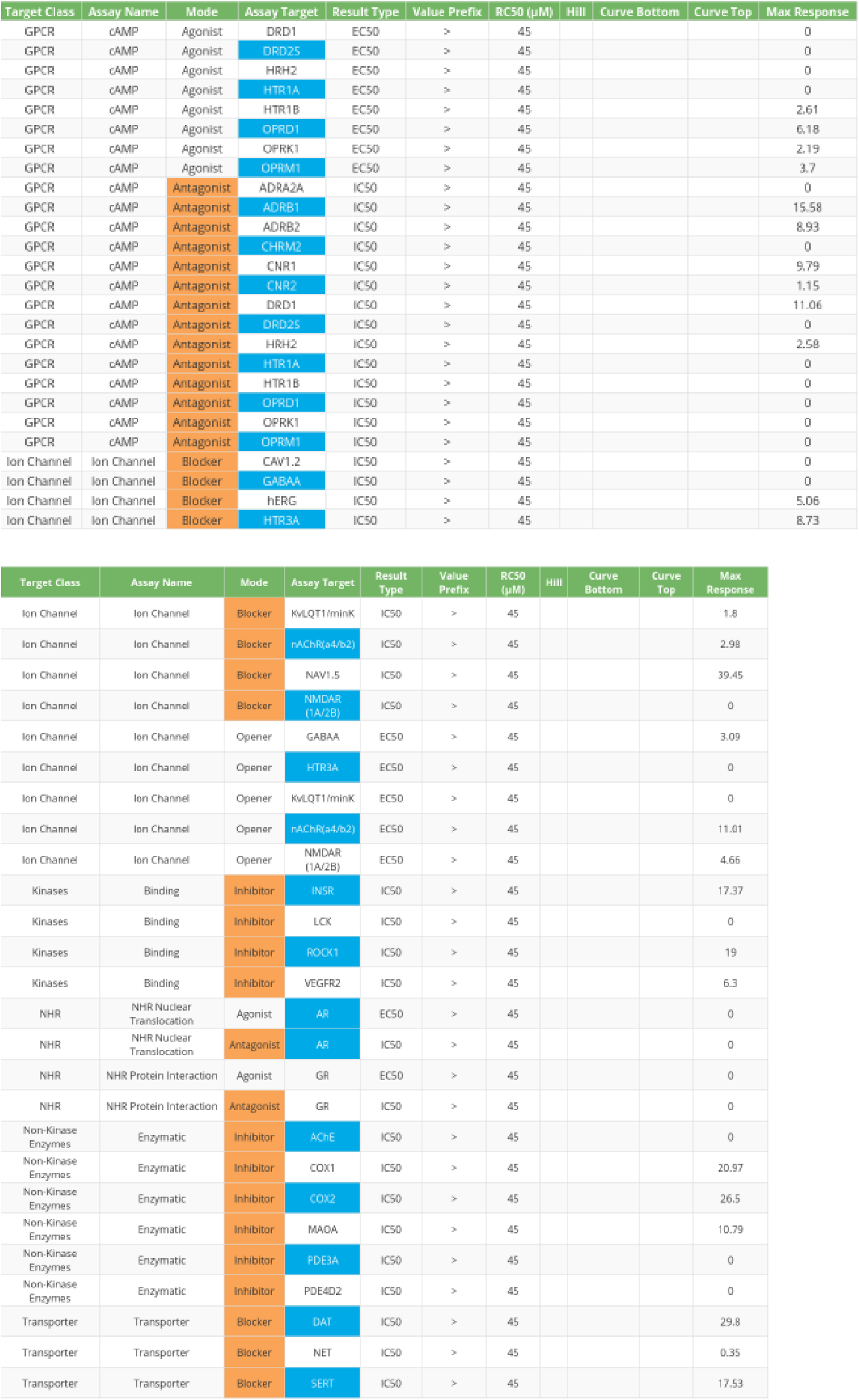
Safetyscan47 data for lassotide **36**. Responses were measured for 78 drug targets. No responses were observed up to 45 μM concentrations.

**Table S2B.**
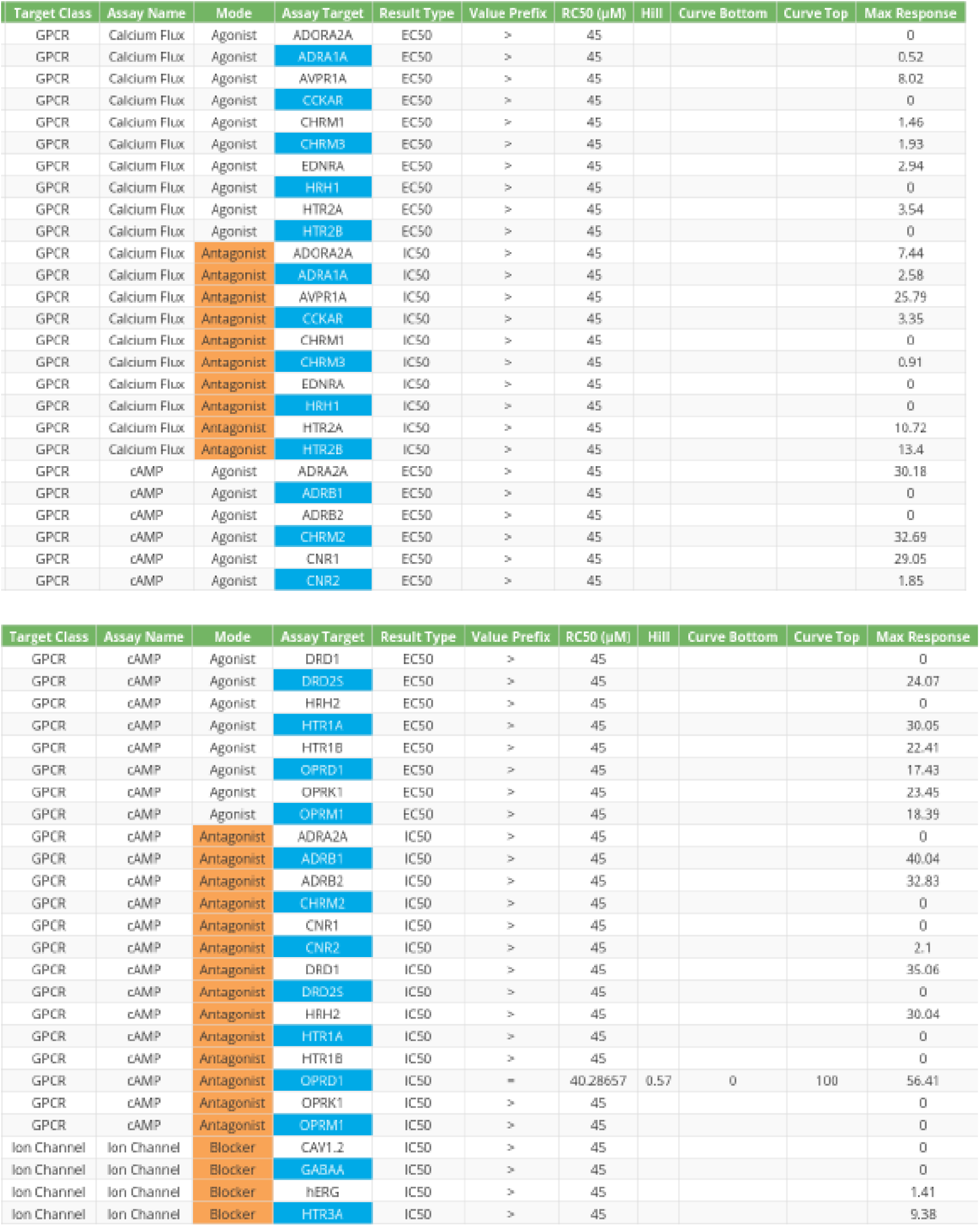
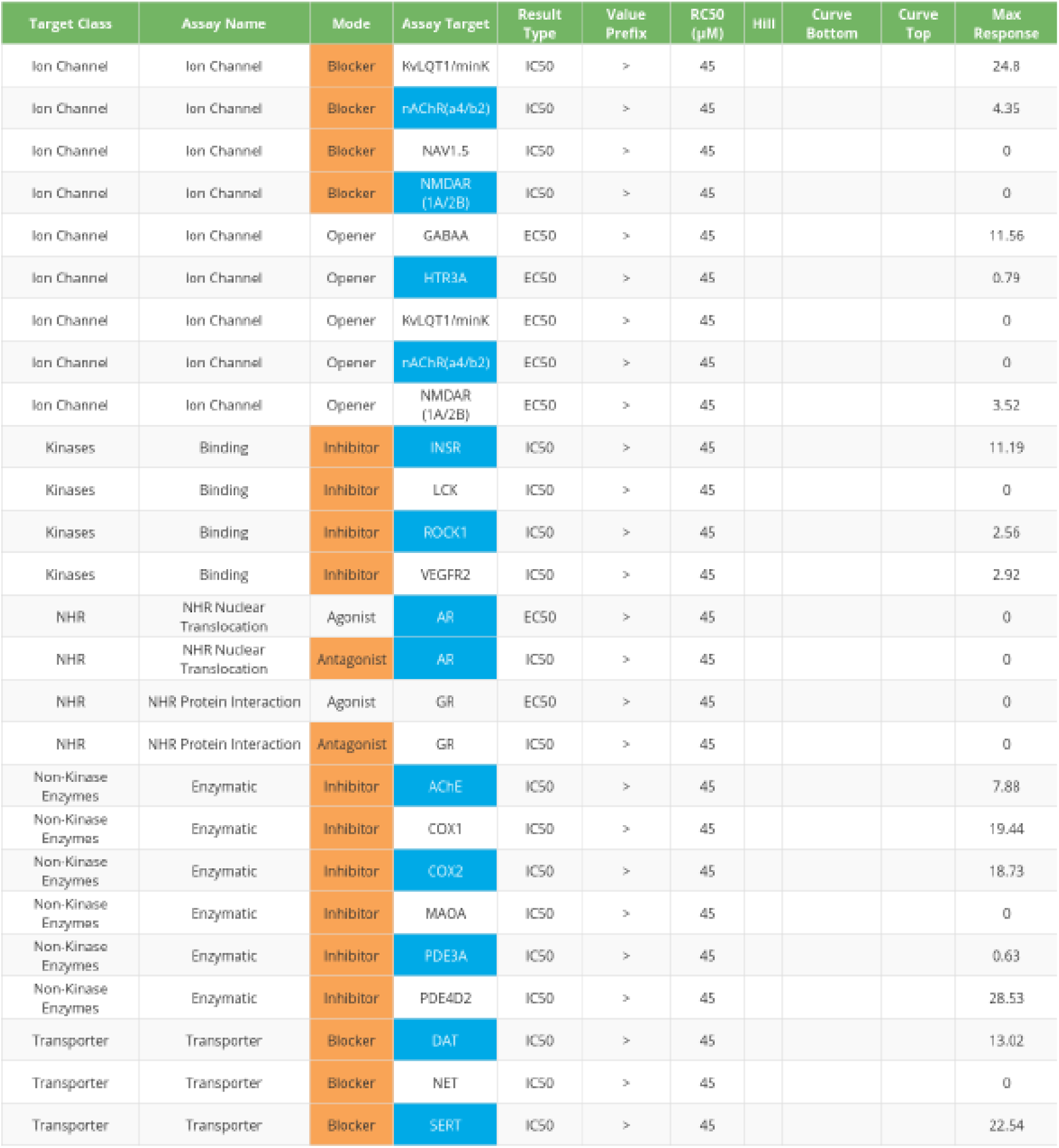
Safetyscan47 data for lassotide **47**. Responses were measured for 78 drug targets. One response was seen at 40.3 μM for GPCR receptor OPRD1.

**Table 3SA.**
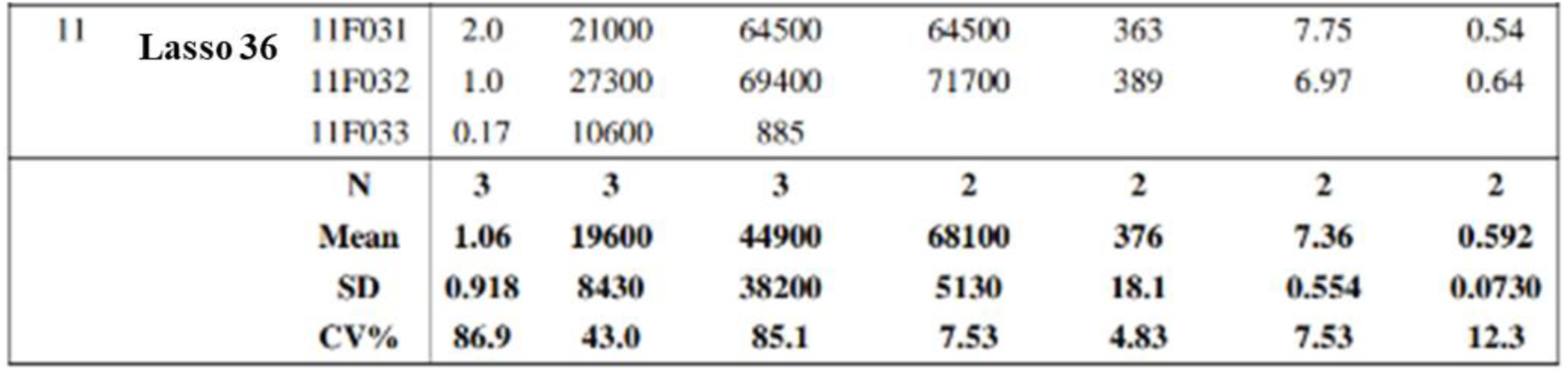
PK parameters and statistics for lassotide **36** dosed by IP administration at 30 mg/kg in 3 female Balb/c mice. Standard deviations in measurements are provided.

**Table S3B.**
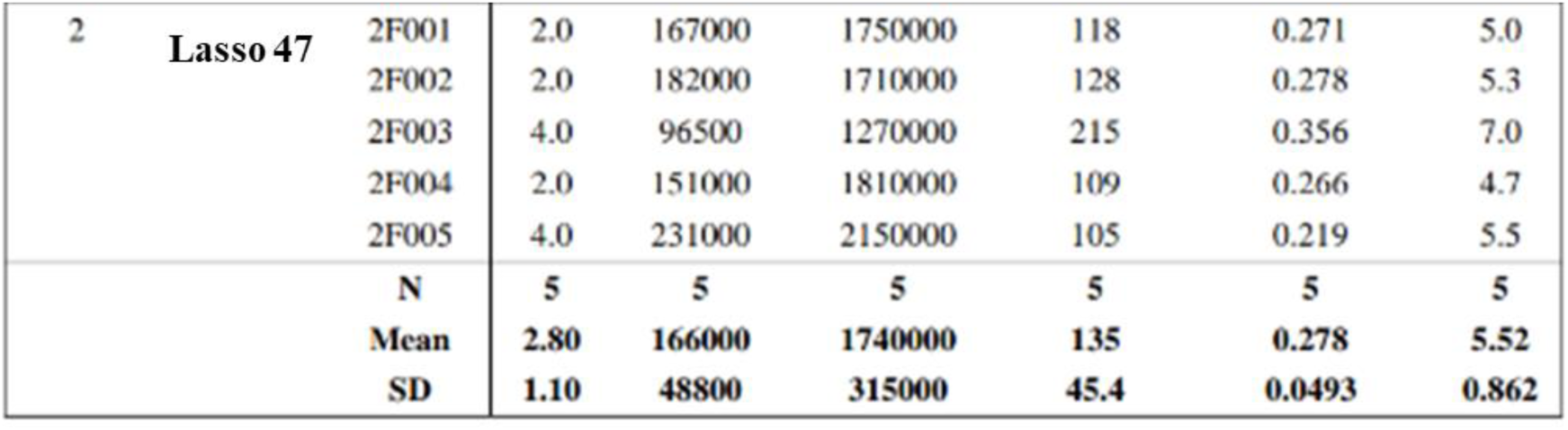
PK parameters and statistics for lassotide **47** dosed by IP administration at 30 mg/kg in 5 female Balb/c mice. Standard deviations in measurements are provided.

**Table S3C.**
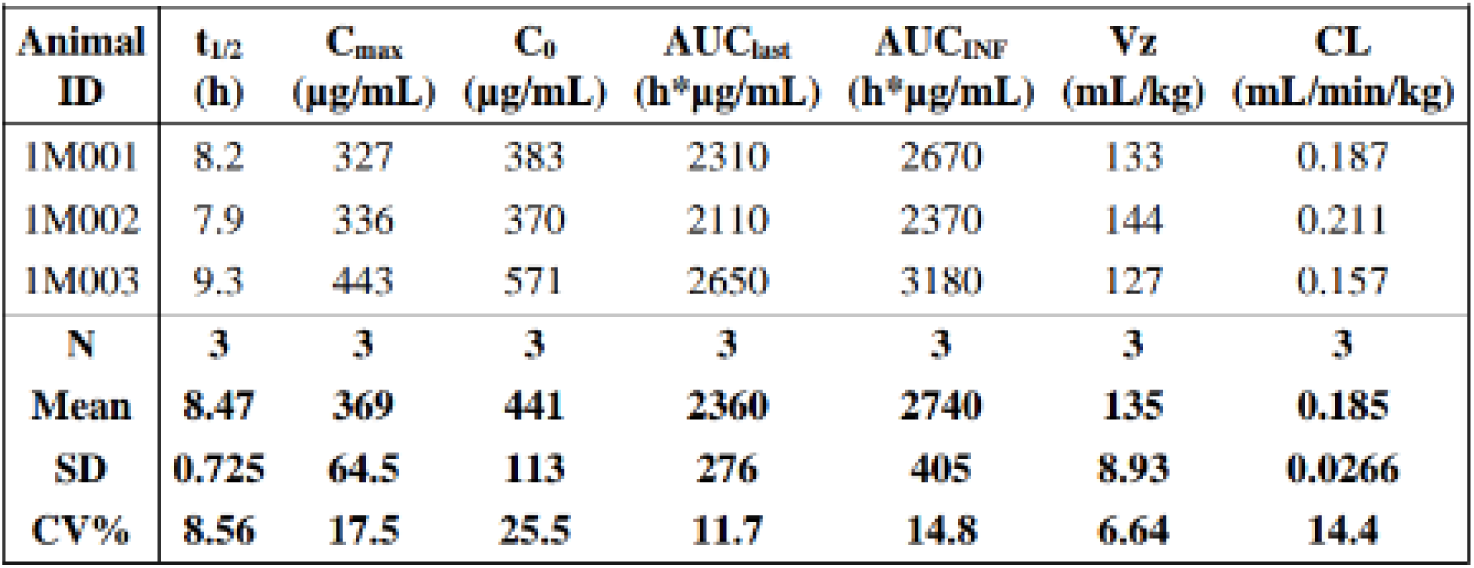
PK parameters and statistics for lassotide **47** dosed by IP administration at 30 mg/kg in 3 female Sprague-Dawley rats. Standard deviations in measurements are provided.

**Table S3D.**
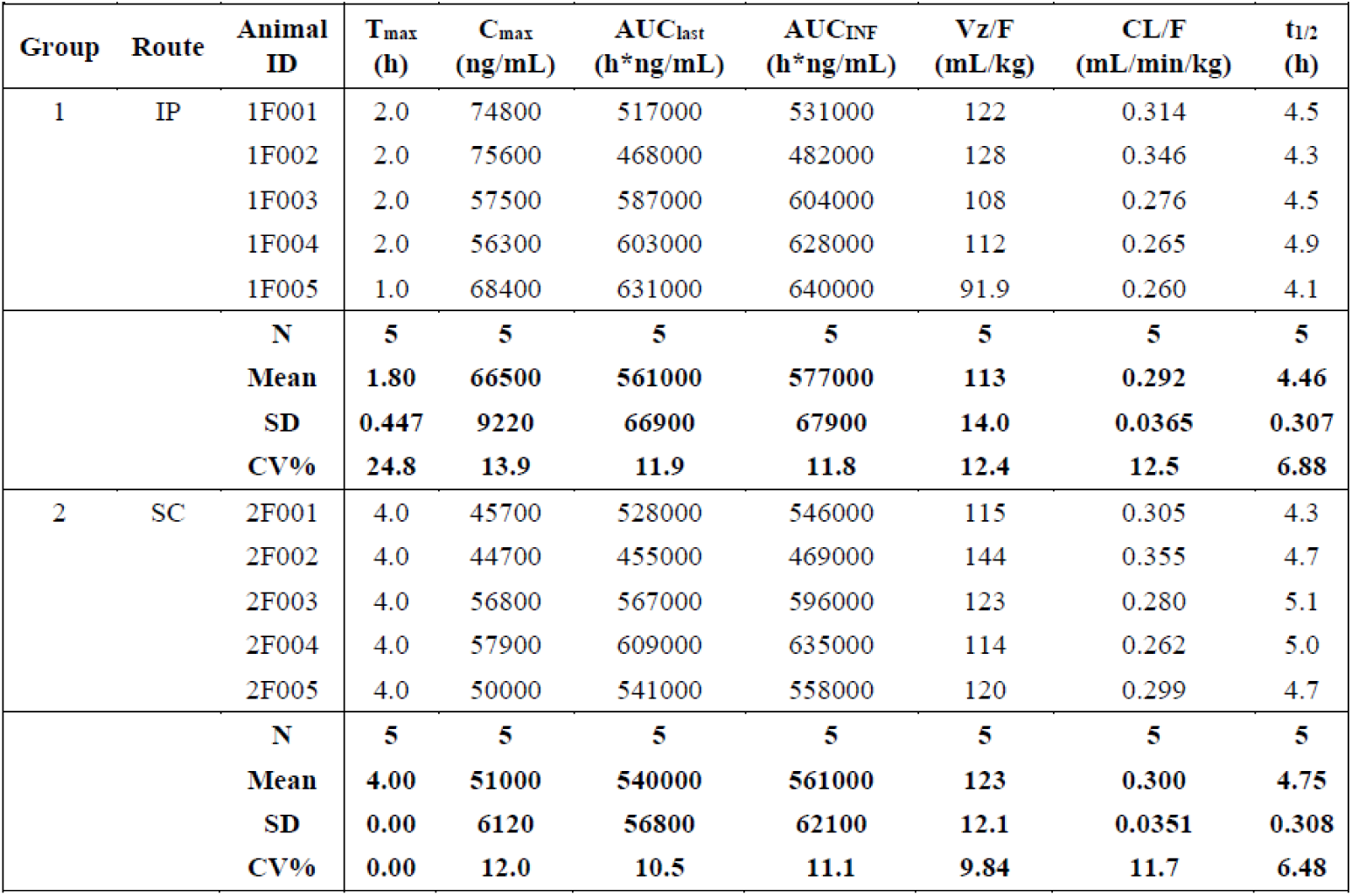
PK parameters and statistics for lassotide **47** dosed IP or SC at 10 mg/kg in 5 female Balb/c mice. Also mean concentration plot for IP and SC PK profiles are shown, indicating strong similarity with two routes of administration.

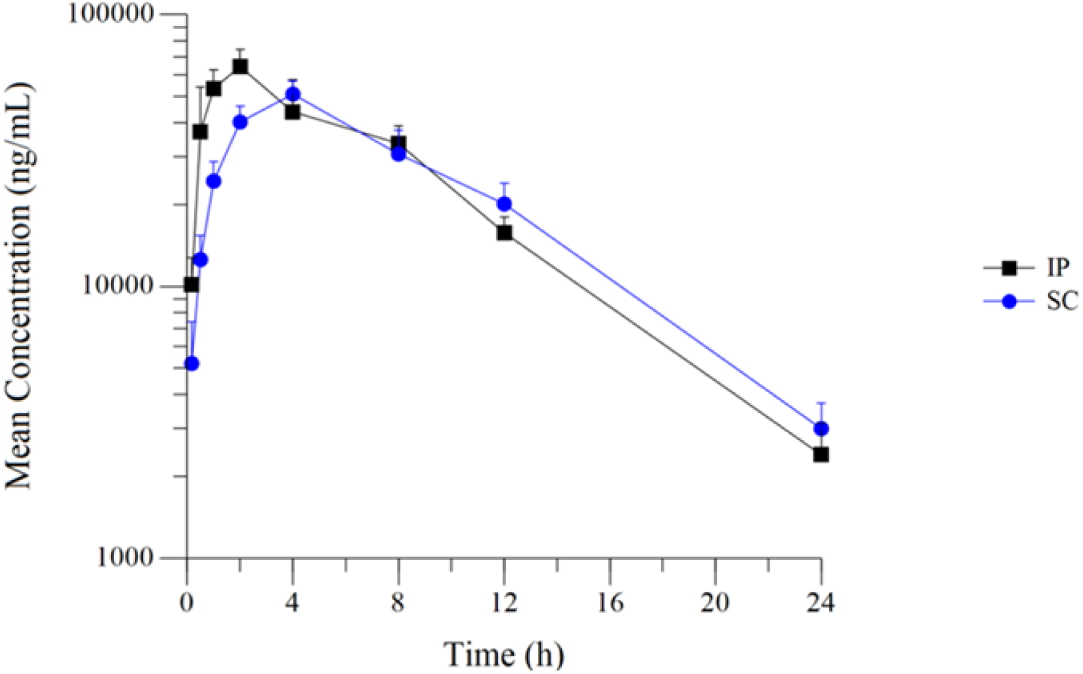

**Table S3E.**
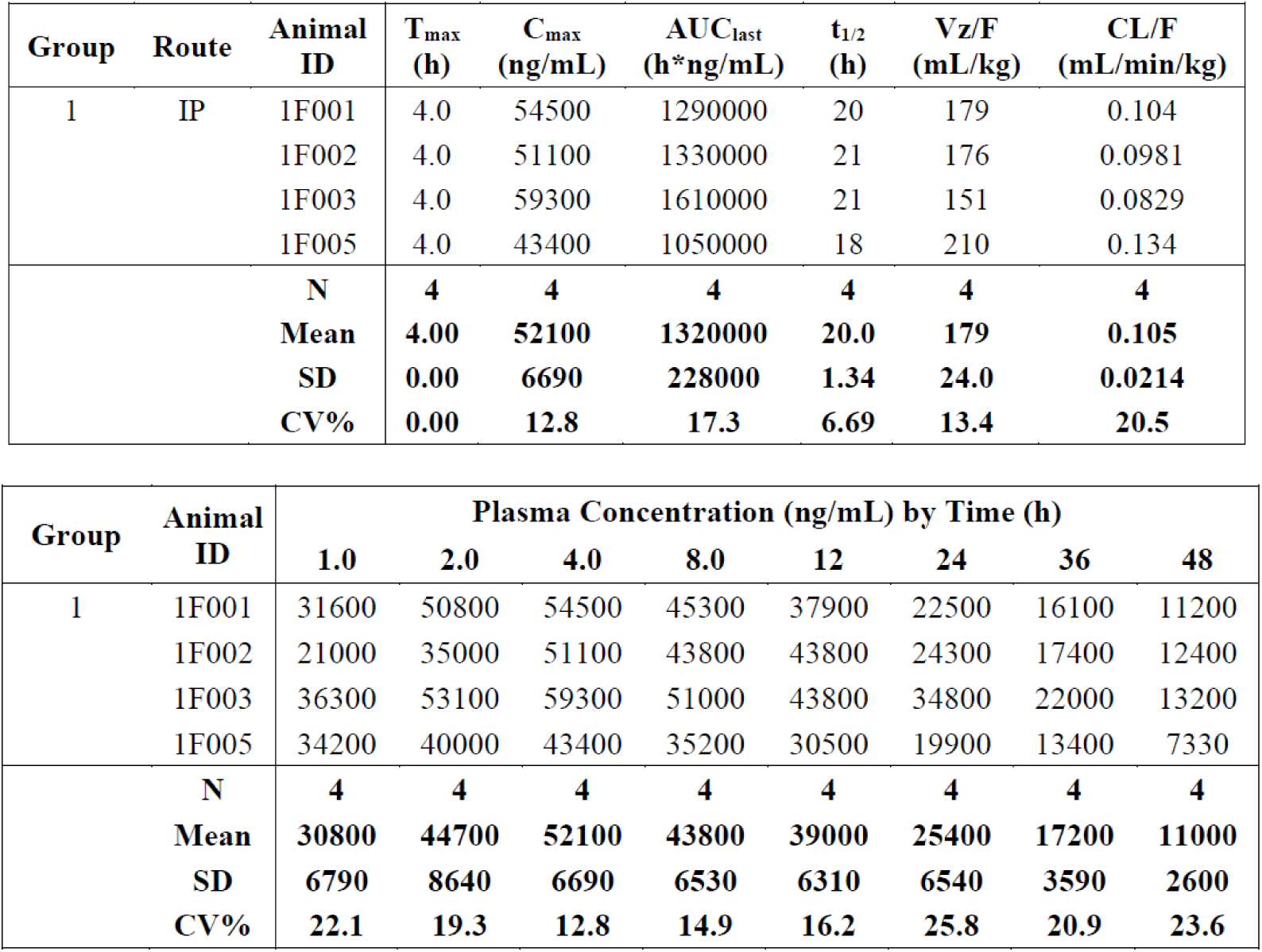
PK parameters and statistics for lassotide **52** dosed IP at 10 mg/kg in 4 female Balb/c mice. Also mean concentration plot for **52** is shown, indicating extended half-life.

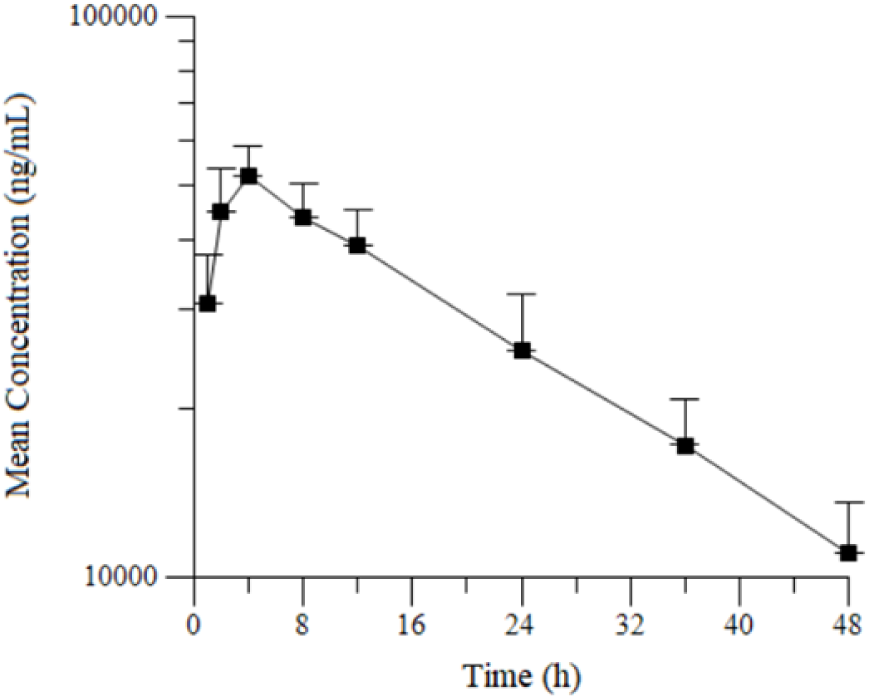

**Table S4.**
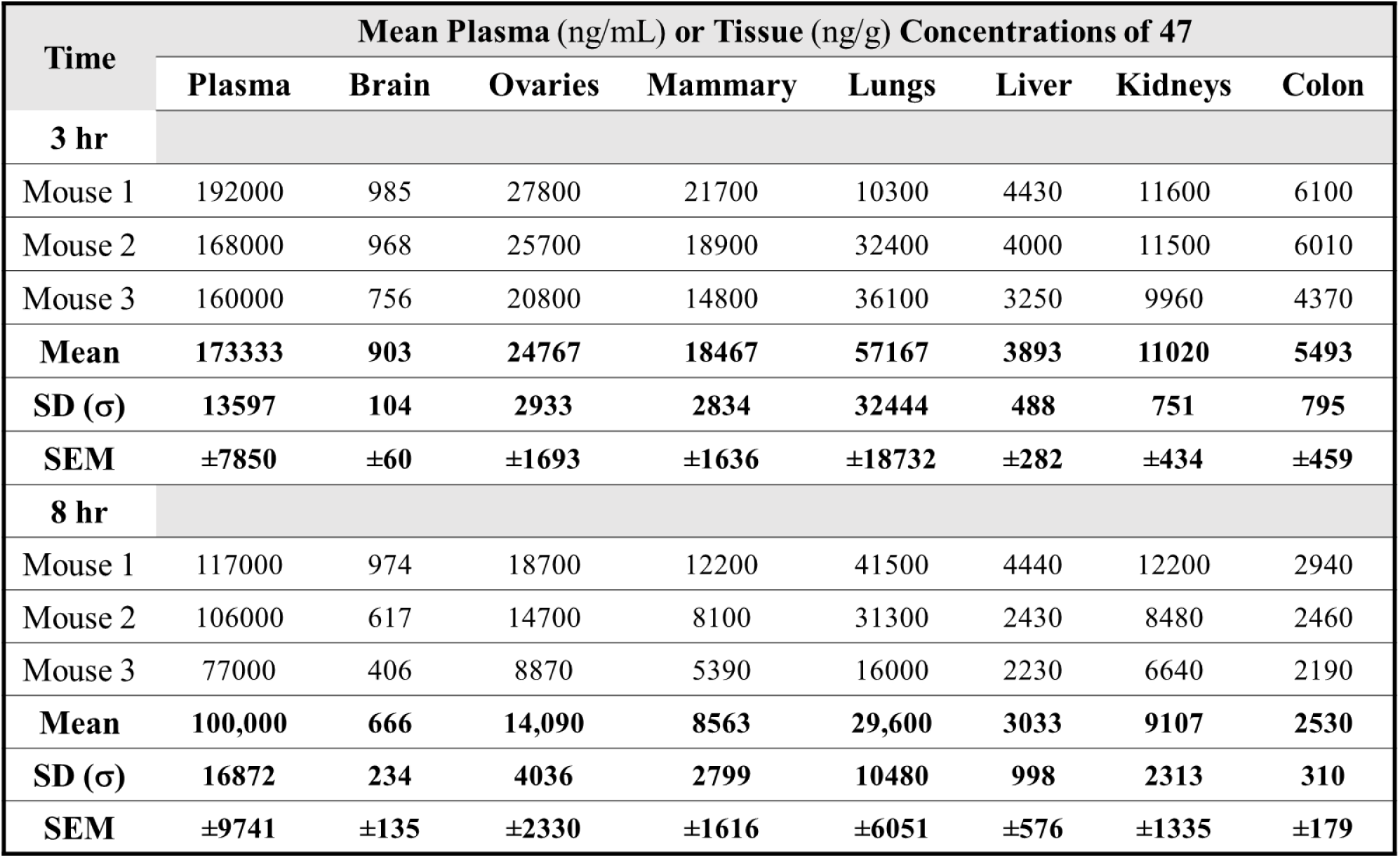
Tissue distribution study for lassotide **47** in six Balb/c mice. Tissues were collected and analyzed for lassotide concentrations at 3 h (n = 3) and 8 h (n = 3). Tissue concentration data for individual mice is shown in the Table below along with statistical analysis.

**Table S5.**
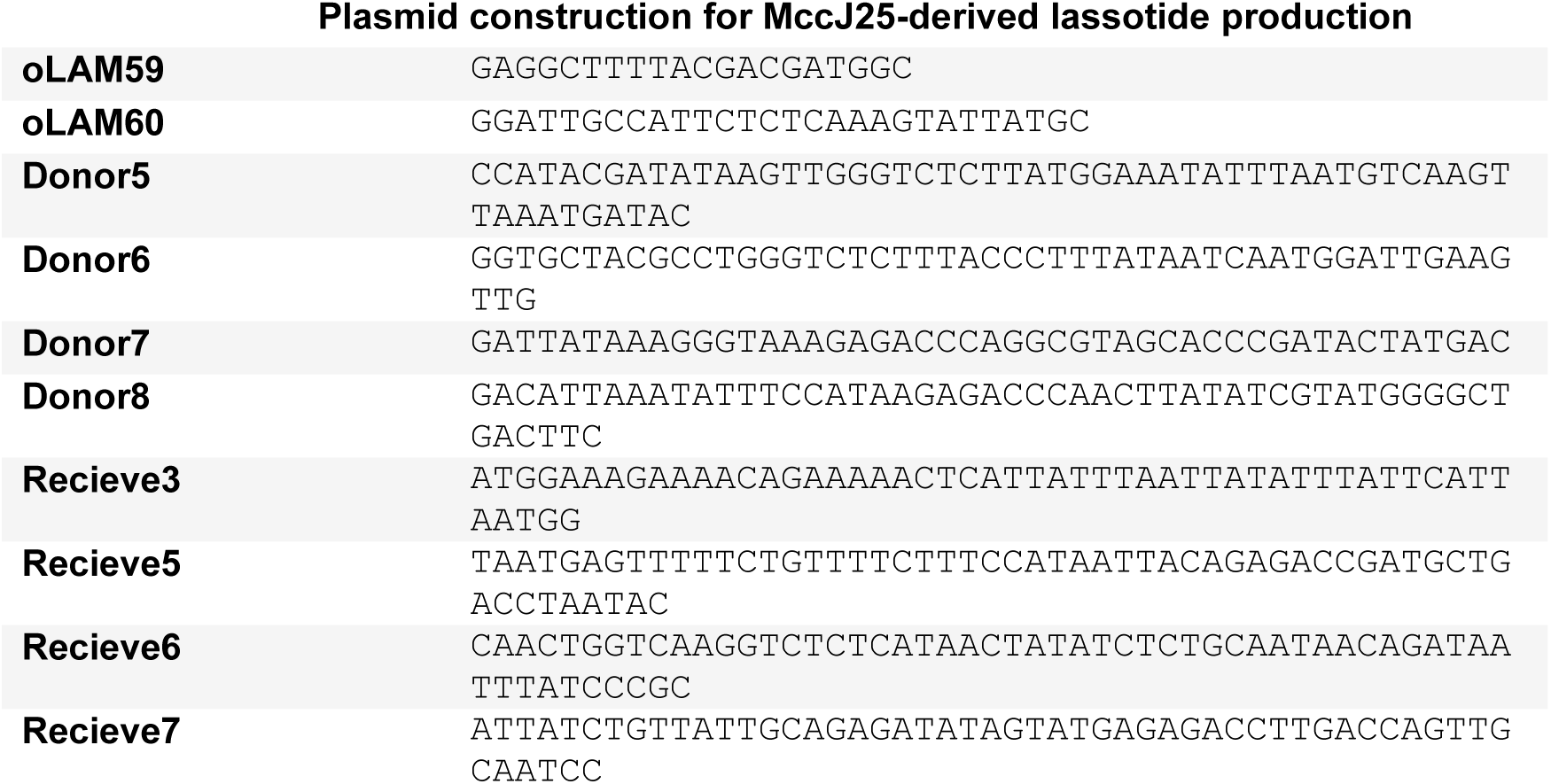
DNA sequences for plasmid construction used for MccJ25 variant production.

**Table S6.**
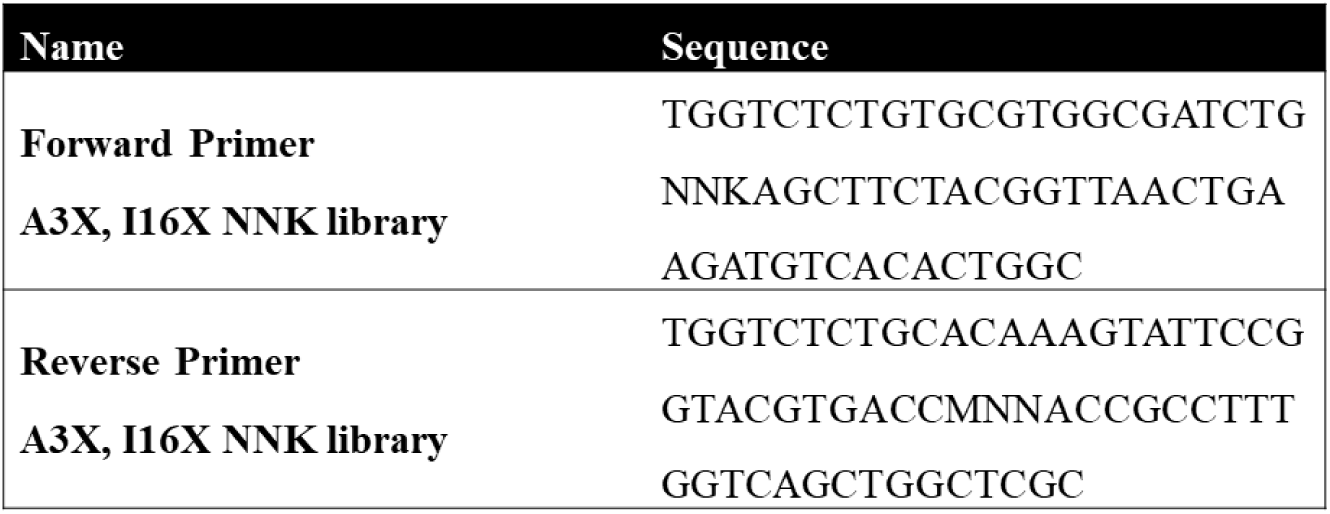
Primers used to generate 2-site NNK library at A3 and I16 positions of parent lassotide **14**.

### Supplementary Figures

**Figure S1.**
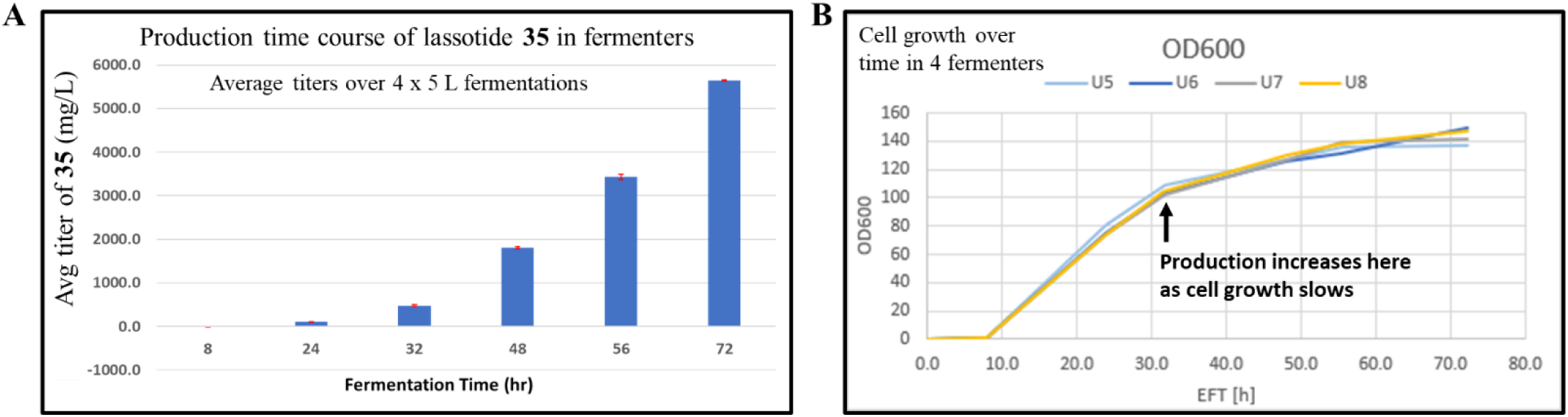
Production time course of lassotide **35** over 72 h in 4 x 5 L fermentations. **B**. Associated cell growth over time, as measured by OD600 through 72 h in 4 fermenters. Titers for tanks 1-4 (U5-U8) were 5.39, 5.64, 5.70, and 5.96 g/L, respectively. SEM was calculated as 0.10 g/L from these values.

**Figure S2.**
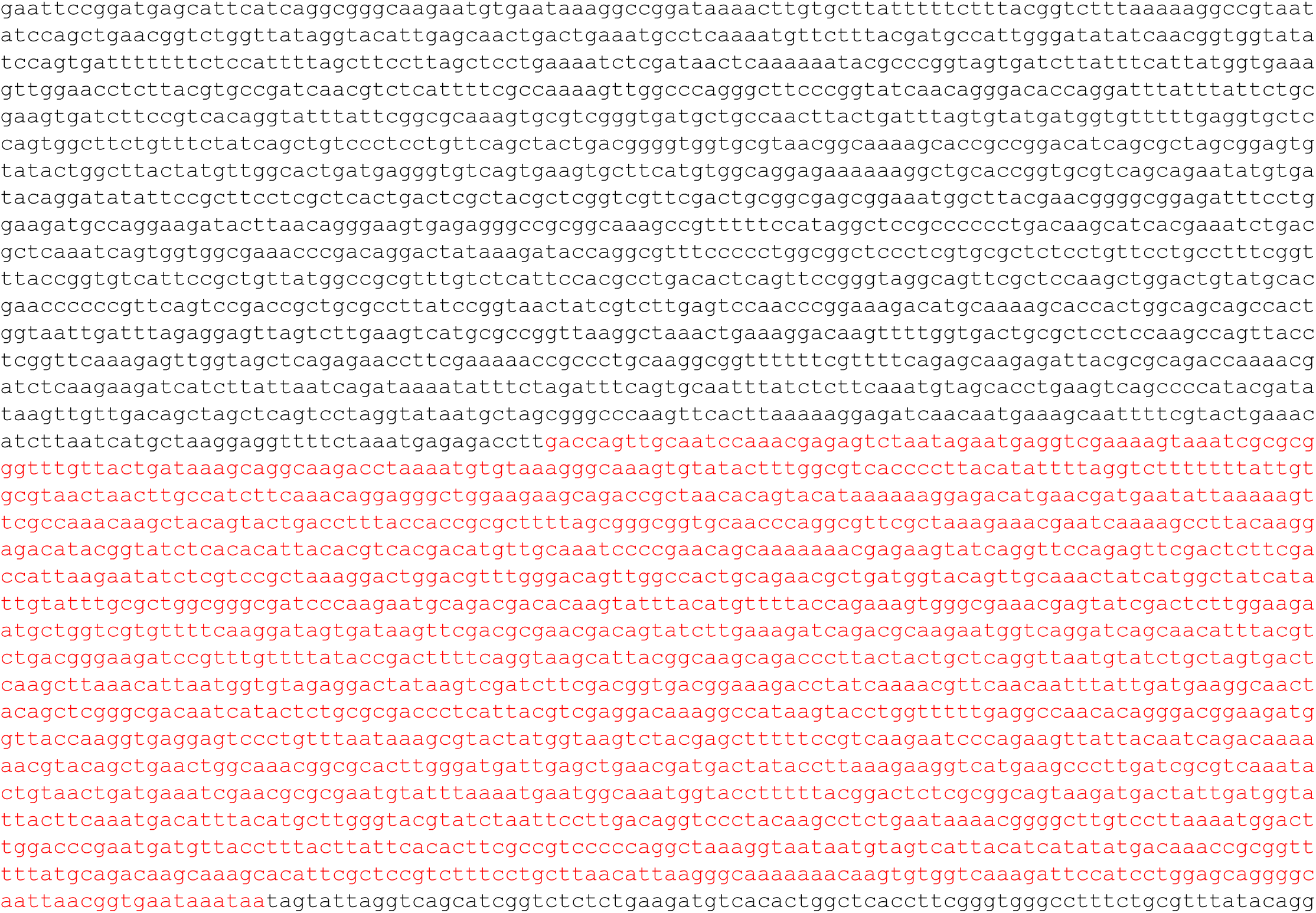
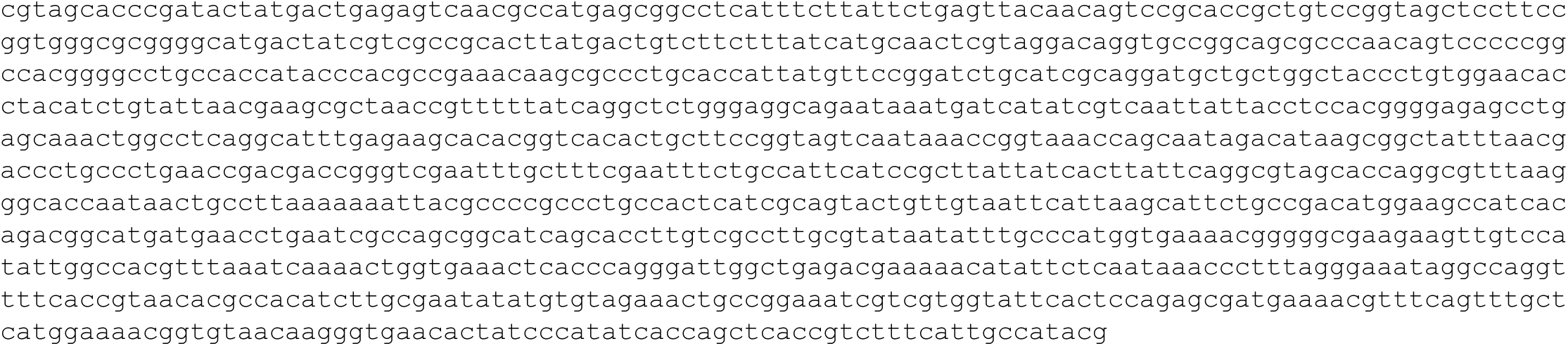
Nucleotide sequence of pLAM106, cloning plasmid for lasso peptide expression. *sacB* counterselection cassette is colored red.

**Figure S3.**
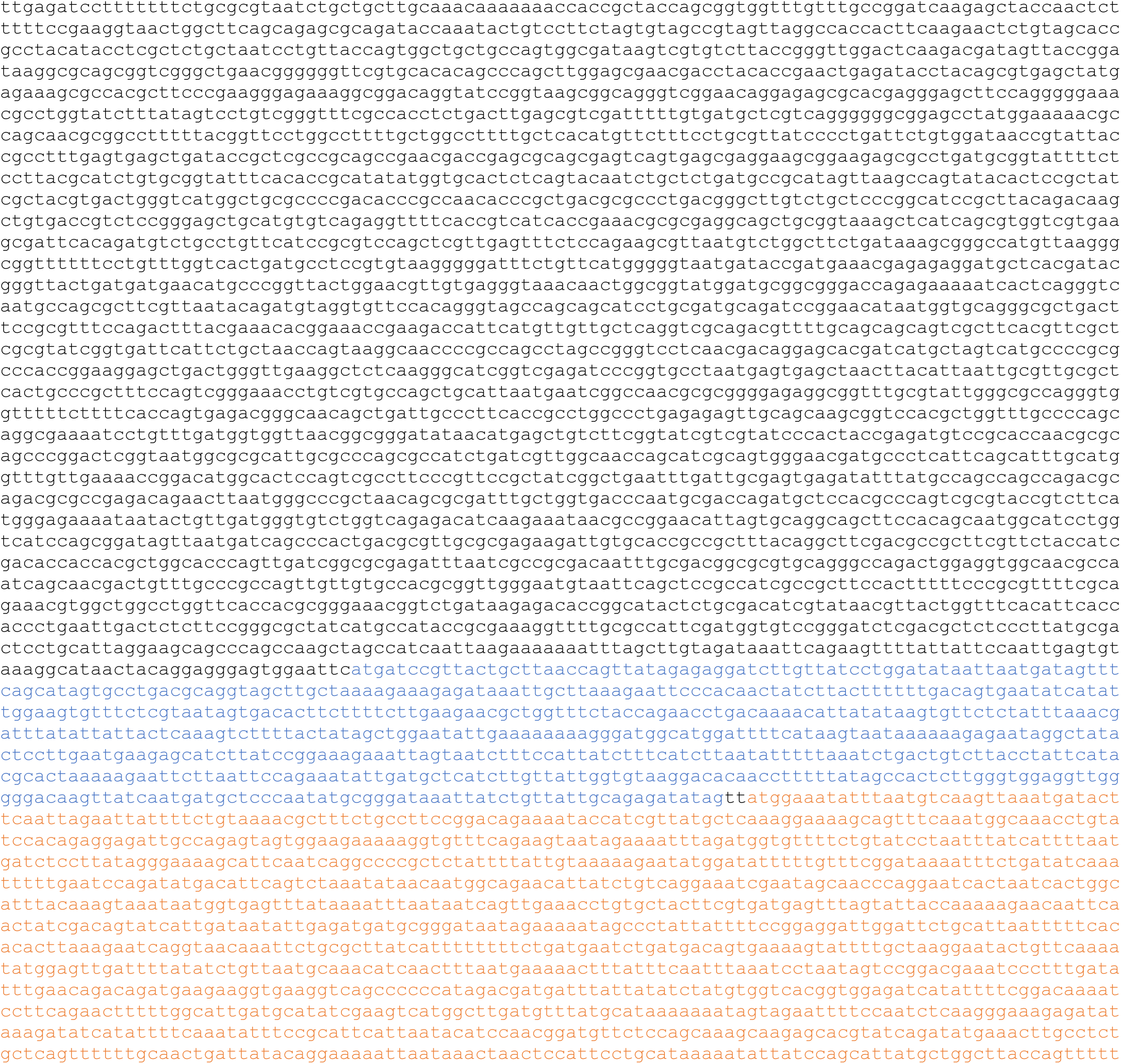
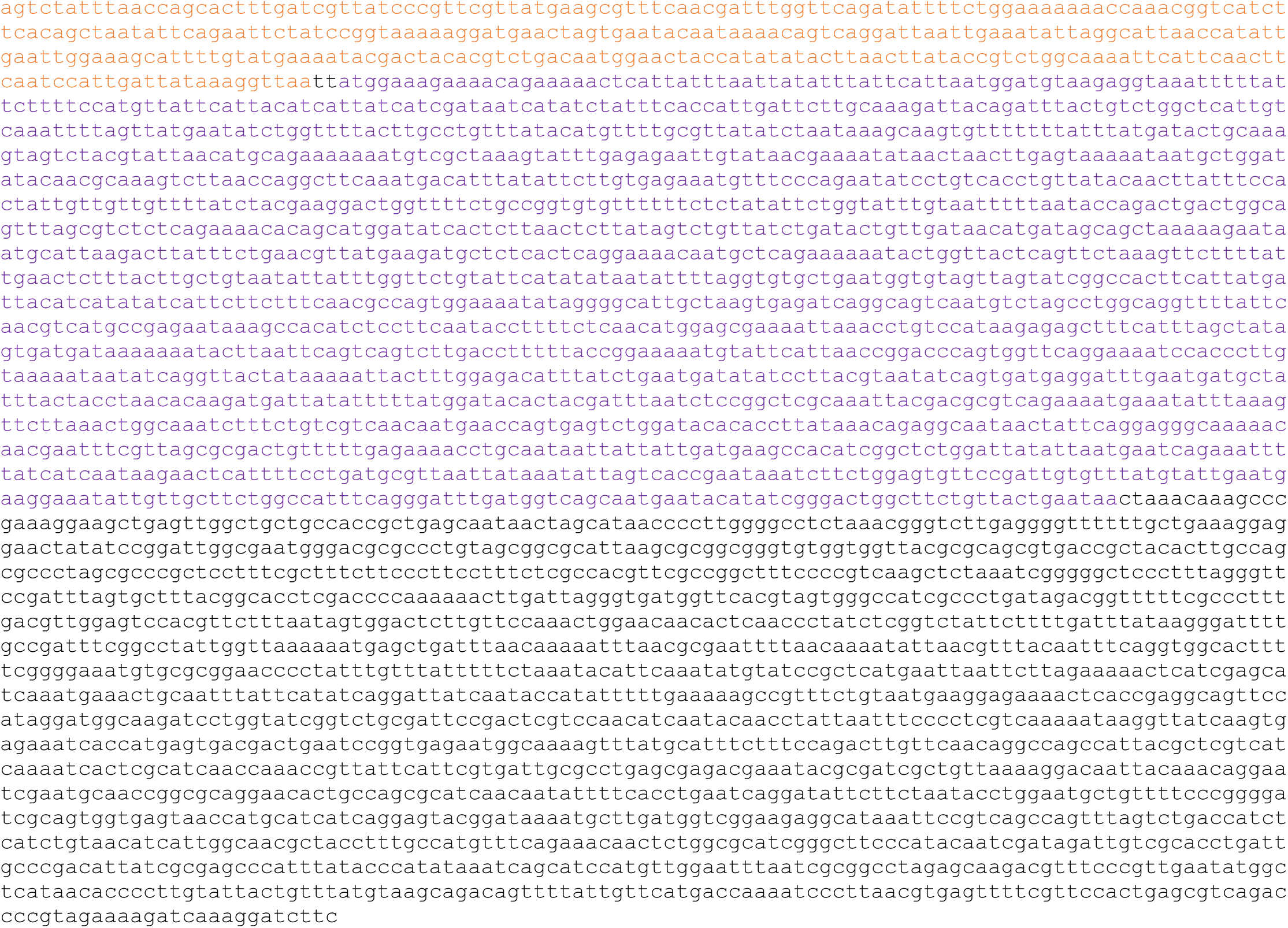
Nucleotide sequence of pLAM58 plasmid encoding the *mcjBCD* biosynthetic operon. The gene *mcjB* is colored blue, *mcjC* is orange and *mcjD* is purple.

**Figure S4.**
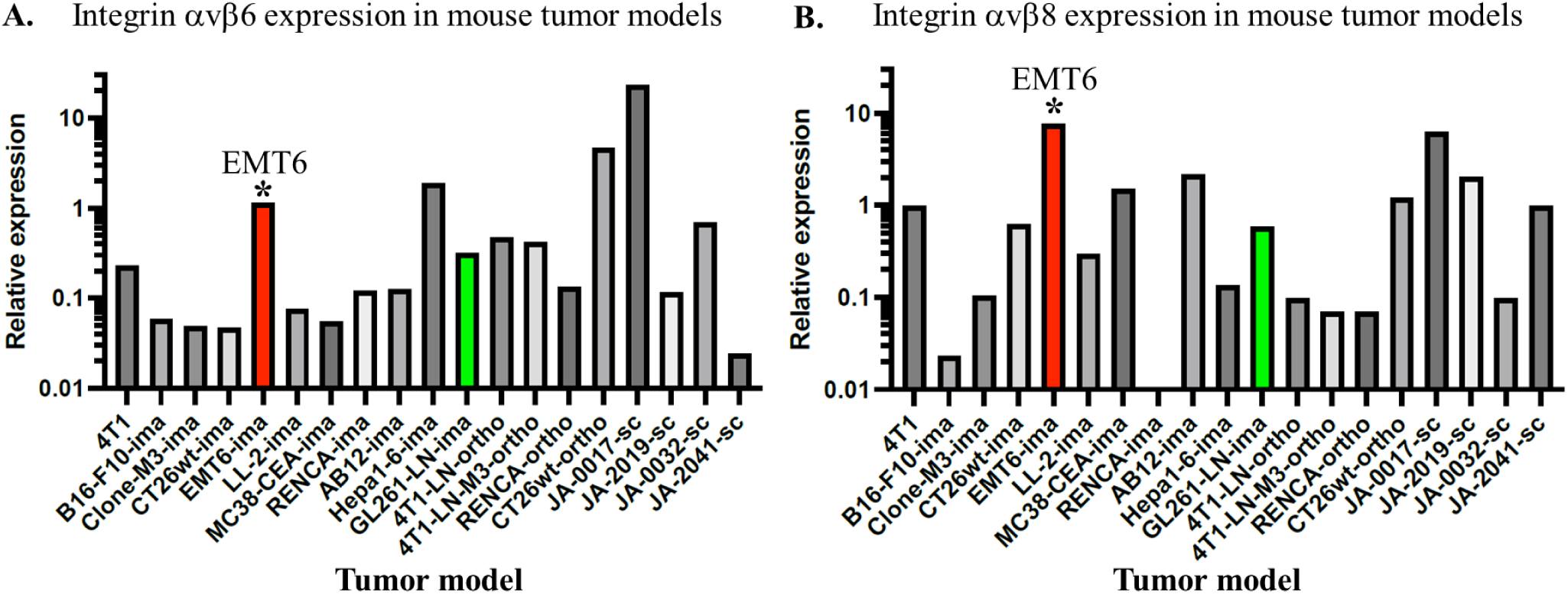
Reaction Biology expression data showing high expression levels of integrins αvβ6 and αvβ8 in ETM6 tumors.

**Figure S5.**
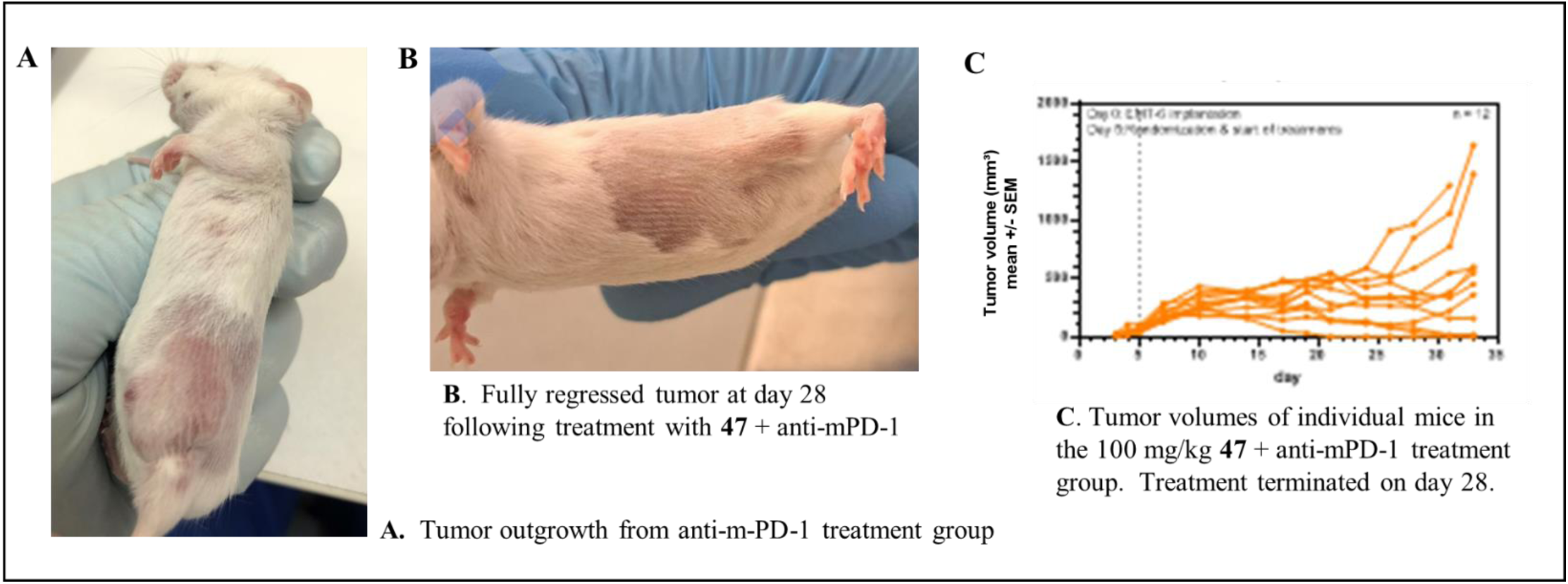
**A**. Photo showing outgrowth of an EMT6 tumor in the inguinal mammary fat pad of a mouse in the anti-mPD-1 treatment group. **B**. Photo showing complete tumor regression from **47** (100 mg/kg) + anti-mPD-1 treatment group. **C**. EMT6 tumor volumes of individual mice in the 100 mg/kg **47** + anti-mPD-1 treatment group over 33 days.

**Figure S6.**
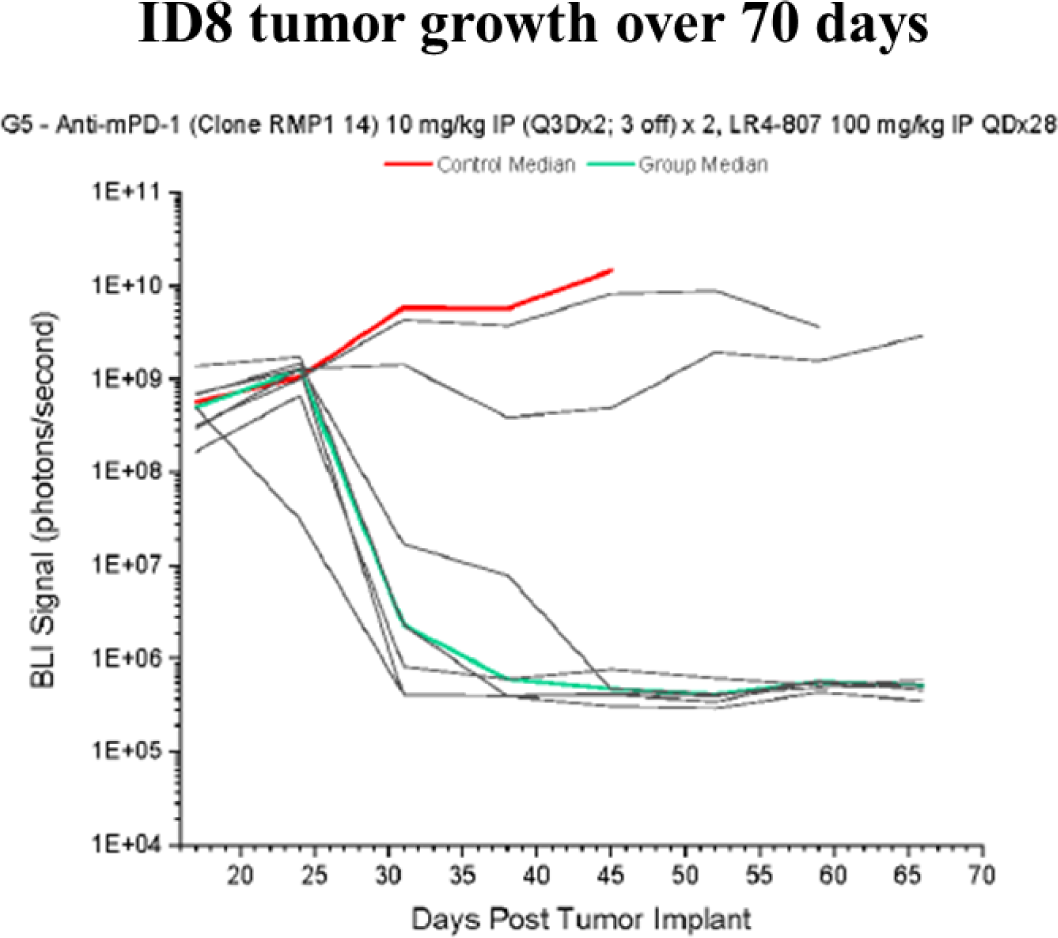
BLI measurements showing ID8 tumor growth during 28 days of treatment with **47**/anti-mPD-1 and 25 days after treatment termination. No tumor growth was observed after dosing stopped, indicating a durable response to treatment.

**Figure S7.**
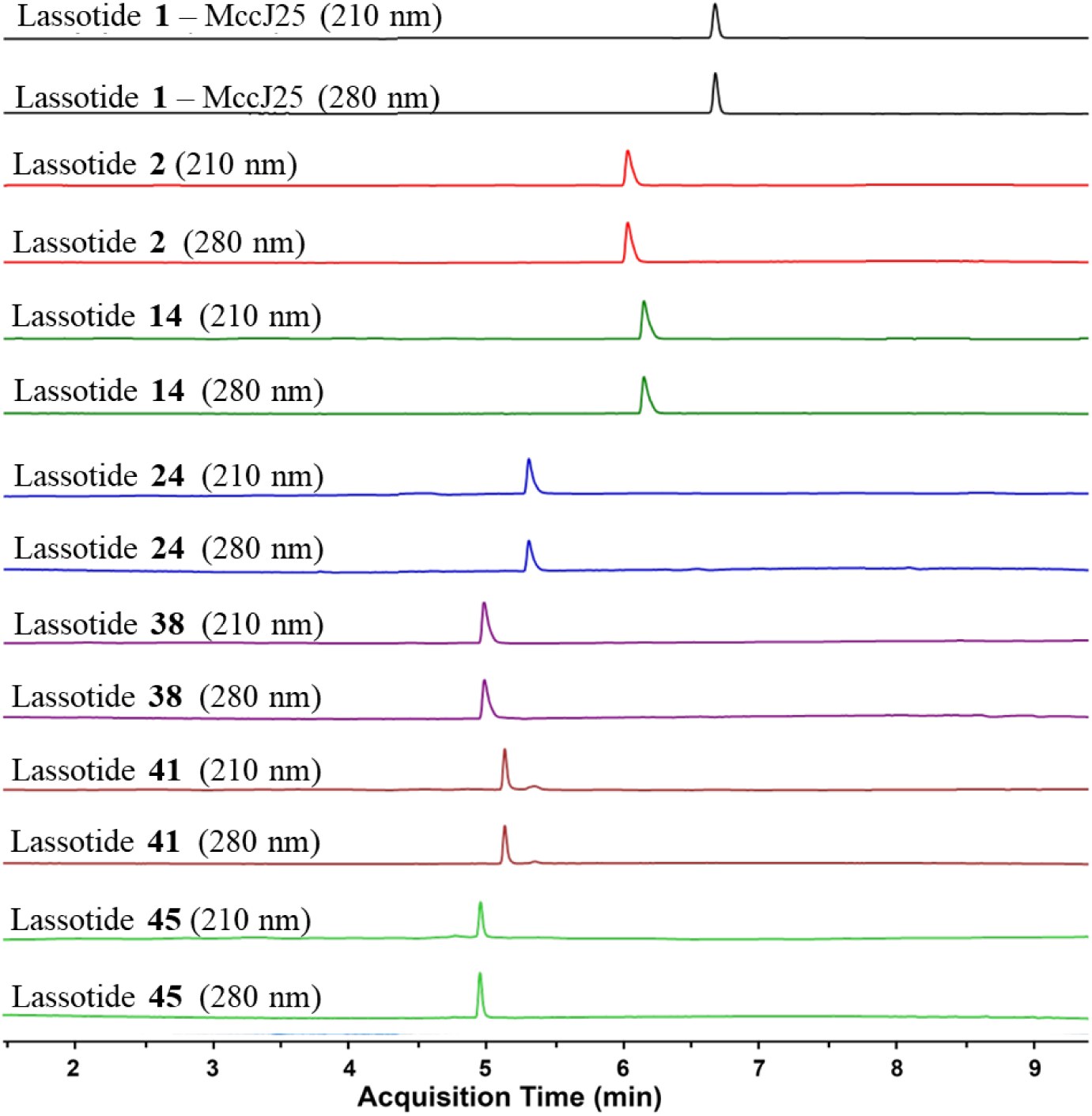
HPLC traces for representative lassotides following semi-preparative purification, as described in the Experimental Methods section. Semi-preparative HPLC afforded all lassotides with purity levels ≥ 97%.

**Figure S8.**
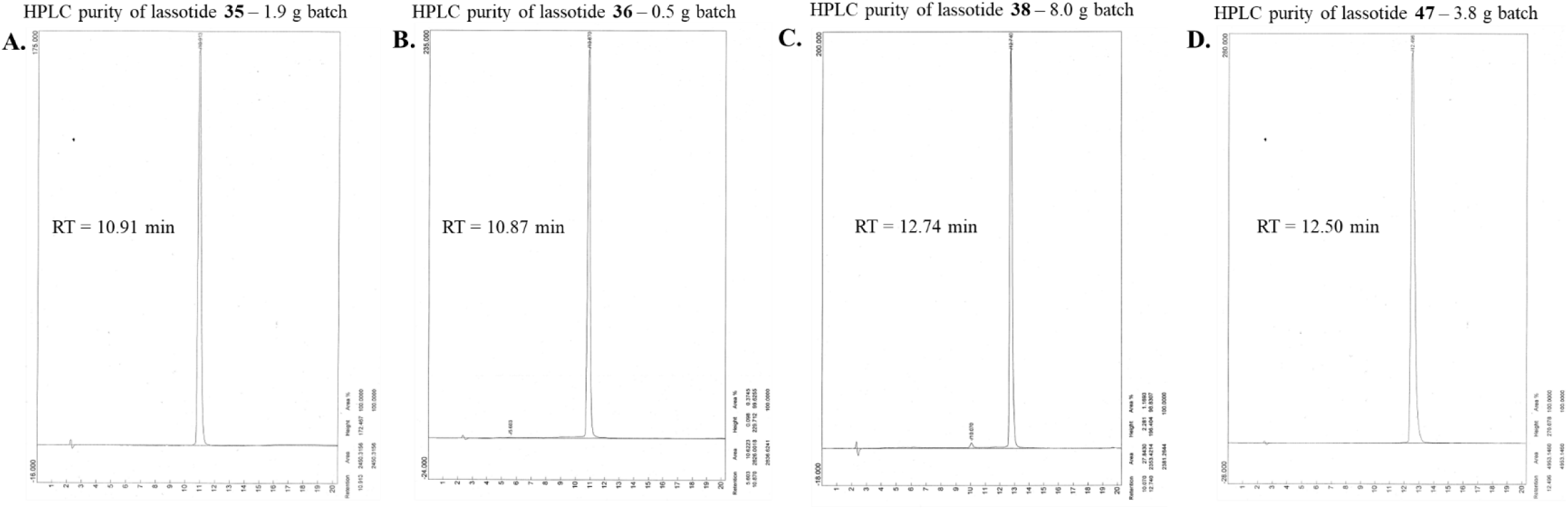
Analytical QC check for lassotides purified by preparative HPLC. Purified lassotides were analyzed using a Shimadzu LC-10ADvp HPLC equipped with a Vydac C18 25 cm x 4.6 mm, 5-micron column. Mobile phase was 0.1% TFA in 15-20% CH3CN increased to 35-50% CH3CN over 20 min. **A**. Lassotide **35**, **B**. Lassotide **36**, **C**. Lassotide **38**, **D**. Lassotide **47**. In all cases, lassotide purity was ≥ 98% in all cases.

**Figure S9A.**
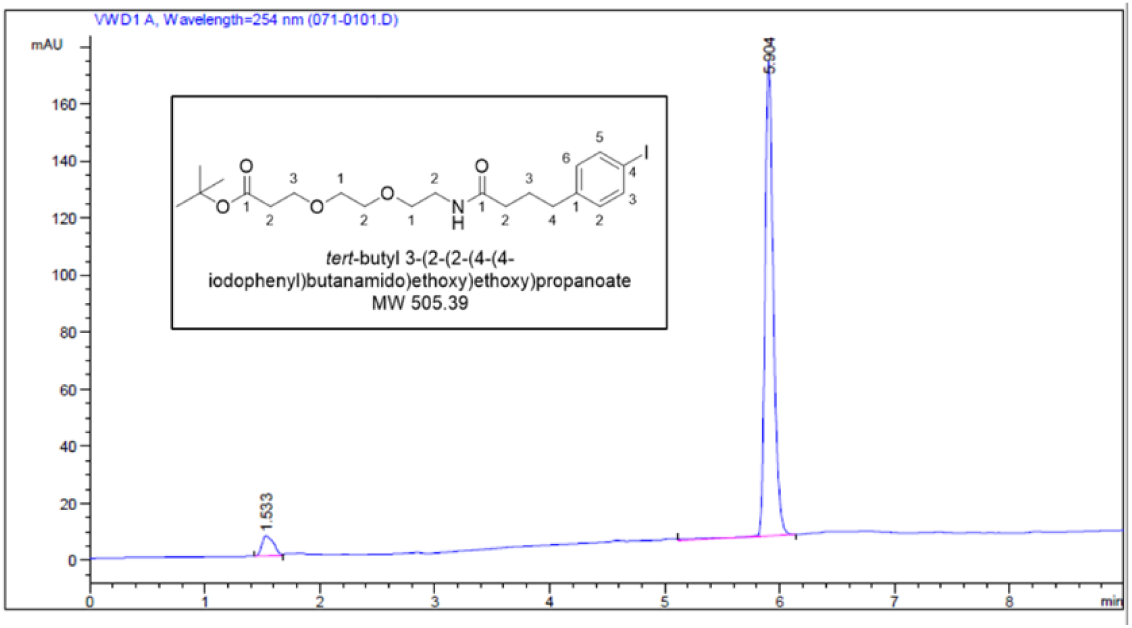
HPLC data for chromatographically purified *tert*-butyl ester of Acyl **50**.

**Figure S9B.**
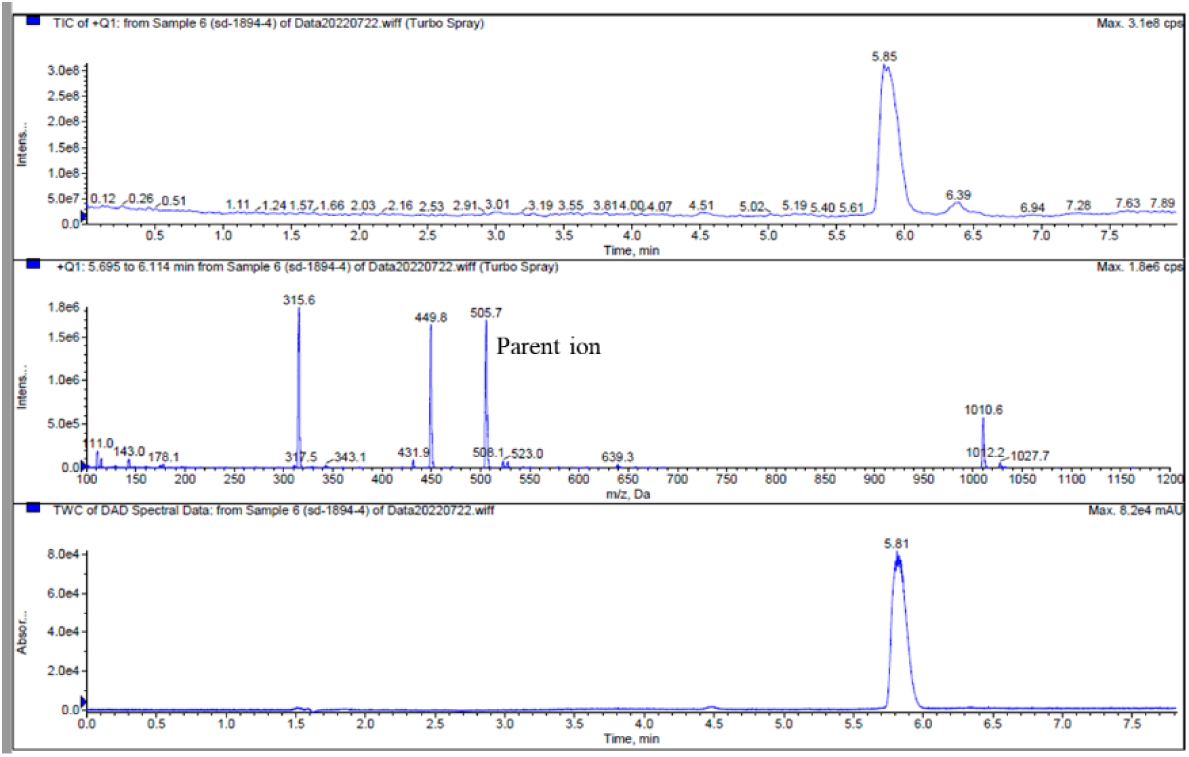
LCMS data for *tert*-butyl ester of Acyl **50** showing expected parent ion *m/z* = 505.7.

**Figure S9C.**
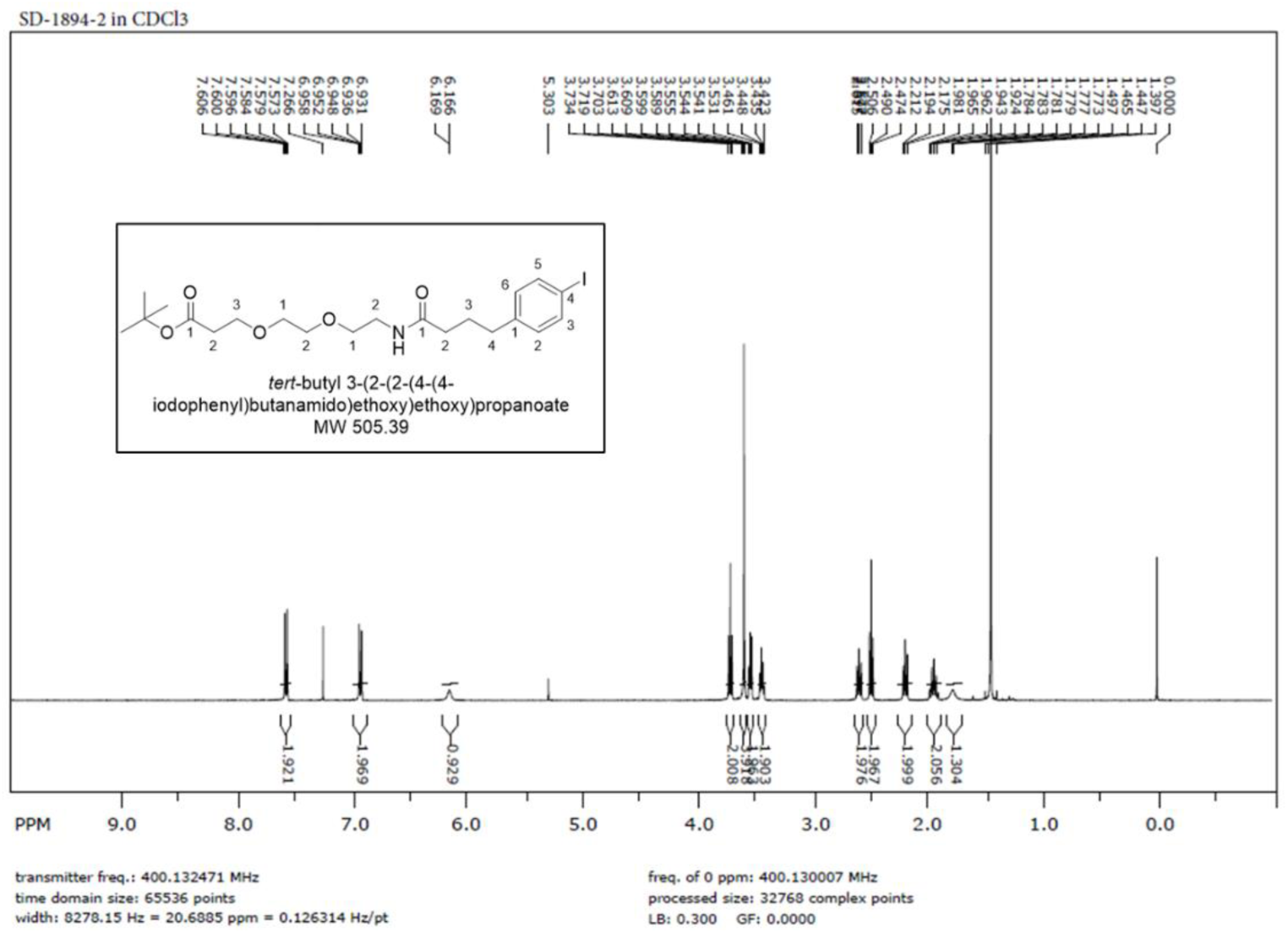
^1^H NMR spectrum of *tert*-butyl ester of Acyl **50** showing correct peaks and integration for structure shown.

**Figure S10A.**
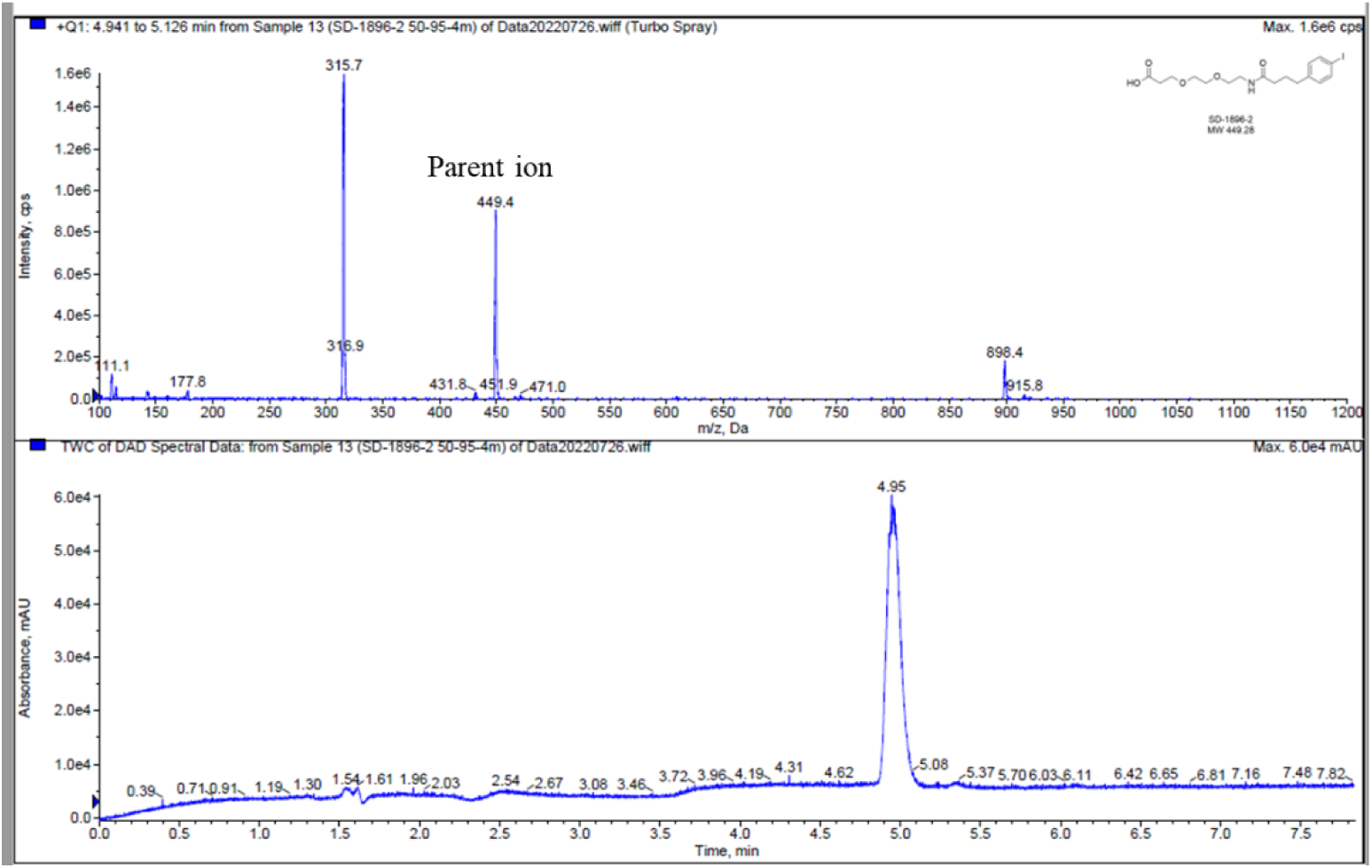
LCMS data for Acyl **50** showing expected parent ion *m/z* = 449.4.

**Figure S10B.**
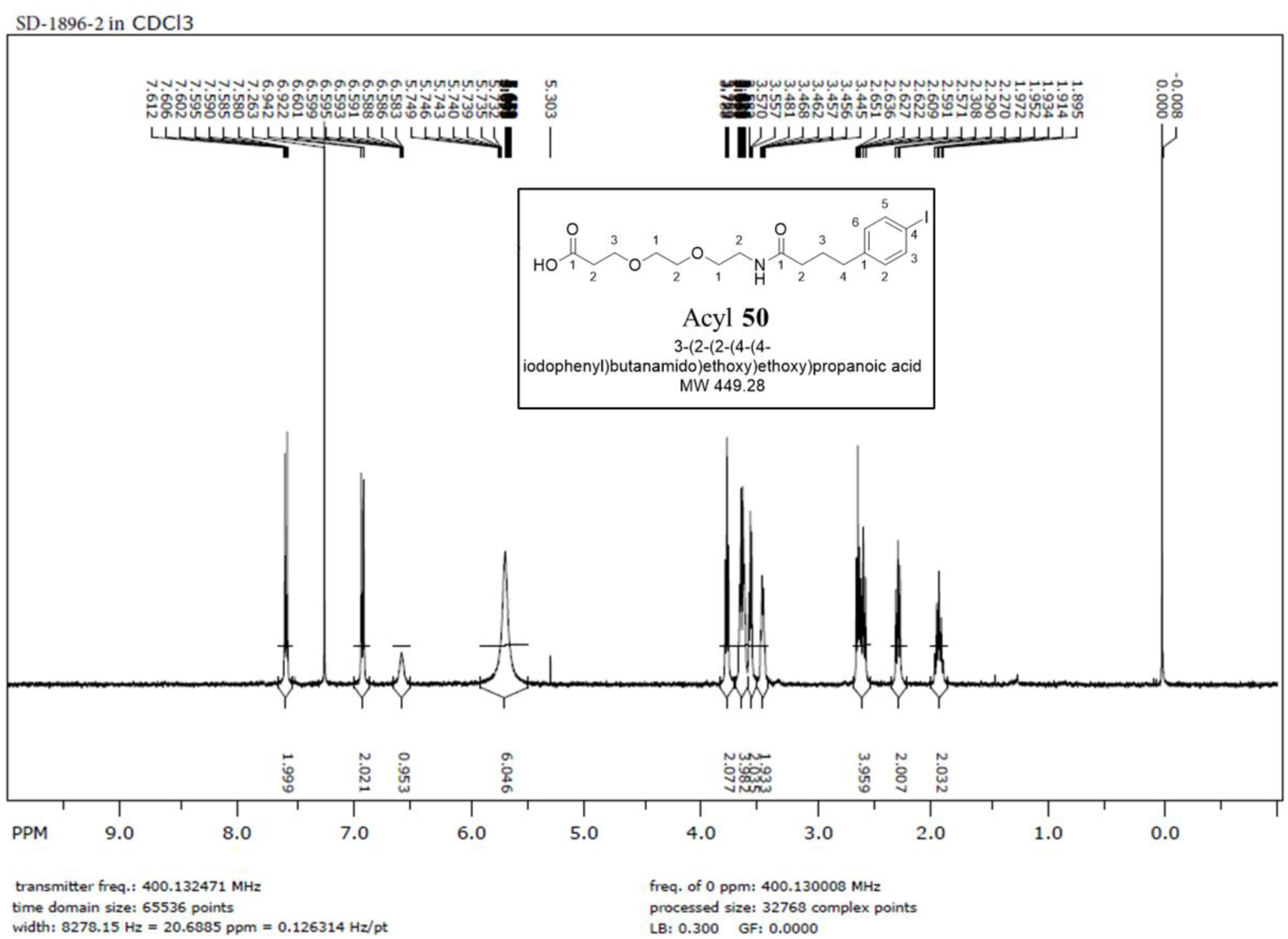
^1^H NMR spectrum of Acyl **50** showing correct peaks and integration for structure shown.

## Notes

### Competing Interest Statement

The authors have declared no competing interest.

